# Colloidal physics modeling reveals how per-ribosome productivity increases with growth rate in *E. coli*

**DOI:** 10.1101/2021.10.27.466129

**Authors:** Akshay J. Maheshwari, Alp M. Sunol, Emma Gonzalez, Drew Endy, Roseanna N. Zia

## Abstract

Faster growing cells must synthesize proteins more quickly. Increased ribosome abundance only partly accounts for increases in total protein synthesis rates. The productivity of individual ribosomes must increase too, almost doubling by an unknown mechanism. Prior models point to diffusive transport as a limiting factor but surface a paradox: faster growing cells are more crowded, yet crowding slows diffusion. We suspected physical crowding, transport, and stoichiometry, considered together, might reveal a more nuanced explanation. To investigate, we built a first-principles physics-based model of *E. coli* cytoplasm in which Brownian motion and diffusion arise directly from physical interactions between individual molecules of finite size, density, and physiological abundance. Using our microscopically-detailed model, we predict that physical transport of individual ternary complexes accounts for ~80% of translation elongation latency. We also find that volumetric crowding increases at faster growth even as cytoplasmic mass density remains relatively constant. Despite slowed diffusion, we predict that improved proximity between ternary complexes and ribosomes wins out, illustrating a simple physics-based mechanism for how individual elongating ribosomes become more productive. We speculate how crowding imposes a physical limit on growth rate and undergirds cellular behavior more broadly. Unfitted colloidal-scale modeling offers systems biology a complementary “physics engine” for exploring how cellular-scale behaviors arise from physical transport and reactions among individual molecules.

## Introduction

Protein synthesis is essential for cell maintenance and reproduction. For example, *Escherichia coli* (*E. coli*) cells synthesize the majority of their dry mass as protein every cell doubling. Accordingly, cells that grow more quickly must produce proteins more quickly. In quantitative detail, as *E. coli* growth speeds up five-fold protein synthesis across the entire cell increases 15-fold (Dennis & Bremer, 2008). Meanwhile, for the same growth rate increase, the quantity of ribosomes increases only nine-fold (**Figure S1**), suggesting that the absolute productivity of individual ribosomes must also somehow increase – almost doubling as growth quickens (**Figure 1**) (Bremer and Dennis, 1996; Dalbow and Young, 1975; Dennis and Bremer, 2008; Forchhammer and Lindahl, 1971; Klumpp et al., 2013; Pedersen, 1984; Young and Bremer, 1976). While it is easy to understand why having more translation machinery increases total protein synthesis capacity, it is not obvious how faster growing cells achieve the translation elongation rates needed to sustain growth.

**Figure 1.**
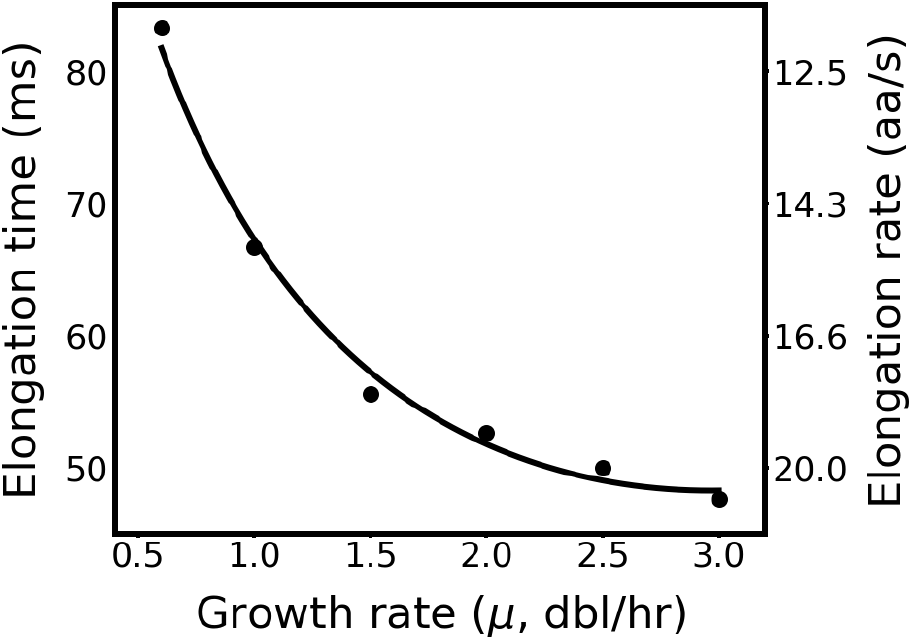
Individual ribosomes make proteins more quickly as growth quickens. Total latency per peptide bond (left y-axis) or elongation rate (right y-axis) versus growth rate (x-axis). Experimental data from Bremer and Dennis, 1996; Dalbow and Young, 1975; Dennis and Bremer, 2008; Forchhammer and Lindahl, 1971; Klumpp et al., 2013; Pedersen, 1984; and Young and Bremer, 1976. Solid line is a second-order polynomial fit of experimental elongation rates.

Bremer & Dennis hypothesized that individual ribosome activity speeds up at faster growth rates owing to increased tRNA charging and also due to shifts in codon distribution among mRNA (Bremer and Dennis, 1996). However, subsequent work has shown that overall tRNA charging remains relatively constant across growth rate indicating that other mechanisms are likely at play (Avcilar-Kucukgoze et al., 2016). Another possibility is that the intrinsic chemical kinetics of peptide bond formation by the ribosome accelerate with increasing growth rates. For example, in exploring how to adapt chemical kinetic rates obtained from *in vitro* experiments for use with *in vivo* models, Rodnina and co-workers fit parameter values to data and showed that faster chemical kinetic rate constants could account for increased rates of peptide bond formation (Rudorf et al., 2014). However, the specific molecular mechanisms that might account for such parameter changes are unknown. As a third possibility, Hwa and co-workers hypothesized that physical processes could play a limiting role in determining the elongation rate of individual ribosomes (Klumpp et al., 2013). More specifically, by accounting for Brownian diffusion of ternary complexes via a growth-rate independent diffusion constant within a Michaelis-Menten kinetics-based model of translation elongation, Hwa and co-workers inferred that physical changes in cytoplasm could lead to changes in growth rate. Taken together, such studies suggest that both chemistry and physics likely play a role in the speedup of translation elongation.

However, understanding any potential speedup mechanism is challenging exactly because the chemistry and physics of translation elongation are complex and coupled. For example, the biochemical processes required are combinatorial: matching must take place between 42 unique ternary complexes and 64 possible triplet codons. Accordingly, any particular elongating ribosome may encounter numerous mismatching ternary complexes prior to a successful matching reaction. As a second example, the length- and time-scales of underlying processes span three and nine orders of magnitude, respectively; specifically, ternary complexes and ribosomes interact with surrounding biomolecules and each other over nanometers and nanoseconds but execute processes over microns and seconds. As a third example, while higher concentrations of ternary complexes might be expected to increase the frequency of encounters with ribosomes, the resulting increase in crowding might slow the physical search process. Such complexities are compounded by the fact that everything is happening in parallel among hundreds of thousands of self-mixing molecules in a growth-rate dependent and crowded cytoplasm.

We address these challenges by modeling both the physics and chemistry of translation elongation in a combined framework. To do so, we adapted an open-source simulation tool (Andrews et al., 2010) to more accurately represent transport and interactions among molecules comprising self-mixing systems. In our framework we explicitly represent the transport dynamics of individual biomolecules as they physically interact and chemically react, with nanometer and nanosecond resolution, to simulate processes spanning minutes in time. A key aspect of our approach is the robust modeling of Brownian motion and colloidal-scale particle interactions such that these molecules undergo the inertialess physical encounters appropriate to the colloidal regime (Ermak and McCammon, 1977; Heyes and Melrose, 1993; Zia, 2018). In particular, we measure diffusion rates for individual molecules, which is influenced directly by their mass, size, and crowding such that resulting emergent behaviors (such as transport and search time) are not merely *ex post facto* fits to expected results. When combined with a well-known multi-step kinetic model for the reactions leading to peptide bond formation (Kothe et al., 2004) our framework enables analysis of the combined physical and chemical dynamics underlying translation elongation.

We employed our framework to explore how protein synthesis rates in *E. coli* should be expected to change with growth rate, directly representing the growth-rate dependent cytoplasm via first-principles modeling of physical and chemical dynamics without parameter fitting. Starting from well-established measurements of macromolecular composition and physical properties of *E. coli* cytoplasm at varying growth rates, we demonstrate how well-known changes in the composition of cytoplasm are entirely sufficient to account for the speedup of translation elongation by individual ribosomes. We also identify the detailed contributions of transport and reaction to total elongation latency by monitoring the trajectories of and reactions between ternary complexes and ribosomes in simulation, finding that transport is the dominant component defining elongation latency. We find that physiological cytoplasmic crowding speeds up the transport mechanism and thus elongation rates overall. We confirm that the expected speedup due to crowding is insensitive to changes in chemical kinetics needed to exactly match observed elongation rates. Finally, we explore how still-greater crowding, beyond naturally observed limits, should lead to a collapse of the colloidal-scale transport speedup mechanism that ultimately limits the performance of self-mixing living systems.

## Results

### I. Embedding chemical kinetics within physical transport

We constructed a spatially resolved chemical and physical framework to model the combined roles of reaction chemistry and transport physics in translation elongation (**Figure 2A, 2B**), tracking the time spent by ternary complexes unbound and in motion (transport latency, *τ*_transport_) as well as reacting with mismatching or matching ribosomes (reaction latency, *τ*_rxn_) until a matching reaction successfully completes. Together, transport latency and reaction latency make up elongation latency (*τ*_elong_).

**Figure 2.**
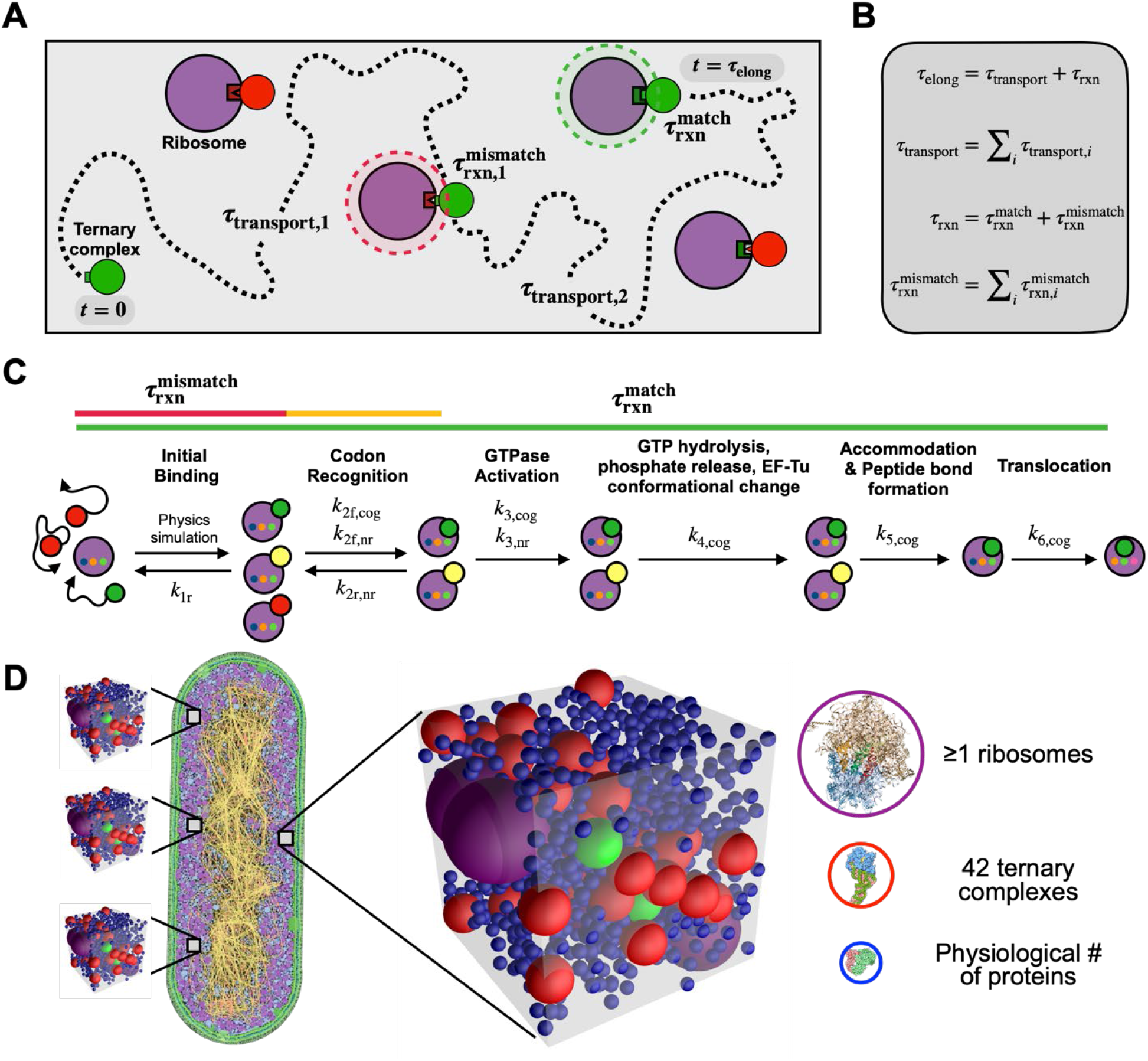
The physical context for translation elongation can be formalized. **(A)** Schematic of physical and chemical processes that contribute to translation elongation latency. Multiple transport and reaction steps (dashed line) may occur before a ternary complex (green/red) encounters and reacts with an unoccupied, matching ribosome (purple). The time ternary complexes spend unbound while searching for ribosomes is defined as transport latency (*τ*_transport_) and the time ternary complexes spend bound in either mismatching (red shaded circle) or matching reactions (green shaded circle) is defined as reaction latency (*τ*_rxn_). The time the entire process takes is defined as elongation latency (*τ*_elong_). **(B)** Mathematical definitions of translation elongation latencies. Elongation latency (*τ*_elong_) is the sum of transport latency (*τ*_transport_) and reaction latency (*τ*_rxn_), the latter of which is the sum of both mismatching 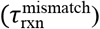 and matching 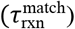 reaction latencies. **(C)** Schematic of the kinetic mechanism of translation elongation within ribosomes (purple). Ternary complexes are either cognate (green), near-cognate (yellow), or non-cognate (red) to any particular ribosome, which determines kinetic rates. Mismatching reaction latency results from reversible reactions with non-cognate and near-cognate ternary complexes (red and yellow lines), while matching reaction latency results from cognate ternary complexes proceeding through the full kinetic process (green line). **(D)** Translation elongation is evaluated by constructing ensembles of statistically representative “translation voxels” that, in their minimal form, contain exactly 42 ternary complexes (cognate: green, non-cognate: red), at least one ribosome (purple), and numerous average-sized proteins representing all other surrounding proteins (blue). Depiction of *E. coli* adapted with permission from Goodsell, 2009; molecular abundances adapted from literature (Main text and Methods).

To estimate reaction latencies we represented the molecular reactions between ribosomes and ternary complexes following the individual chemical steps of protein synthesis, accounting for differences due to reactions involving cognate, near-cognate, and non-cognate ternary complexes (**Figure 2C**). We used well-established *in vitro* kinetic measurements to parameterize our model (**Table S5**) and developed physiologically accurate distributions of expected reaction latencies via a Markov-process (**Methods**). We analyzed the resulting reaction latency distributions, finding that matching reactions between cognate ternary complexes and ribosomes take 42 ms on average when successful (68% probability) (**Figure S2**). We also found that cognate ternary complexes can be rejected (32% probability), in which case reactions take 1.4 ms on average. Mismatching reactions involving near-cognate or non-cognate ternary complexes take on average 4.6 ms and 1.4 ms, respectively. We did not consider mis-incorporation events due to their low overall likelihood (<1% probability).

Next, recognizing that translation elongation takes place within a crowded cytoplasmic milieu, we developed a molecular-mechanistic model for how protein synthesis occurs as a physical process. To start, we estimated the smallest volume of cytoplasm sufficient to enable protein synthesis. We assumed that the molecules required for protein synthesis are homogeneously distributed within the nucleoid-excluded cytoplasm. This volume is ultimately determined by the concentration of ternary complexes as the most-limiting species. So defined, each ‘translation voxel’ contains exactly one of each of 42 unique ternary complexes, one or more ribosomes, and many native proteins (represented as average-sized molecules that represent all other surrounding proteins) (**Figure 2D**).

Brownian diffusion allows each molecule to sample the voxel volume and encounter one another. We modeled diffusion explicitly as a random walk of each of the molecules throughout the voxel where, due to their finite size, they exclude one another entropically or, in the case of a ternary complex and unbound ribosome pair, initiate a reaction (**Methods**; **Table S4**). The benefits of explicit modeling (rather than using prior approaches that insert a diffusion coefficient gleaned from experiment) are that we can follow the detailed motion of each translation molecule, and that diffusivity increases or decreases automatically and naturally with changes in size and crowding (**Appendix A**).

Due to non-uniform codon usage and non-uniform relative abundance of each type of ternary complex (**Table S6**) as well as stochastic variation in the physical distribution of translation molecules in cytoplasm, for any given cell-wide condition, individual translation voxels should be expected to vary in the exact combinations of unique ternary complexes and elongating ribosomes. For example, a translation voxel might contain more than one of a highly abundant tRNA. Accordingly, starting from our basic translation voxel platform we constructed ensembles of thousands of translation voxels to capture the natural distribution of chemical identities and spatial configurations that, together, better represent the natural variation expected within cytoplasm. We used these more-accurate voxel ensembles to examine the physical and chemical mechanistic relationship between growth rate and elongation rate by simulation (below).

### II. Stoichiometric crowding accompanies faster protein synthesis

We gathered and analyzed well-established experimental data for cell mass, cell volume, and the sizes and abundances of ternary complexes, ribosomes, and proteins in cells across growth rates (Dennis and Bremer, 2008; Volkmer and Heinemann, 2011; Dong et al., 1996; Pedersen et al., 1978; Schmidt et al., 2015; Woldringh and Nanninga, 1985) (**Methods, Figure S1, Tables S1–S4**). We deduced that, as growth rate increases (from μ = 0.6 to 3.0 dbl/hr) and translation elongation speeds up (from 12 to 21 aa/s), ternary complexes and ribosomes monotonically increase in number by nearly an order of magnitude, while proteins monotonically increase by three-fold (**Figure 3A**). We remark that occupied volume fraction can change even if overall mass density remains constant in any physical system (**Figure S18, Supplementary Note S5**). In particular in *E. coli*, while there have been conflicting reports of overall mass density changing or remaining constant over different ranges of growth rate (Dennis and Bremer, 2008; Oldewurtel et al., 2021), the occupied volume fraction changes regardless and thereby impacts packing and diffusion of molecules.

**Figure 3.**
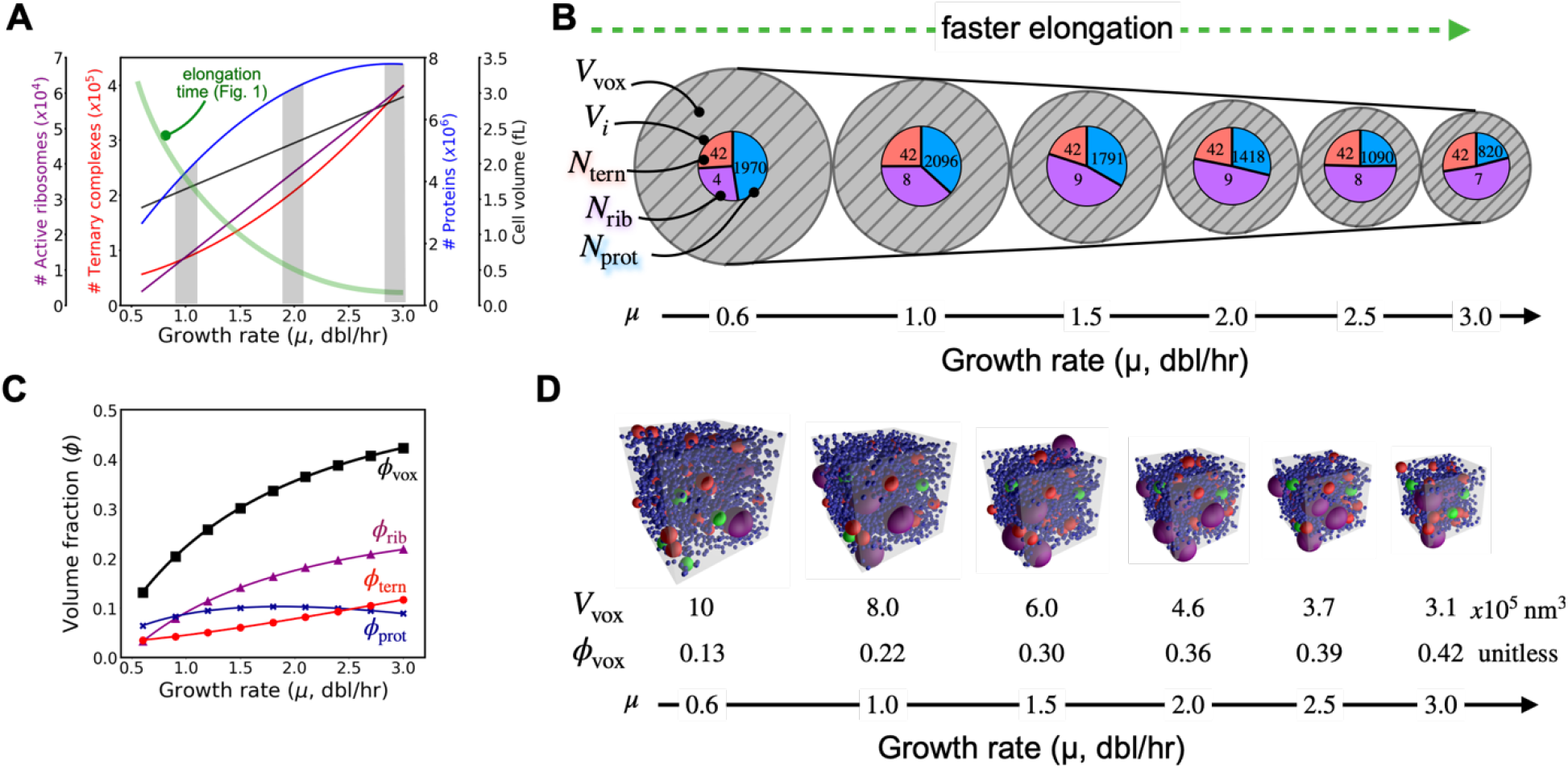
The relative abundances, concentrations, and volume fractions of translation molecules change as growth quickens. **(A)** Colloidal stoichiometry. Experimental observations of active ribosomes, ternary complexes, and protein abundances in *E. coli*, as well as *E. coli* cell volume, reveal varying levels of increase with increasing growth rate (gray bars highlight values at particular growth rates). **(B)** Stoichiometric crowding. An abstracted representation of translation voxels as a function of growth rate reveals that differential changes in molecular abundances are accompanied by an overall increase in crowding (i.e., stoichiometric crowding). The volume of translation voxels (*V*_vox_, hatched gray circles) decreases while the total volume of constituent biomolecules (concentric pie charts) remains relatively constant. The total number of each particular type of biomolecule species (*N*_tern_, *N*_rib_, and *N*_prot_, shown within corresponding colors of the pie chart – red: ternary complexes, purple: ribosomes, blue: proteins) in a given translation voxel as well as the total volume each biomolecule species occupies (*V_i_*, the area of corresponding colors within the pie chart) change at different growth rates. **(C)** The volume fractions (*ϕ_i_* = *V_i_*/*V*_vox_) of ribosomes (*ϕ*_rib_), ternary complexes (*ϕ*_tern_), and proteins (*ϕ*_prot_) change differently with increasing growth rate, leading to an overall increase in the total occupied volume fraction of translation voxels (*ϕ*_vox_). **(D)** Representative snapshots of translation voxel simulations at increasing growth rates, along with their respective volumes (*V*_vox_) and volume fractions (*ϕ*_vox_).

We described the coupled abundances and volume fraction of each constituent biomolecule as the ‘colloidal stoichiometry’ of the translation voxel; mathematically, the abundances and volume fraction of each constituent biomolecule *i* is described as N_*i*_ and *ϕ_i_* = *V_i_*/*V*_vox_, respectively, where *V_i_* is the total volume occupied by a particular biomolecule species and *V*_vox_ is the total volume of the voxel. The colloidal stoichiometry of translation voxels – which captures both chemical and physical features of cytoplasm - changes with growth rate. Thus, we hypothesized that growth-rate dependent changes in colloidal stoichiometry might contribute to the speedup of translation elongation.

More specifically, our modeling revealed changes in the colloidal stoichiometry of translation voxels as growth quickens (0.6 to 3.0 dbl/hr): voxels shrink three-fold (*V*_vox_ = 10E5 nm^3^ to 3E5 nm^3^) and become three-fold more crowded (*ϕ*_vox_ = 0.13 to 0.42). However, the growth in packing fraction is not uniform across species: as voxels shrink in size the number of ribosomes doubles while the number of proteins halves (**Figure 3B, 3D**). That is, increased total crowding is dominated by ribosomes: the volume fraction of ribosomes increases by seven-fold, more than double that of ternary complexes (*ϕ*_rib_ = 0.03 to 0.22; *ϕ*_tern_ = 0.04 to 0.12). Proteins dominate the packing fraction at low growth rates but then plateau (*ϕ*_prot_ = 0.06 to 0.10 for μ = 0.6 to 2.0 dbl/hr; *ϕ*_prot_ = 0.10 to 0.09 for μ = 2.0 to 3.0 dbl/hr), thus contributing minimally to the overall increase in crowding (**Figure 3C**).

We referred to this growth-rate dependent change in colloidal stoichiometry as ‘stoichiometric crowding,’ which we expected should impact both the interactions and motion of translation molecules at different growth rates. For example, in the growth-rate trends noted above, as growth quickens ternary complexes and ribosomes should encounter each other more frequently relative to encountering proteins. As a second example, the distribution of molecule sizes matters: for a fixed total volume fraction the diffusion of individual particles is faster in a suspension of large versus small particles and, as the most dominant particle size shifts from smaller to larger, diffusion of all particles speeds up (Farris, 1968; Gonzalez et al., 2021; Lionberger, 2002); since ribosomes increase in their relative volume fraction with growth rate, voxels should mix more quickly for any given total volume fraction as growth quickens.

### III. Physical transport of ternary complexes accounts for most of elongation latency

We next sought to better understand and quantify any such impacts of crowding and composition on transport rates, reaction rates, and elongation rates overall. To do so we established a baseline quantification of the relative importance of physical transport to chemical reactions in setting elongation latency. We constructed translation voxels, from very simple to biologically faithful forms, and analyzed expected transport, reaction, and elongation latencies by simulation. From this we inferred the mechanisms by which colloidal stoichiometry regulates overall elongation latency as a function of growth rate. We used a slow growth rate (0.6 dbl/hr) as a benchmark, where bulk elongation takes about 87 ms on average (**Figure 1**).

More specifically, we first studied transport and reaction dynamics in detail using an idealized case in which only a single cognate ternary complex interacts with one matching ribosome (**Figure 4A**). Here, there is no competition with mismatching ternary complexes, no other molecules blocking the way, and the sought-after ribosome is unbound; rather, a lone ternary complex searches pure cytosolic fluid for a waiting matching ribosome, an idealized scenario often depicted in ‘textbook’ representations of translation elongation (e.g., Sannuga and Ramakrishnan, 2004). As expected, we found that transport latency – the time a ternary complex spends not bound to a ribosome, diffusively searching for a match – is nearly instantaneous 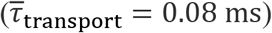 and that nearly all of the elongation process is taken up by reaction latency – the time a ternary complex spends bound to ribosomes 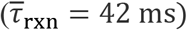 (**Figure 4A**). While such a result seems to support the conclusion that chemistry alone determines translation elongation rate (**Figure 2C**), the notion of a two-molecule translation voxel operating at 3% volume fraction – far below physiological conditions – is unrealistic (**Figure 3C**).

**Figure 4.**
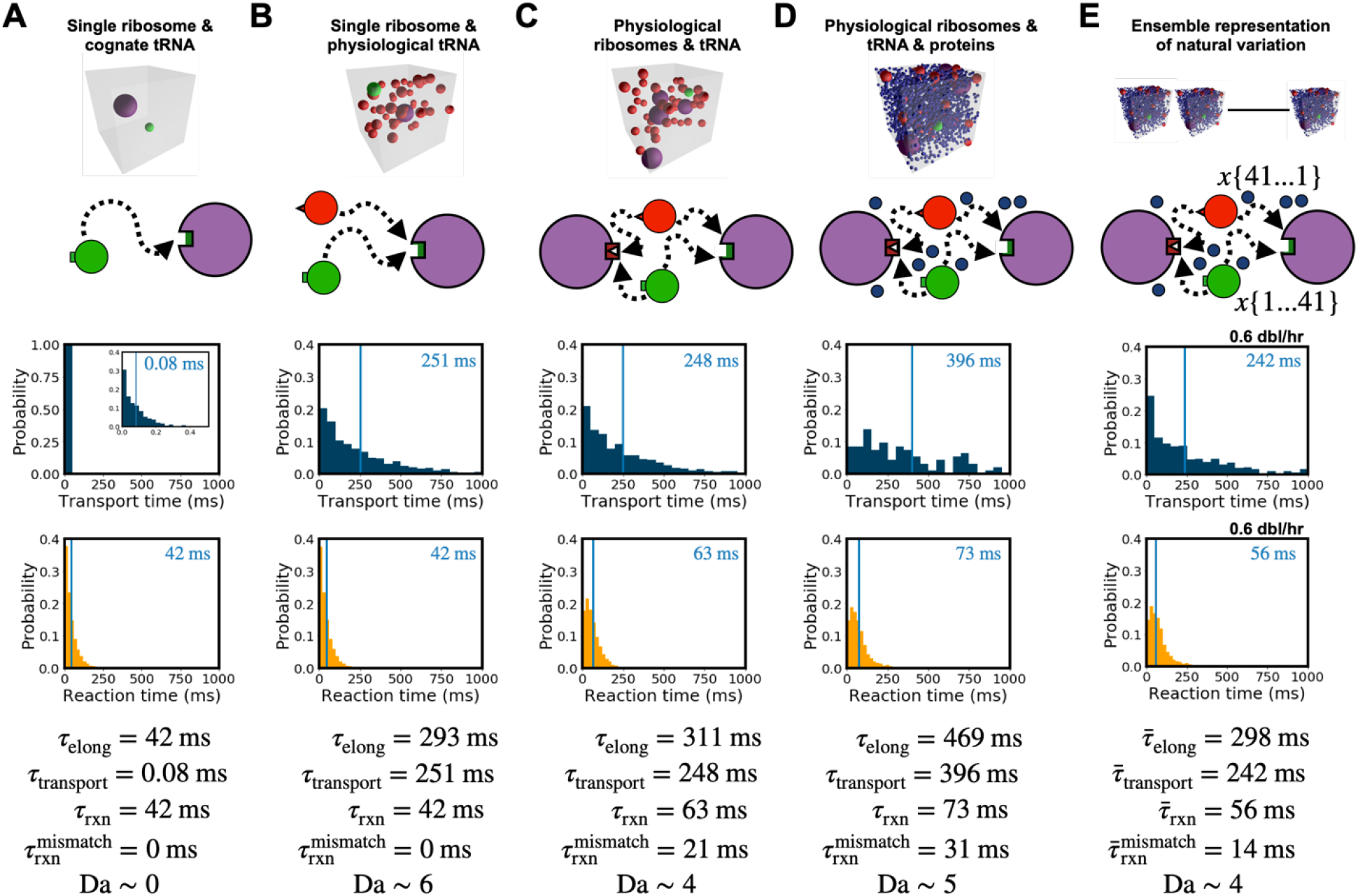
Most of the latency in translation elongation arises from physical transport of ternary complexes. **(A-E)** Simulation snapshots, model schematics, and simulation results (top to bottom) for increasingly realistic (left to right) translation voxels at growth rate μ = 0.6 dbl/hr. In each plot the average latency is marked by a blue vertical line and displayed on the top right in milliseconds. **(A)** A highly simplified translation voxel containing only a single ribosome and cognate ternary complex. **(B)** A translation voxel containing a single ribosome and 42 ternary complexes. **(C)** A translation voxel with 42 ternary complexes and four ribosomes. **(D)** A translation voxel with 42 ternary complexes, four ribosomes, and 1970 proteins. **(E)** An ensemble of translation voxels that capture the expected natural variation in cognate, near-cognate, and non-cognate ternary complexes due to non-uniform ternary complex and codon abundances coupled with spatial stochasticity. The standard errors in the estimate of the mean relative to the mean for transport latency and elongation latency are 3% for (A), (B), and (C), 9% for (D), and 6% for (E) while for reaction latency are all below 1%.

We next added a physiologically correct number of ternary complexes to the voxel (i.e., one cognate, 41 non-cognate) such that each ternary complex competes to reach and bind to the ribosome (**Figure 4B)**. We found that the total reaction time remains the same 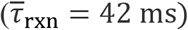. However, the transport latency of the cognate ternary complex increases markedly 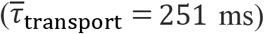 and is greater than reaction latency, supporting arguments that translation elongation requires substantial reactant transport time. The increased transport time is also coupled to reactions: a cognate ternary complex must “wait” to bind with the ribosome while that ribosome is already bound to non-cognate ternary complexes; thus, there is an interplay between reactions and transport.

We then added a physiologically correct number of ribosomes to the translation voxel, holding the ternary complex population fixed at one cognate and 41 non-cognates (**Figure 4C**). Only one ribosome was available for a matching reaction with the cognate ternary complex; the other ribosomes were mismatching for all ternary complexes. Having just one matching ribosome means we need track only the elongation events of a single ribosome, while simultaneously tracking mismatching events at other ribosomes; this approximation provides a lower-bound estimate for bulk elongation rates while allowing a more accurate accounting of transport and reaction effects (below). We found by simulation that transport latency remains similar to voxels containing a single ribosome 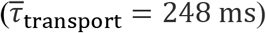 because the single matching ribosome remains bound for almost the same amount of time, indicating that the cognate ternary complex still needs to ‘wait’ nearly as long (i.e., the indirect impact of mismatching reactions). A slight decrease in transport latency can be attributed to some non-cognate ternary complexes being bound by mismatching ribosomes, meaning fewer non-cognate ternary complexes are available to occupy the matching ribosome. However, the reaction latency of the cognate ternary complex increases 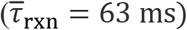 due to the direct impact of mismatching reactions: cognate ternary complexes spend more time in futile interactions with the more abundant mismatching ribosomes.

Next, we added a physiologically correct abundance of proteins to the translation voxel resulting in further increases in both transport and reaction latencies 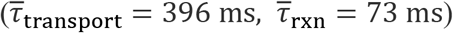 (**Figure 4D**). These predicted increases arose because proteins increase the number of mismatching reactions by trapping non-cognate ternary complexes near ribosomes, which in turn both promotes repeated mismatch reactions and reduces cognate ribosome availability.

Finally, we represented the expected statistical variation in cytoplasm by constructing thousands of different voxels that, together, capture the physiological distribution of relative abundances of translation molecules as reported in the literature (**Methods**). Specifically, in *E. coli*, there are 42 unique ternary complexes, each with its own abundance, and 64 codons, each with its own usage rates, and these are present in many permissible combinatoric configurations in translation voxels throughout cytoplasm (**Figure 2D**). We randomly sampled all permissible configurations using reported *E. coli* codon usage and whole-cell tRNA abundances (**Methods, Table S6, Figure S4**). Recognizing that bulk elongation measurements correspond to the time needed to complete as many successful reactions as ribosomes are in a voxel, we computed the transport, reaction, and elongation latencies of each translation voxel from the time taken for a single matching reaction within the voxel. A weighted average of the per-ribosome latency for all permissible translation voxels corresponds to the typical time for just one matching reaction to occur in a voxel and thus provides a lower-bound estimate of bulk experimental elongation time as obtained from cellular measurements (**Methods**). While our prior simulations (**Figures 4A – 4D**) included only non-cognate and cognate ternary complexes, our calculations of the ensemble latencies (**Figure 4E**) also included the more detailed classification of some ternary complexes as near-cognate, which affects system dynamics further because near-cognates are well-known to have a slower rejection-time than non-cognates (**Table S5**, **Figure S2**). We monitored transport, reaction, and elongation latency during simulation in each of these thousands of voxels, and computed a weighted-average value for transport, reaction, and elongation latency. We found that the weighted-average transport, reaction, and elongation latencies decrease in the ensemble representation 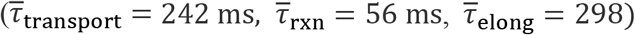 (**Figure 4E**), compared to the single-voxel simulation (**Figure 4D**). This across-the-board decrease in latencies emerges naturally from the majority of voxels in which there is more than one cognate ternary complex, partly a result of the biological phenomenon of more frequently used codons being associated with more abundant cognate tRNAs (**Figure S8**).

Quantitatively, the Damköhler number – the ratio between the latency of transport and reaction (Da = *τ*_transport_/*τ*_rxn_) – highlights the dependency of translation elongation latency on physical transport relative to chemical reaction. In our simplest model, Da ~ 0, suggesting that reaction latency dominates elongation latency. However, our increasingly accurate models estimate Da ~ 6, Da ~ 4, Da ~ 5, and Da ~ 4 respectively (**Figures 4B – 4E**). We thus concluded that processes that modulate transport latency play a dominant mechanistic role in regulating the overall speed of translation elongation.

### IV. Stoichiometric crowding speeds up translation elongation

We returned to the puzzle of what mechanism(s) might cause the productivity of individual ribosomes to increase with increasing growth rates. We first evaluated the impact of stoichiometric crowding on transport latency by considering both molecule proximity (i.e., how close molecules are to one another) and molecular mobility (i.e., how fast molecules move). We also evaluated the impact of stoichiometric crowding on reaction latency by considering both local availability (i.e., to what extent ternary complexes are free from repeated mismatching reactions) and global availability (i.e., to what extent ternary complexes are free from mismatching reactions generally). Taken together we determined if and how each of these coupled physico-chemical mechanisms might regulate elongation latency as a function of growth rate.

We hypothesized that crowding should tend to reduce transport latency because ternary complexes need to search smaller volumes to find a matching ribosome. To explore this idea, we computed the average surface-to-surface distance between ternary complexes and their closest ribosomes (i.e., the shortest distance a ternary complex needs to travel to find a ribosome) across hundreds of translation voxels at multiple growth rates (**Methods**). We found that stochiometric crowding brings ternary complexes and ribosomes five-fold closer on average (16 nm to 3 nm), which supports our hypothesis (**Figure 5A, left axis**). But stoichiometric crowding could also increase transport latency as ternary complexes become hindered in their motion and thus take longer to search. To explore this second idea, we examined the influence of stoichiometric crowding on molecule mobility by estimating the viscosity of cytoplasm as well as the hindered diffusivity of ternary complexes, ribosomes, and proteins via simulation of hundreds of translation voxels at multiple growth rates (**Figure 5A, right axis; Figure S3; Methods**). We found that stochiometric crowding increases viscosity monotonically (1.0 to 2.4, normalized to viscosity at *μ* = 0.6 dbl/hr) while reducing diffusivity monotonically for all biomolecules (e.g., the diffusivity of ternary complexes, *D*_tern_, slows from 35 μm^2^/s to 16 μm^2^/s), which would support the opposite conclusion: that crowding should hinder transport. Recognizing this competition between an increased proximity reducing transport latency and an increased viscosity increasing transport latency, we more systematically considered the contribution of each mechanism to transport as stoichiometric crowding increased due to increased cell growth rate.

**Figure 5.**
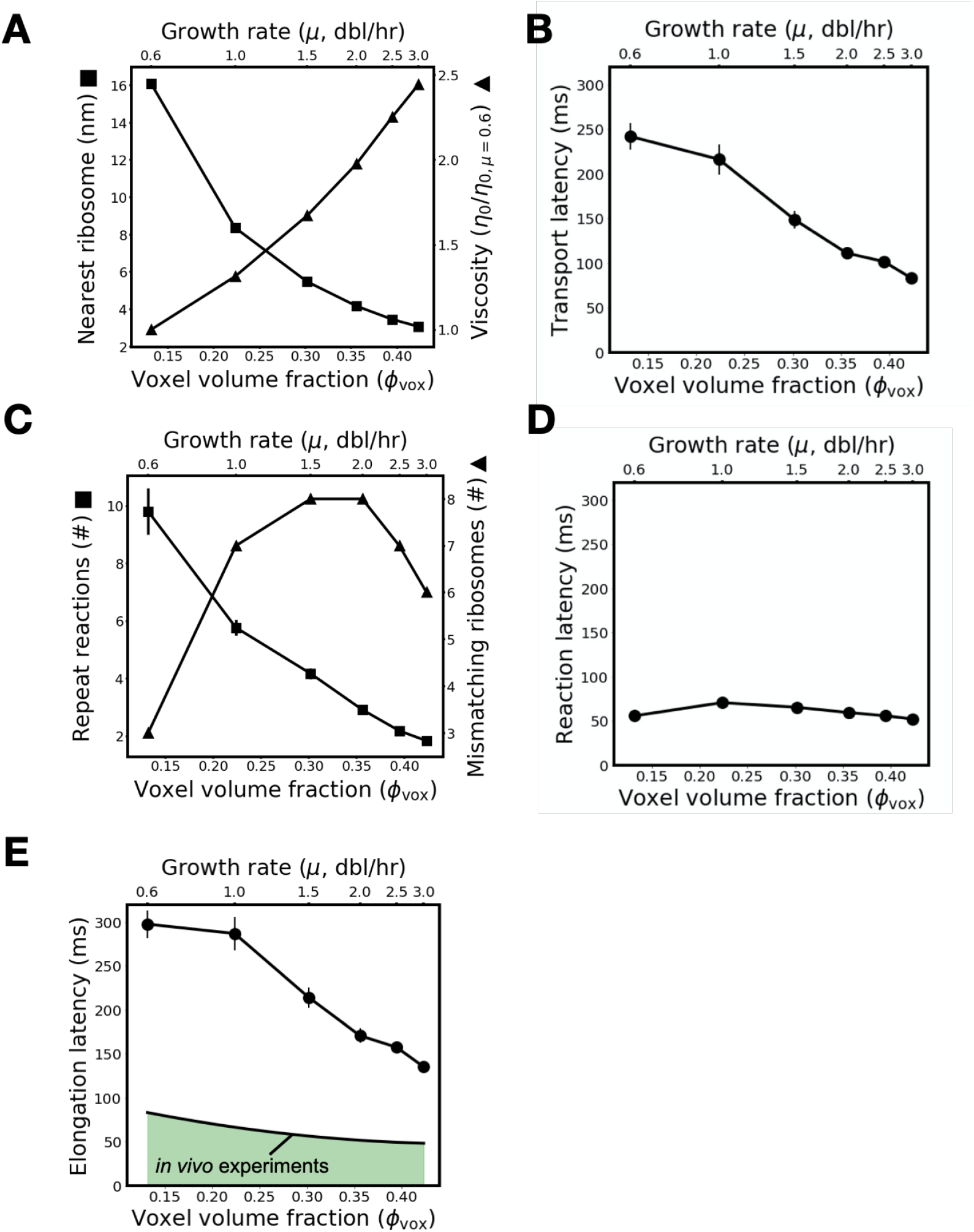
Stoichiometric crowding reduces both intermolecular distances and transport latency resulting in increasingly productive ribosomes as growth rate increases. **(A)** As crowding and growth rate increase (x-axes) ternary complexes become closer to their nearest ribosome (left y-axis) and translation voxel viscosity increases (right y-axis). Distance is reported as a surface-to-surface estimate. Viscosity is reported normalized to viscosity at μ = 0.6 dbl/hr. **(B)** Simulation results showing that transport latency (y-axis) decreases with increased crowding and growth rate (x-axes). **(C)** As crowding and growth rate increase (x-axes) the average number of repeat reactions between ternary complexes and ribosomes decreases (left y-axis) while the absolute number of mismatching ribosomes in a translation voxel first increases then decreases (right y-axis). **(D)** Simulation results showing that reaction latency (y-axis) first increases then decreases with increased crowding and growth rate (x-axes). **(E)** Simulation results showing that the predicted absolute elongation latency (filled circles, solid line) decreases with increased crowding and growth rate (x-axes). Experimentally measured per-ribosome elongation latency (solid line upon green area; replotting of Figure 1) also speeds up with growth rate but is faster than predicted across all growth rates. The standard errors in the estimate of the mean for all model results (A-E) are shown (error bars).

For example, we simulated ensembles of translation voxels representing the full statistical distribution of ternary complexes and codon abundances from low- to high-growth rates (**Methods**). We found that transport latency monotonically decreases with stoichiometric crowding 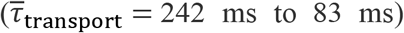 (**Figure 5B**). We deduced that, mechanistically, crowding drives faster transport because reducing the search distance between ternary complexes and ribosomes is more important than increased viscosity. However, while this net decrease in transport latency will decrease elongation latency overall, it could be that coupled changes in reaction latency either reverse or reinforce this trend.

Thus, we examined two mechanisms that could modulate reaction latency. First, we recalled that proteins can induce repeat reactions by trapping ternary complexes and ribosomes together (**Section III**). These repeat reactions should reduce the local availability of ternary complexes, making it more difficult for ternary complexes to find matching ribosomes, driving up reaction latency. To examine whether increased stoichiometric crowding amplifies this effect, we tracked the number of times ternary complexes consecutively re-react with the same ribosome following a mismatching reaction across hundreds of translation voxels at varying growth rates. We were surprised to find that repeated reactions decrease five-fold (from ~10 to ~2 repeat reactions per ribosome on average) as growth rate quickens and total crowding increases (**Figure 5C, left axis**). We resolved this apparent paradox by recognizing that the increased crowding arises primarily due to tighter packing of ribosomes, while the volume fraction of proteins hardly changes. We deduced that this provides more local ribosome alternatives (i.e., higher local availability) for ternary complexes but with no increase in trapping by proteins (**Figure S6**). However, having more ribosomes, regardless of how well packed they are, provides more opportunities to preoccupy ternary complexes in mismatch reactions, reducing ternary complex global availability, which should drive up reaction latency. We found that the number of mismatching ribosomes in a voxel first increases and then decreases with growth rate, which should contribute an initial increase and then decrease in reaction latency as growth rate increases (**Figure 5C, right axis**). Taken together, the total impact on reaction latency depends on the relative strengths of each of these effects.

We next computed reaction latencies across our ensembles of translation voxels, capturing how physiological variation influences the competition between local and global availability. We found that total reaction latency increases 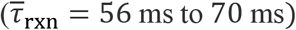 at low but increasing growth rates and then decreases monotonically thereafter 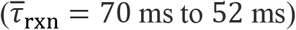 (**Figure 5D**). Low global availability of ternary complexes dominates at low growth rates, slowing reaction latency as growth rate increases. However, at higher growth rates, the increase in both global and local availability combine to drive down reaction latency. Overall, the growth-rate trend in reaction latency (**Figure 5D)** follows ternary complex global availability (**Figure 5C, right axis**). Practically, even with a 25% increase that subsequently reverses, reaction latency changes only 7% over low-to-high growth rates, suggesting that transport plays the more substantial role in speeding elongation.

Indeed, the quantitative speedup of reaction latency with growth rate (**Figure 5D**) is minor compared to the corresponding speedup of transport latency (**Figure 5B**), indicating that transport mechanisms should be expected to dominate over reaction mechanisms in regulating the growth-rate dependent productivity of individual ribosomes. Our ensemble simulations show that the dominance of transport manifests in the total elongation latency as a monotonic speedup of elongation with growth rate 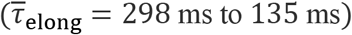, recovering the experimental trend of faster elongation at higher growth rates 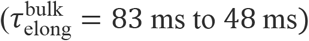 (**Figure 5E**).

Finally, although our model correctly predicts and recovers the qualitative behavior and overall trend (i.e., an increase in ribosome productivity with increasing growth rate), we noted that our unfitted bottom-up modeling and simulations result in absolute predictions of translation elongation latencies that are ~3-fold too slow compared to experimental observations (**Figure 5E**). Thus, we conducted sensitivity analyses in which chemical kinetic rates were fitted to match observed overall translation elongation latency (**Supplement**). We found that the speedup in translation elongation is insensitive to changes in the chemical kinetics of translation elongation (**Figure S13**).

## Discussion

The observed increase in individual ribosome activity as growth quickens has not been explained mechanistically (**Figure 1**). While many have focused attention on how translation initiation contributes to protein synthesis latency, we were intrigued by how the individually fast steps of translation elongation add up during translation elongation and should dominate overall process latency (**Supplement**). In the context of translation elongation, while prior studies of protein synthesis rates focused on chemical kinetic measurements or kinetics-based modeling, there have been persistent signals that Brownian diffusion of translation molecules plays a role in setting elongation rates.

With this in mind, we explored the idea that either chemistry or physics, or both, contribute to the speedup of translation elongation utilizing dynamic simulations. We proposed that reactions between ternary complexes and ribosomes are nontrivially coupled to their physical transport and that understanding this coupling is essential to explaining increased ribosome productivity at faster growth rates. To systematically interrogate the role of coupled physico-chemical processes in translation elongation, we adapted an open-source simulation tool to accurately represent transport of and interactions between translation molecules in cytoplasm. A key aspect of our approach is the robust modeling of Brownian motion and colloidal-scale particle interactions such that these molecules undergo the inertialess physical encounters appropriate to the colloidal regime. We defined translation voxels as naturally emergent from the constituent biomolecules required for translation and captured the natural distribution of chemical identities and spatial configurations of translation molecules in cytoplasm by constructing ensembles of thousands of voxels. We monitored in simulations the reactions and transport of molecules in these voxel ensembles to study the physical and chemical mechanistic relationships between growth rate and elongation rate (**Figure 2**).

We found that transport latency – the time ternary complexes spend searching for cognate ribosomes – is an essential component of elongation latency. Furthermore, we predict that transport latency dominates over reaction latency – the time ternary complexes spend reacting with ribosomes (**Figure 4**). Indeed, physical transport of individual ternary complexes accounts for ~80% of elongation latency. By examining the elongation process as growth rate increases, we identified two competing mechanisms that underlie transport latency – proximity between ternary complexes and ribosomes that sets search distance, and cytoplasmic crowding that sets diffusive speed. Additionally, we observed that translation molecules become three-fold more crowded with increasing growth rate, suggesting that, beyond any absolute increase in the abundance of translation machinery, the machinery itself becomes packed closer together. The abundance and packing are physical as well as chemical (i.e., colloidal stoichiometry) and their changing due to changes in growth rate is a phenomenon we call stoichiometric crowding (**Figure 3**). Overall, we find that increased packing at faster growth rates improves proximity that, along with changes in stoichiometry, increases the frequency of cognate reactions that in turn increases individual ribosome productivity (**Figure 5**), revealing a mechanistic explanation for why individual ribosomes can produce proteins more quickly in faster-growing cells.

Our colloidal-scale, mechanistic conclusions complement existing phenomenological modeling work that describe how protein synthesis and growth can be predicted by resource allocation kinetics and optimization (Erickson et al., 2017; Klumpp et al., 2013; Kostinski and Reuveni, 2020). We also help resolve a paradox arising from the observations by Klumpp et al., which suggested slower diffusion of ternary complexes leads to slower growth but then cannot explain faster growth in these more crowded conditions (Klumpp et al., 2013). Here, we explain how slower diffusion accompanies faster growth, which is to be expected given increased crowding in faster growing *E. coli* (**Figure 5; Supplementary Note S4**). Translation speeds up as diffusion goes down: closer proximity (i.e., molecules closer together at higher crowding) outpaces reduced diffusivity, resulting in faster translation elongation and thus faster growth overall. Crowding favors ribosomes and translation molecules at faster growth rates: the more-crowded cytoplasm has a higher relative abundance of translation molecules compared to other proteins (**Figure 3**). The resulting improved proximity and changed stoichiometry speed up translation.

We stress-tested our model and confirmed that the speedup of elongation requires physical transport, and that our prediction of speedup is robust to changes in the values of the input chemical parameters. We also found that increasing three-fold the values for all nine *in vitro* literature values for chemical kinetics parameters closes the quantitative gap between predicted and observed elongation rates (**Figure S13**). This suggests the rather straightforward chemistry-only explanation for the gap: *in vitro* measurements being “off” by ~300%, uniformly across all nine parameters. But, interestingly, we also found a 30-fold increase in only the ternary complex unbinding rate (*k*_1r_) – the only reaction that takes place exclusively outside the ribosome – could also close the quantitative gap (**Figure S13, Figure S9**), suggesting there may be a mechanism involved *in vivo* that quickens ternary complex exchange or one that obviates the need for fast rejection (i.e., a mechanism for favoring matching reactions near to the ribosome).

Overall, our model reveals new opportunities for discovery. For example, better representation of electrostatic and hydrodynamic interactions or detailed molecular shape and orientation for site-specificity may be useful. More specifically, attractive interactions between the ribosomal L7/L12 domain and ternary complexes (Mustafi and Weisshaar, 2018) or between cognate ternary complexes and mRNA (Grosjean and Chantrenne, 1980) could have the effect of pre-loading or pre-sorting ternary complexes. As a second example, hydrodynamic models of small and large particles confined in a cavity show that both types of particles tend to concentrate near the cavity surface with minor impact on the mobility of small particles (Gonzalez et al., 2021), indicating that ternary complexes and ribosomes may concentrate by the cell membrane (not currently represented in our model) and effectively improve in proximity to each other.

More generally, our work supports exploration of the role of coupled, colloidal-scale physico-chemical interactions in cytoplasm. For example, we predicted that ternary complexes and ribosomes should be up to five-fold closer together in faster growing cells (**Figure 5A**), a major shift in the colloidal-scale structure of cytoplasm that can be expected to modulate molecular interactions broadly across the cytoplasm. Such colloidal-scale structure is being increasingly measured experimentally (e.g., ribosome spatial positioning via cryo-electron tomography of entire cells) and merits increased attention for its role in cytoplasm behavior (Ortiz et al. 2006, Robinson et al. 2007). As a second example, the phenomenon of repeat reactions we found critical to the speedup of protein synthesis has also been identified as critical to efficient activation of Mitogen-Activated Protein (MAP) Kinases (Takahashi et al., 2010), suggesting a wider role for repeat reactions in cell functions. One can also infer the possibility that stoichiometric crowding with changing growth rate may impact cell signaling in general. As a third example, using our model we predicted that the viscosity of the nucleoid-excluded cytoplasm increases up to 2.5-fold with growth rate, which indicates a decrease in the mobility of all constituent molecules. Such a broad growth-rate dependent shift in colloidal-scale dynamics may suggest currently unappreciated forms of physical regulation in cells and motivates a renewed analysis of diffusive processes in cells with consideration to volume fraction and growth rate. Broadly, a more complete understanding and representation of the colloidal-scale dynamics that underlie cellular processes can offer a practical first-principles foundation for systems biology and whole-cell modeling (Karr et al., 2012; Macklin et al., 2020; Thornburg et al., 2022).Advances in both computational modeling and experimental technique are needed to improve the accuracy of our predictions and promote broader exploration for how coupled colloidal-scale physics and chemistry in cytoplasm might regulate cellular behaviors. For example, dynamic simulation of the motion of solvent-suspended particles requires discretization of the time domain, where equations of motion are integrated forward in time. The selection of time step size impacts not only computational expense but also the fidelity of particle encounters where, for example, too-large timesteps can produce pathological displacements in response to steeply attractive or repulsive forces (e.g., the hard-sphere repulsion that represents entropic exclusion is singular at particle contact), a phenomenon that becomes more severe as crowding increases. Here, we performed a careful study to prioritize physical accuracy first and then optimize efficiency by, for example, developing a kinetic scaling method that leveraged the natural disparity between diffusive and reactive timesteps (**Figures S3, S5, S10, S14**). Even so, further capturing the complexity of cytoplasm (e.g., protein polydispersity, general protein-protein interactions, polysome dynamics, or cell-cycle dependency) will ultimately require modeling other microscopic forces at play in cytoplasm, including electrostatic or hydrodynamic interactions or membrane confinement, all of which lead to many-body interactions that increase computational expense. Modeling such forces, in addition to other molecular details like shape, softness, flexibility, and site-specificity, is becoming possible with other algorithms such as Stokesian dynamics for large or confined systems (Aponte-Rivera et al., 2018; Gonzalez et al., 2021; Maheshwari et al., 2019; Ouaknin et al., 2021; Zakhari et al., 2017), but will require substantial integration and iteration with experiments as well as improvements in computational efficiency to achieve accurate simulations over the timescales of cellular behavior (Endy and Brent, 2001). Capturing detailed molecular dynamics, such as those involved with ternary complex-ribosome binding and reactions, will also necessitate multi-scale modeling and experimentation from atomic to cellular scales.

In closing, we note that protein synthesis is inextricably tied to growth rate and fitness; cells cannot grow more quickly than they can reproduce their proteome, including the proteins that remake the proteome. Since stoichiometric crowding facilitates faster protein synthesis, could further crowding enable still-faster growth or, conversely, limit how fast cells can grow? To speculate, as a simple extension to our modeling, we projected what cytoplasm of faster-than-observed growing *E. coli* would look like (**Figure S11, Methods**). We found that *E. coli* would eventually reach maximum packing (i.e., with little to no space for molecular mixing) as growth rate continues to increase (**Figure 6A**). That such hypothetical growth rates are not observed suggests, among other possibilities, that as growth rate increases beyond the maximum observed, the beneficial effects of increased crowding (e.g., increased proximity) become outpaced by the deleterious effects of less available free volume (e.g., further increased viscosity) (**Figure 6B**). Our theoretical observation expands upon a past proposal that growing cells optimize total protein volume fraction for both reaction and transport (Dill et al., 2011) and hints that stoichiometric crowding may be linked to fitness and evolutionary fine tuning. If so, then we would expect that genes encoding currently unknown functions may serve to establish a physical, as well as chemical, basis for fitness (e.g., proteome polydispersity) undergirding cellular behavior broadly.

**Figure 6.**
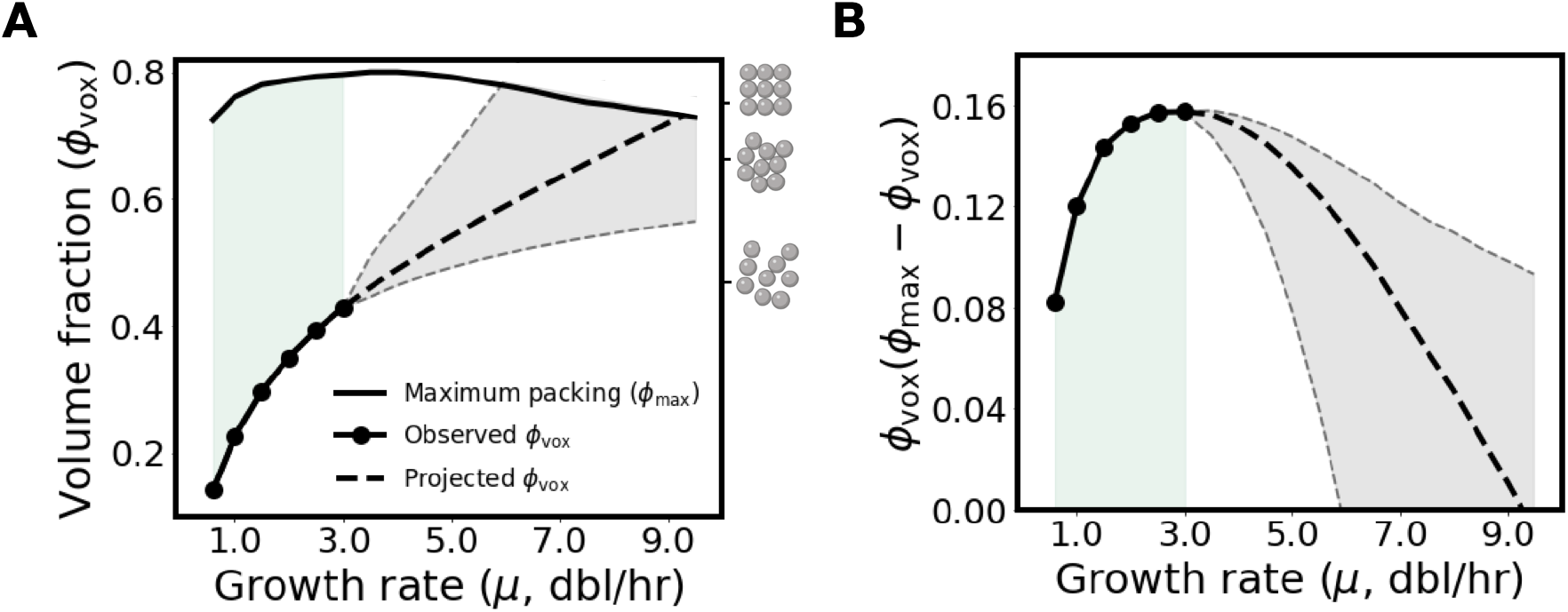
Stoichiometric crowding has diminishing returns that may impose a physical limit on growth rate. **(A)** Volume fraction of translation voxels at observed growth rates and projected growth rates compared to theoretical maximum random close packing (maximum packing changes with size polydispersity (Farr and Groot, 2009) and size polydispersity changes with growth rate). Bounds for volume fraction at projected growth rates are shown (gray dashed line and shading). Maximum packing increases across observed growth rates (green shading). The volume fraction for the most-crowded observed voxel (ϕ_vox_ = 0.42, μ = 3.0 dbl/hr) is shown (right axis, bottom schematic). For reference, the volume fractions at which long-term molecular motion in monodisperse suspensions is hindered or halted due to random close packing (ϕ = 0.64) or crystallization (ϕ = 0.74), respectively, are also shown (right axis, top and middle schematic). **(B)** The product of voxel volume fraction (ϕ_vox_) and remaining available volume fraction (ϕ_max_ – ϕ_vox_) increases across observed growth rates (green shading) before decreasing across faster-than-observed growth rates.

## Methods

### I. Construction of a representative translation voxel

We developed computational representations of translation voxels by analyzing the abundances and sizes of molecules comprising *E. coli* cytoplasm in relation to overall cell volume and mass. Where needed we inferred abundances across growth rates by fitting polynomials to reported measurements (described below).

#### I.1. Calculation of biomolecular abundances in cells

We computed the average abundances of ribosomes (*N*_rib_) and ternary complexes (*N*_tern_) in single cells across physiological growth rates using existing data from literature (**Table S1**). To compute the abundance of proteins surrounding ribosomes and ternary complexes, we first calculated the total dry mass of proteins in cytoplasm, *M*_cytoplasm,prot_ (**Equation 1**). Protein mass encompasses the mass of all biomolecules in cytoplasm other than ribosomes, ternary complexes, mRNA, and DNA:

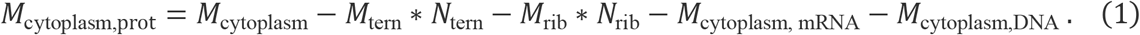

The masses of ternary complexes (*M*_tern_) and ribosomes (*M*_rib_) were specified using known molecular structures and the total mass of mRNA, DNA, and cytoplasm (*M*_cytoplasm,mRNA_, *M*_cytoplasm,DNA_, and *M*_cytoplasm_, respectively) were taken from literature (**Tables S4, S1**). We then calculated the average effective spherical radius of proteins 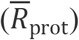, as well as the mass of resulting average-sized proteins 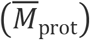, using single-cell *E. coli* mass spectrometry data and average protein density (*ρ*_prot_) (ProteinComputation.py):

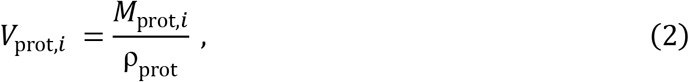

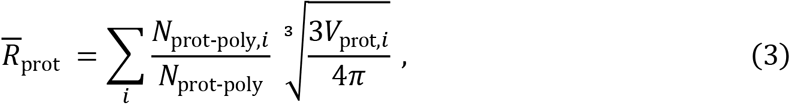

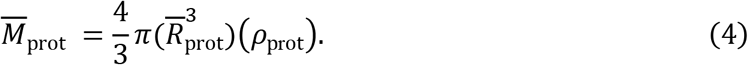

We obtained the mass and abundances of each protein in the *E. coli* cytoplasm (*M*_prot,*i*_ and *N*_prot-poly,*i*_ respectively), as well as the total number of proteins (*N*_prot-poly_), from the mass spectrometry measurements of Heinemann and colleagues (Schmidt et al., 2015). We also computed the volume of each protein in the *E. coli* cytoplasm (*V*_prot,*i*_) using these measurements. We then determined the abundance of proteins in a cell as the ratio of total mass occupied by proteins in cytoplasm (**Equation 1**) and the mass of average-sized proteins (**Figure S1B**),

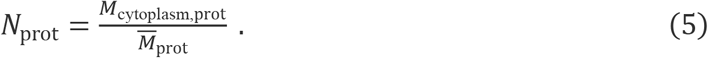

We note that the size polydispersity of proteins is weak – meaning they do not deviate much form the average size (**Figure S16**). Although we show that the presence of proteins deeply influences translation dynamics (**Figure 4, Figure S3**), the fact that proteins are typically much smaller than ribosomes and ternary complexes means that protein size variation is a vanishingly small influence in comparison. As a result, we expect negligible change to ternary complex dynamics and overall translation rates, justifying our representation of protein crowding with an average size.

#### I.2. Calculation of translation voxel size and biomolecular abundances in translation voxels

On a relative basis amino-acid specific ternary complexes are the least-concentrated molecules involved in translation elongation and thus their concentration determines the minimum volume of cytoplasm capable of supporting protein synthesis. Therefore, we determined the size of translation voxels from our estimates of ternary complex abundances (*N*_tern_) and *E. coli* volume (*V*_cell_) across growth rates (**Figure S1, Tables S1, S2**). More specifically, we defined a translation voxel to be the volume of cytoplasm that contains 42 ternary complexes (*N*_tern,vox_ = 42), assuming a spatially homogeneous distribution of ternary complexes within cytoplasm:

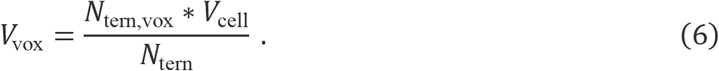

We then calculated the average number of proteins (*N*_prot, vox_) and ribosomes (*N*_rib, vox_) within the translation voxel volume (*V*_vox_) assuming a homogeneous distribution of both species, but with ribosomes excluded by the nucleoid (**Figure S1C**):

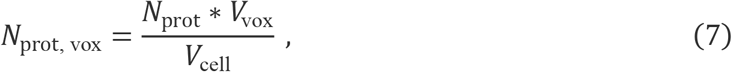

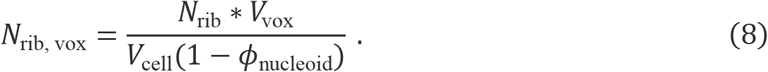

We estimated the volume fraction of the nucleoid (*ϕ*_nucleoid_) based on published values (**Table S3**).

#### I.3. Polynomial regression fitting

We computed polynomial fits for ribosome abundances, ternary complex abundances, cell mass, cell volume, and nucleoid volume using a bootstrapping method that minimized the mean absolute error (**Figure S1A**, TranslationVoxelParameterization.py); we used mean absolute error instead of mean squared error to penalize all variation equally.

### II. Calculation of translation voxel volume fractions and polydispersity

We calculated the volume fraction of ribosomes, ternary complexes, and proteins (**Figure 3C**) using ribosome, ternary complex, and protein abundances in translation voxels; ribosome, ternary complex, and average-sized protein single-molecule volumes; and translation voxel volume:

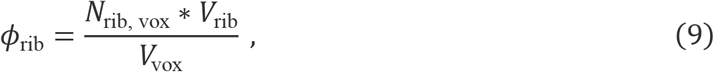

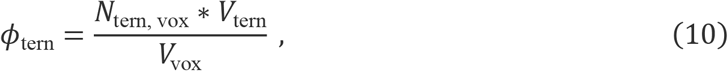

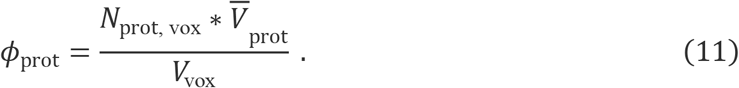

The abundances of ribosomes, ternary complexes, and proteins (*N*_rib,vox_, *N*_tern,vox_, *N*_prot,vox_) in translation voxels as well as the translation voxel volume (*V*_vox_) are as described in **Equations 6, 7, and 8**. The volume of a single average-sized protein 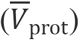 is defined as its mass divided by average protein density (**Equation 4**). We computed the volume of single ribosomes (*V*_rib_) and ternary complexes (*V*_tern_) based on their longest length (i.e., estimating the molecules as spheres with diameters equal to the longest length of the molecules), which we measured from their detailed atomic resolution structures (PDB 4V4Q and PDB 1B23, respectively) (**Table S4**).

We calculated the size polydispersity, *s*, of translation voxels as in Fairhurst, 1999:

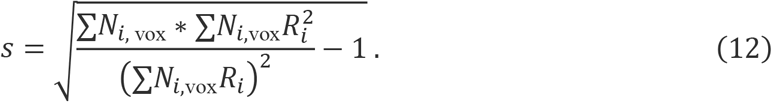

Here, *N*_*i*,vox_ and *R_i_*, correspond to the translation voxel abundances and effective spherical radius of each biomolecule type, denoted by the subscript *i*, respectively (**Table S4**).

### III. Simulation of translation voxels

We simulated transport and reaction of biomolecules within translation voxels using Brownian dynamics and single-molecule reaction kinetics, respectively. We implemented our simulations using “Colloidal Smoldyn”, our adaptation of the open-source simulation software Smoldyn (Andrews et al., 2010). Colloidal Smoldyn accurately represents single-molecule resolution colloidal transport dynamics and reaction dynamics as described in detail in **Appendix A**. The timestep we used in our simulations is Δ*t* = 62 picoseconds.

### IV. Simulation of translation voxels with varying composition

To measure the relative contributions of transport and reactions to protein synthesis rate, we simulated translation voxels with increasingly accurate composition (*μ* = 0.6 dbl/hr) (Results section III, ColloidalStoichiometryEffects.py). Specifically, we simulated five progressively accurate scenarios: (i) a single ribosome & cognate ternary complex matching pair; (ii) a matching pair surrounded by a physiological number of ternary complexes; (iii) a matching pair surrounded by physiological number of ternary complexes and ribosomes; (iv) a matching pair surrounded by physiological numbers of ternary complexes, ribosomes, and proteins; and (v) a statistically representative ensemble of translation voxels each with a physiological number of ternary complexes, ribosomes, and proteins. We performed 900 simulation replicates each for the first three cases and 100 simulation replicates for case (iv); we detail case (v) in the next section. Replicates were assigned random initial conditions, chosen using a Mersenne Twister random number generator with seeds set at multiples of five (i.e., 0, 5, 10, …). Since reaction kinetics inside the ribosome (following codon recognition) are well known and unidirectional (see supplemental figure 2C), we do not explicitly model intra-ribosomal kinetics within voxel simulations. Instead, we modeled reaction kinetics following codon recognition separately by producing and sampling from a distribution of 30,000 post-codon recognition reaction times with the kinetic rates summarized in Table S5.

### V. Construction of statistically representative translation voxel ensembles

To construct statistically representative ensembles of translation voxels (used in Results sections III, IV, and V), we incorporated reported relative abundances of different types of ternary complexes and frequencies of codons among mRNA in *E. coli* at different growth rates (**Table S6**). We computationally constructed 100,000 translation voxels for cells in each of six different growth conditions (0.6, 1.0, 1.5, 2.0, 2.5, and 3.0 dbl/hr), sufficient to represent the statistical distribution of translation voxels across each condition. Individual translation voxels comprise different types of ternary complexes and codon-specific elongating ribosomes, randomly chosen using a Mersenne Twister random number generator with a seed of zero.

For each translation voxel, we randomly picked a single ribosome to track. We then classified ternary complexes in each translation voxel as either non-cognate or cognate to the chosen ribosome, leading to a growth-rate dependent distribution in the abundance of cognate ternary complexes (between zero to forty-two) across translation voxels (**Figure S4**, CognatetRNADistributionCalculation.ipynb). For each of our six modeled growth conditions, we used the distribution of cognate ternary complexes calculated at the closest growth rate (measured at 0.4, 0.7, 1.07, 1.6, 2.5 dbl/hr). The speed with which the single chosen ribosome in a translation voxel successfully finds and reacts with a cognate ternary complex provides a good lower bound for the bulk translation elongation rate; the bulk elongation rate corresponds to the speed with which as many peptide bonds are formed as number of ribosomes, and the speed with which a single ribosome finds and successfully reacts with a cognate ternary complex will typically be faster than the speed of as many successful reactions as ribosomes in the voxel.

### VI. Simulation of statistically representative translation voxels ensembles

To compute the transport, reaction, and elongation latencies of statistically representative ensembles of translation voxels (Results sections III, IV and V), we simulated translation voxels across the six different growth conditions (0.6, 1.0, 1.5, 2.0, 2.5, and 3.0 dbl/hr). Statistically representative ensembles of translation voxels correspond to the full set of possible translation voxels, meaning translation voxels can contain between zero to forty-two cognate ternary complexes for a single chosen ribosome (distributed as in **Figure S4**). For translation voxels containing between one to forty-two cognate ternary complexes belonging to cells growing at each of the six growth rates, we simulated 100 replicates starting from different random initial conditions (42 × 6 × 100 = 25,200 total simulations). We set conditions for each replicate using the Mersenne Twister random number generator with seeds set as multiples of five (i.e., 0, 5, 10, …, 495). Simulations were terminated when the ribosome being tracked successfully reacted with a cognate ternary complex.

#### VI.1. Post-simulation analysis of statistically representative translation voxel ensembles

For each translation voxel simulation (*i*), we computed the elongation latency (*τ*_elong,*i*_), transport latency (*τ*_transport,*i*_), and reaction latency (*τ*_rxn,*i*_) of the cognate ternary complex that successfully reacted with the ribosome being tracked (StatisticallyRepresentativeTranslationVoxelAnalysis.ipynb). We incorporated the impact of near-cognate ternary complexes by scaling the time taken by a statistically accurate portion of non-cognate reactions at the tracked ribosome. Specifically, we leveraged our calculation that translation voxels have eight near-cognate ternary complexes and 32 non-cognate ternary complexes, on average (**Figure S4**), and that near-cognates have an average latency of 4.6 ms while non-cognates have an average latency of 1.4 ms (**Figure S2**), to randomly scale non-cognate reaction times 3.3-fold with 20% probability. We note that this representation of near-cognates does not capture the impact of near-cognate ternary complexes on other ribosomes in the voxel; near-cognates could slightly reduce overall latencies by occupying mismatching ribosomes for longer than noncognate ternary complexes, allowing cognate ternary complexes to find their match more quickly.

We subsequently computed an overall transport latency (*τ*_transport_), reaction latency (*τ*_rxn_), and elongation latency (*τ*_elong_) for each growth rate. We did so by calculating weighted averages of each of transport latency, reaction latency, and elongation latency acquired from translation voxel simulations for each particular growth rate (*μ*), averaging over replicates:

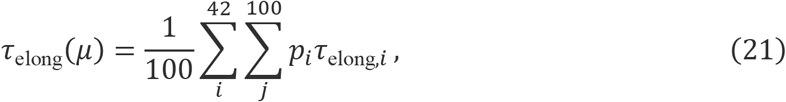

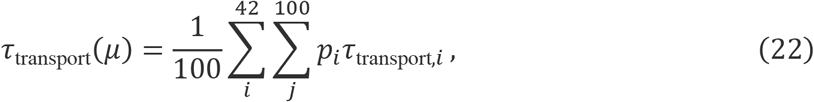

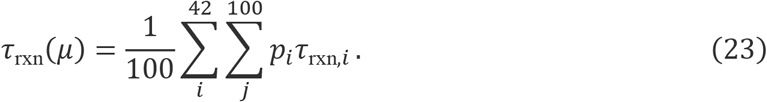

The probability of each translation voxel configuration (*p*_*i*_) is conditional on both the number of cognate ternary complexes in the particular translation voxel as well as growth rate (**Figure S4**). We did not consider the latency of translation voxels that contained zero cognate ternary complexes (~22% of translation voxel instances), since such voxels would have infinite latency and are an artifact of constraining translation voxels to 42 total ternary complexes. In particular, if larger voxels with more than 42 ternary complexes are considered, the resulting proportion of cognate ternary complexes is similar but with fewer instances of zero cognates (e.g., we found that voxels with 42, 84 or 168 ternary complexes have the same number of average cognates when normalized by number of total ternary complexes, but have 22%, 10%, and 3% of instances with zero cognates respectively). Not considering the zero cognate ternary complex voxels thus provides a lower bound estimate of elongation, transport, and reaction latency.

### VII. Event-based stochastic simulations of statistically representative translation voxel ensembles

To measure the effect of removing transport physics from our simulations (Supplement), we developed an event-based stochastic simulation algorithm of statistically representative translation voxel ensembles (EventBasedStochasticSimulation.py). As in our other simulations, the ensemble of voxels captures the relative abundances of ternary complexes and frequencies of codons among mRNA, but unlike our other simulations, physical space is not represented.

In our stochastic simulation algorithm, all ribosomes in translation voxels are initialized as bound to randomly chosen ternary complexes. Each reacting ternary complex-ribosome pair is then assigned a time until either disassociation or successful amino acid incorporation, drawn from the distribution of non-cognate, near-cognate, and cognate reaction latencies we computed as in Methods Section III.2. (**Figure S2**). The simulation proceeds in an event-based fashion, iteratively transitioning to the next event that occurs in the translation voxel (i.e., the timestep of simulation is not fixed). Following a disassociation event, the disassociated ternary complex joins the available (unbound) ternary complex population, and the newly available ribosome instantly binds to a randomly chosen ternary complex. The simulation ends when a cognate ternary complex successfully reacts with a matching ribosome.

We computed the elongation latency at particular growth rates by simulating the statistical distribution of possible translation voxels (i.e., with the full permissible range of cognate ternary complexes, distributed as in **Figure S4**) and then averaging their resulting elongation latencies. For each growth rate and permissible number of cognate ternary complexes, we simulated 5000 replicate translation voxels with different random initial conditions. Conditions were set for each replicate using the Mersenne Twister random number generator with seeds set as multiples of five (i.e., 0, 5, 10, …, 24995).

### VIII. Computational tools and costs

All fixed time-step simulations of translation voxels were performed using Colloidal Smoldyn (based on Smoldyn v2.61) deployed on Amazon Web Services. Simulations required ~300,000 CPU-hours in total. Our longest simulations, for translation voxels at a growth rate of 0.6 dbl/hr, took up to ~3 weeks for some replicates, while our shortest simulations took seconds. The cost of all our simulations was approximately US $10,000. Output file sizes for most simulation runs were small (<1 MB). All measurements and validation with LAMMPS were performed using the Sherlock High Performance Computing Cluster at Stanford University. Modeling, analysis, and event-based simulations were performed using Python 3.7.

### IX. Acceleration of translation voxel simulations to reduce runtime and cost

Simulations of translation voxels were originally forecast to cost US $6 million with the longest simulations taking ~36 years, making them intractable. To achieve feasible costs and run times, we implemented a procedure for accelerating our fixed-timestep simulations ~600-fold, reducing costs and run times as detailed above. In our acceleration procedure, kinetic rates of unbinding and codon recognition (i.e., the possible exits to the initially bound state) are increased 600-fold during simulations (*k*_1_ = 717 s^−1^ to 430200 s^−1^, and *k*_2*f*_ = 1474 s^−1^ to 884400 s^−1^). Simulations are run until completion following a successful match between a cognate ternary complex and matching ribosome. Subsequently, during post-simulation analysis, the time spent by ternary complexes in the initially bound state is re-scaled to be 600-fold slower, and re-scaled times are used to compute reaction, transport, and elongation latencies. Reaction latency is calculated as the time the cognate ternary complex spends bound in reactions; elongation latency is calculated as the total time the matching ribosome spends unbound or bound in reactions; and transport latency is calculated as the difference between elongation latency and reaction latency.

Our estimates of overall reaction, transport, and elongation latencies are not sensitive to this scaling procedure at the ~600-fold acceleration used (**Figure S5A-C**). This insensitivity is a result of unbinding kinetics remaining slow enough that, for a certain range of kinetic scaling, ternary complexes mix within the translation voxel in between unbinding events to a sufficiently similar extent (**Figure S5D**).

### X. Calculation of long-time self-diffusivity

We tracked the motion of individual biomolecules as they wandered far from their original positions, executing a random walk through the cytoplasm. This sampling of many configurations in a voxel is termed the long-time self-diffusion 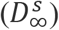 (referred to as diffusivity in the Results) and is a monotonically decreasing function of volume fraction at fixed molecule size polydispersity. We computed the long-time self-diffusion of particular biomolecule species (denoted by subscript *i*) at different growth rates by tracking the absolute position of biomolecules and computing their mean squared displacement over time (Results section IV):

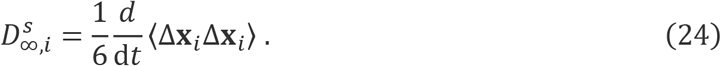

Here, the angle brackets signify an ensemble average over the motion of every biomolecule of a given species in a translation voxel.

### XI. Calculation of viscosity

We calculated the viscosity of translation voxels at different growth rates (Results section IV) by performing shear rheology simulations in LAMMPS. For each growth rate, we initialized suspensions representative of multiple contiguous translation voxels. We imposed a simple shear flow on the suspensions at a constant shear rate in the x-direction 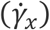 and measured the resulting interparticle stress 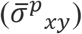. The shear rate imposed was chosen to be small enough to remain in the linear-response regime (i.e., with insignificant deformation), allowing measurement of the intrinsic or so-called zero-shear viscosity (*μ*_0_) (ViscosityCalculation.py) (Batchelor, 1977):

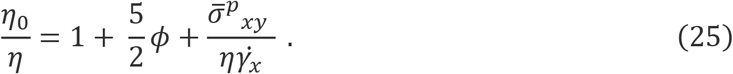

Here, the first two terms on the right-hand side of the equation are the Einstein viscosity that approximate the hydrodynamic contribution of particles to viscosity at equilibrium. The third term describes the interparticle contribution to viscosity and is equivalent to the Green-Kubo equilibrium interparticle contribution at the small shear rates used here. 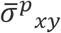 is computed as the *xy*-component of the interparticle stress 〈***xF^p^***〉, where ***x*** corresponds to the position vectors of the particles and ***F^p^*** is the (negative of) the gradient of a nearly hard-sphere, spherically symmetric repulsive potential.

### XII. Calculation of molecular proximity

We computed the proximity between ternary complexes and ribosomes at different growth rates (Results section IV). For each growth rate, we initialized 100 translation voxels with random initial spatial configurations, chosen using a Mersenne Twister random number generator with seeds set at multiples of five (i.e., 0, 5, 10, …, 495). Following a brief equilibration period, we measured the distance from each ternary complex to its closest ribosome. Our reported values of proximity for any particular growth rate are an average of the minimal distance for all ternary complexes across all corresponding 100 replicate translation voxels.

### XIII. Calculation of repeat reactions

We computed the average number of repeat reactions between ternary complexes and ribosomes at different growth rates (Results section IV). For each growth rate, we initialized 100 translation voxels with random initial spatial configurations, chosen using a Mersenne Twister random number generator with seeds set at multiples of five (i.e., 0, 5, 10, …, 495). We subsequently tracked the number of times a ternary complex consecutively re-reacted with the same ribosome following a mismatching reaction within each translation voxel. Our reported values for repeat reactions for any particular growth rate are an average across all corresponding 100 replicates for the given growth rate.

### XIV. Chemical kinetics sensitivity analysis

To measure the sensitivity of our predicted elongation latencies to changes in chemical kinetics, we simulated the impact of slowing down or speeding up intra-ribosomal kinetic rates on elongation latency. To do so, we simulated ensembles of translation voxels as in Methods section VI while varying kinetic rates individually or all together and measuring the resulting elongation latency (**Figure S9**). Since the kinetic rate *k*_1r_ impacts the mix time of voxels (**Figure S5**), we varied the level of kinetic acceleration in our simulations for different *k*_1r_, ensuring that our translation voxels were simulated in regimes in which elongation latency is insensitive to changes in kinetic acceleration (our kinetic acceleration scheme is described in Methods section IX).

### XV. Calculation of maximum packing and projected growth rate voxel parameters

We computed the theoretical maximum packing for translation voxels between observed growth rates, μ = 0.6 dbl/hr to 3.0 dbl/hr, as well as faster hypothetical growth rates, μ=3.0 dbl/hr to 8.0 dbl/hr, using the theoretical calculations of maximum packing for tridisperse systems from Farr and Groot, 2009. Our translation voxels are comprised of molecules having a 1:3:6.5 size ratio, which differs from the particle size ratio used by Far and Groot (1:3:9), giving an overprediction of our computed maximum packing of less than 10%.

To estimate ribosome abundances, ternary complex abundances, cell mass, cell volume, and nucleoid volume at hypothetical growth rates between 3.0 dbl/hr and 8.0 dbl/hr, we extrapolated from observed growth rates **(Figure S1)**, guided by observed trends below 3.0 dbl/hr and allowing uncertainty while rejecting unphysical projections (e.g., negative cell mass and nucleoid volume fraction) (**Figure S11, Figure 6)**. We calculated bounds by perturbing the extrapolated fits while still maintaining all expected trends (e.g., the lower bound of ribosome abundances never decreases with increasing growth rate). We computed volume fractions for translation voxels at hypothetical growth rates as in Methods Section II, setting upper and lower bounds by considering all permutations of fits for ribosome abundances, ternary complex abundances, cell mass, cell volume, and nucleoid volume fraction (ProjectedGrowthRateCalculations.ipynb).

## Acknowledgements

We thank Jennifer Hofmann for her input on transport physics in cells; Jonathan Calles for discussions surrounding systems biology, biophysics, and biochemistry; Dr. Steve Andrews for his assistance with using Smoldyn; and Anton Jackson-Smith, Rolando Perez, and other members of the Endy and Zia labs for helpful conversations and feedback. This work was supported by a National Institutes of Health T32 Training Grant No. GM007365 for A.J.M., a Stanford Bio-X Graduate Fellowship for E.G., and a National Science Foundation Graduate Research Fellowship under Grant No. – 1656518 as well as a Stanford Graduate Fellowship for A.M.S. Additional support was provided by the National Institute of Standards and Technology (70NANB15H268).

## Supplemental Information

## Appendix A. Colloidal Smoldyn theoretical framework and simulation implementation

Here, we briefly describe the theoretical framework that underlies Colloidal Smoldyn and provide simulation implementation details for transport and reaction dynamics.

### Simulating colloidal transport dynamics

Here, we briefly describe the theoretical framework that underlies Colloidal Smoldyn and then provide simulation implementation details. The transport dynamics of water-suspended particles of size ~ 1.5 nm to O(10) microns (colloids) have been thoroughly studied and characterized in the microhydrodynamics and colloid physics literature. Fluid motion generated by particle motion at these length scales is governed by the Stokes equations, while particle motion is described by the N-particle Langevin equation (Langevin, 1908),

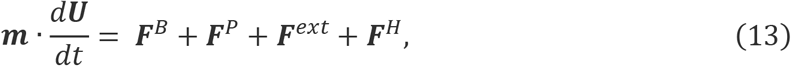

a stochastic force balance, where particle momentum ***m*** · *d**U**/dt* equates the sum of the forces ***F*** that act on the particles. Here ***m*** is a diagonal tensor containing the details of the mass of the particles and ***U*** is the particles’ velocity vector. The forces ***F*** represent the stochastic forces ***F**^B^* that give rise to Brownian motion, the deterministic interparticle forces ***F**^P^* that here arise from a hard-sphere potential, the external forces ***F**^ext^* that could represent either active motion or externally induced motion, and hydrodynamic drag forces ***F**^H^* that arise due to the difference between the particle velocity ***U*** and imposed fluid motion ***u***^∞^. Each of the vectors (***U*** and ***F***) has dimensions 3N describing the three-dimensional space in which all N particles move. In our system there are no external forces on the particles, thus ***F**^ext^* = 0.

The Brownian force 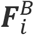 on a given particle “*i*” arises from collisions between the colloidal particle and the solvent molecules as the fluid fluctuates thermally and follows Gaussian statistics,

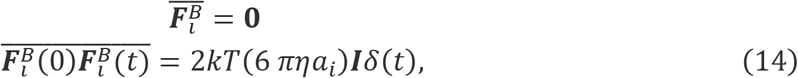

where *k* is the Boltzmann constant, *T* is the absolute temperature, *η* is the solvent viscosity,*a_i_* is the particle radius, ***I*** is the identity tensor, and *δ*(*t*) is the Dirac delta function. The overbar considers averages over times long compared to the solvent molecule timescale, such that a colloidal particle has undergone many random and decorrelated impacts from solvent molecules. Although the mean Brownian force is zero, its covariance, 2*kT*(6 *πηa_i_*)***I*** has an amplitude set by the fluctuation-dissipation theorem, which dictates that colloidal motion due to thermal forces is dissipated viscously back into the solvent. In our system, we approximated the drag force on the colloid, (6 *πηa_i_*)***I***, to be constant, where each particle experiences solvent drag and excludes other particles entropically with finite size, but coupled hydrodynamic interactions are neglected. This common simplification, the “freely draining model” is appropriate for gaining understanding of the basic colloidal physics of a suspension (Hoh and Zia, 2016a, 2016b; Russel, 1984; Russel et al., 1989; Zia and Brady, 2010) and is accurate in many physiological conditions. Incorporating hydrodynamic interactions in such models is routine (Banchio and Brady, 2003; Durlofsky et al., 1987; Ouaknin et al., 2021; Sierou and Brady, 2001) although computationally expensive and will be discussed in future work.

Interactions between colloidal particles can be attractive or repulsive and range from short- to long interparticle distances. In the present work, biomolecules are represented as hard spheres: they interact only at contact at which point we stipulate that they may not overlap. This can be represented mathematically via a hard-sphere potential that enforces an infinite penalty for overlap. The gradient of this entropic exclusion gives rise to a deterministic interparticle force between two particles of radius *a_i_* and *a_j_*

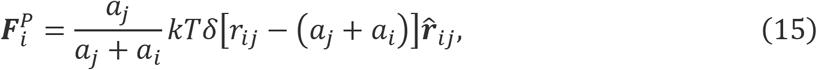

where *r_ij_* is the interparticle distance between the particles’ centers, 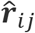 is the unit vector along the line that connects the particle centers, and the delta function *δ* enforces the infinite penalty for overlap.

Finally, the hydrodynamic drag forces are given by the Stokes drag on each particle according to

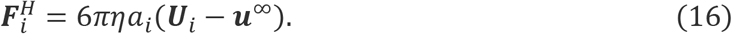

Overall, the fluid motion is governed by Stokes’ equations; hence, particle dynamics operate in the “overdamped limit” of the Langevin equation, where integrating twice over a time interval Δ*t* large enough for a colloid to have received many uncorrelated solvent molecule impacts and to have relaxed their momenta (Δ*t* ≫ *m*/6*πηa_i_*) results in the displacement equation (see Ermak and McCammon, 1977 for further details)

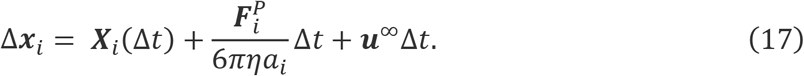

Here, Δ***x**_i_* is the change in position over a time step Δ*t*, owing to the combined contributions of the Brownian force, the interparticle potential, and the imposed flow. In **Equation 17**, the Brownian displacement ***X**_i_* (Δ*t*) obeys Gaussian statistics given by

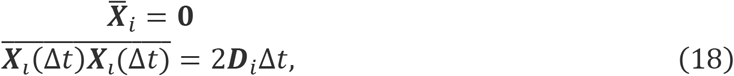

where the amplitude of the co-variance in this freely draining model is set by the diffusion coefficient of a single particle defined by the familiar Stokes-Einstein relation, ***D**_i_* = *kT*/6*πηa_i_**I*** (Einstein, 1905; Stokes, 1850). In other approaches, the diffusion coefficient used to compute Brownian displacements is sometimes obtained from *in vivo* measurements in order to implicitly encode the effects of crowding but limits the applicability of the model to a specific temperature or crowding condition of the cell (among other issues). In our approach, the effects of crowding on particle motion emerge naturally from the excluded volume interactions enforced by the interparticle force (**Equation 15**): each particle undergoes a Brownian displacement commensurate with **Equation 18**, reflecting the dissipation of a thermal kick via its own solvent drag. However, over many displacements, a particle will encounter other particles and must “wait” for them to move in order to sample that space, leading naturally to a slow-down of diffusion over longer distances through a crowded suspension. Changes in volume fraction thus seamlessly lead to changes in particle dynamics and interactions.

The hard-sphere interparticle force (**Equation 15**) imposes a singular force between particles at contact. The modeling of singular forces in dynamic simulation is intractable; at best, very steep repulsions can be captured. To overcome this limitation, rather than to impose a singularity, we utilized a “potential-free” algorithm (Foss and Brady, 2000; Heyes and Melrose, 1993), where once Brownian and imposed flow displacements are made, particles are permitted tiny overlaps which are then corrected. A pair of particles do not interact at all unless they come into direct contact or overlap and once they do interact, the hard-sphere displacement of each corrects any overlap:

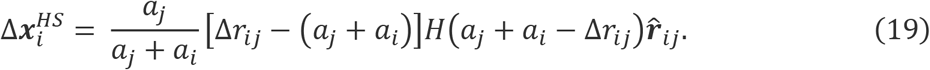

Here, Δ*r_ij_*–(*a_j_*+*a_i_*) is the overlap being corrected, the Heaviside step function *H* ensures that particles are displaced only when they overlap, and 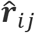 is the unit vector along the line that connects the particle centers, giving the hard-sphere displacement along the line of centers. This approach is equivalent to applying to each particle an interparticle force 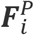 that is proportional to the overlap to ensure entropic exclusion. The physical consequence of the hard-sphere interaction and the overlap correction is a contribution to the osmotic pressure that arises from the finite volume particles occupy in the suspension (Brady, 1993).

Incorporating in **Equation 17** the potential-free algorithm and considering that in our system there are no imposed flows ***u***^∞^ gives a displacement equation of the form

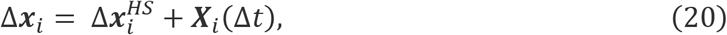

where 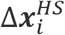 is the displacement that enforces the hard-sphere potential. **Equations 18 - 20** constitute the framework underlying dynamic simulations in Colloidal Smoldyn, our adaptation of the original Smoldyn.

In our simulations, we chose a timestep Δ*t* = 62 picoseconds empirically by testing the temporal resolution necessary to recover expected mean-squared displacements over long timescales, which we computed using the open-source LAMMPS Molecular Dynamics Simulator, across the levels of crowding we modeled (**Figure S3**) (Plimpton, 1995). Practically, this timestep corresponds to an average Brownian displacement per-timestep of 0.1 nm for ribosomes, 0.14 nm for ternary complexes, and 0.25 nm for proteins (**Equation 18, Table S4**).

At each time step all particles have a Brownian displacement, after which the algorithm checks for overlaps between particles. In Colloidal Smoldyn, overlapping biomolecules are separated along their line of centers and placed at contact according to Δ***x**^HS^* (**Equation 19**) and in accordance with well-established and experimentally validated models of colloidal physics (Heyes and Melrose, 1993). Our implementation is a direct modification to the physics represented by the original Smoldyn algorithm; the prior method over-corrects the overlap by separating the pair’s surfaces by the full distance, meaning a gap is introduced almost as though the collision was being modeled as inertial (Andrews, 2017). In addition, the particles were moved not along their line of centers in the original algorithm but rather at a reflected angle again reminiscent of an inertial collision which is not appropriate for colloids in the overdamped dynamics limit. The original method of Smoldyn, although intuitive, is physically incorrect for systems in which fluid motion is governed by Stokes equations, and a strong drag force is exerted on the approaching molecules. In such systems inertia is irrelevant, meaning that when two particles approach after receiving a Brownian kick they will simply slow down until another Brownian displacement sets them in a different direction. Placing particles at contact along the line of centers upon overlap approximates this entropic exclusion interaction and the interparticle forces associated with this displacement contributes to the osmotic pressure of the suspension. The Heyes and Melrose algorithm has been shown to recover suspension osmotic pressure as predicted by statistical mechanics models (Foss and Brady, 2000; Zia and Brady, 2012).

In simulations of crowded systems, correcting a set of overlaps typically produces additional new overlaps, necessitating additional iterations of overlap correction. Two strategies are applied to accurately process overlaps: first, utilizing short, discretized time steps to generate smaller and fewer overlaps; second, executing several overlap-correction cycles. For computational efficiency, the original Smoldyn algorithm executed only one round of overlaps, leaving many still-overlapping particles. Only in dilute systems is such an approximation physically accurate. In colloidal Smoldyn, we conducted an optimization study to identify an appropriate balance of efficiency and accuracy, from which we devised an algorithm with three iterations of overlap resolution at the chosen timestep of Δ*t* = 62 picoseconds.

In all our simulations, we avoided finite-size and boundary effects by implementing periodic boundary conditions. The voxel sizes are large enough to avoid finite-size effects; we show that voxels produce accurate physical structure and dynamics, by showing that the pair-correlation function for all pairs decays well within the voxel (**Figure S15**). As further evidence, the long-time self-diffusivity measured in LAMMPS was done with a system 100 times larger (in volume); agreement with our results in Smoldyn is excellent (**Figure S3)**.

### Simulating reaction dynamics

In our simulations, after a ternary complex and unbound ribosome interact both biomolecules are converted to form a single bound-ribosome that either disassociates or forms a peptide bond following single-molecule rate kinetics. A bound ribosome has the same radius as an unbound ribosome (an approximation of bound ribosomes internalizing tRNA during codon recognition). In the native Smoldyn algorithm, the distance at which chemical interactions are initiated between molecules is set *a priori* to be different than contact (e.g., molecules may be set to only react when halfway overlapping or conversely while still separated by several molecule diameters). In particular, the initiating distance is computed such that the overall reaction rate of all molecules in simulation is equal to experimentally-measured kinetic rates set for individual reactions (Andrews and Bray, 2004). This approximation is based on the Smoluchowski model for reactions and is only reasonable for dilute systems (or systems of point particles). To achieve a more physical representation for crowded colloidal systems (as in our modeling of transport) that allows ab initio estimates of latency without fitting, we further modified the Smoldyn algorithm in Colloidal Smoldyn to only allow for initiation of chemical reactions upon the physical (entropic) interaction of reactant biomolecules. We did not consider factors such as binding site orientation that would restrict reactions to only certain interactions or reaction activation energies that may need to be overcome prior to reaction. Instead, we made the approximation that all physical interactions between ternary complexes and ribosomes (i.e., overlaps between ternary complexes and unbound ribosomes at the end of a diffusive step) lead to reactions. Ribosomes and ternary complexes that disassociate following a reaction are placed at contact, congruent to the Brownian dynamics underlying our modeling of transport. This was a final modification to the Smoldyn algorithm, which places biomolecules at a distance following disassociation to artificially reduce repeat reaction probability based on fitting of experimental single molecule reaction kinetic parameters (an approximation that, as before, is reasonable only for dilute systems) (Andrews and Bray, 2004).

We computed the time a ternary complex spends bound to a ribosome via a Markov process consisting of well-defined intra-ribosomal states and transition kinetic rates (*k_i_*) (Kinz-Thompson et al., 2016). Specifically, the algorithm we used to model biochemical reactions involving ternary complex-bound ribosomes is:

1. For each state transition *i* → *j* out of the current state *i*, draw a transition dwell time,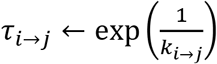;
2. Transition to state *j* corresponding to the fastest transition time, argmin_*i*_(*τ_i→j_*);
3. Draw an updated dwell time for the transition to state *j*, 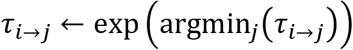;
4. Add the updated dwell time, *τ_i→j_*, to total dwell time *τ*_tot_;
5. Repeat until either the ternary complex dissociates from the ribosome or a peptide bond is successfully formed.

Using our algorithm and reported kinetic rates, we computed the distributions of latencies for non-cognate, near-cognate, and cognate ternary complexes reacting with ribosomes (**Figure S2, Table S5**, ReactionKineticsCalculations.ipynb). We did not consider the unlikely events of incorrect reactions leading to mis-incorporated amino acids (≤ 1%) or cognate ternary complexes being rejected following successful codon recognition (0.04%). The overall processing order of our algorithm is: first diffusive transport of molecules, then reactions between overlapping molecules, and then overlap resolution.

#### Note S1. Sensitivity analysis and the impact of parameter values

We sought to understand if our prediction that the productivity of individual ribosomes increases due to stoichiometric crowding is sensitive to the fact that our unfitted model does not exactly match observations of absolute elongation latencies. One natural place to start is to make a change in the values of the chemical kinetic parameters used in our model, which are taken from *in vitro* measurements (**Table S5**). The rationale for modulating these parameters is that *in vitro* kinetic rates may differ from their *in vivo* values due to, for example, differences in salt concentrations *in vitro* compared to *in vivo* (**Figure S13A**). To test this idea, we implemented a uniform three-fold increase of all intra-ribosomal chemical kinetic rates in our model (**Figure S13C**). We then simulated ensembles of translation voxels at varying growth rates and measured the resulting elongation rates. This chemical-parameter change closed the quantitative gap: elongation latency (*τ*_elong_ = 100 ms at 0.6 dbl/hr to 45 ms at 3.0 dbl/hr) now matched experimental latency (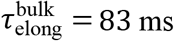 at 0.6 dbl/hr to 48 ms at 3.0 dbl/hr) (**Figure S13B**). Importantly, the speedup in elongation as growth quickens occurs even with the higher chemical kinetic rates used to fit the bulk experimental observations.

We also sought to test if our predicted increase in ribosome productivity requires physical transport. To do so we stress-tested the sensitivity of elongation to changes in transport by modeling instantaneous transport within our voxel ensembles (**Methods**). We found that the growth-rate dependence of elongation latency is lost when transport is modeled as an instantaneous process (**triangles in Figure S13B, Figures S6, S7**), demonstrating the essential role of transport physics in the growth-rate dependence of elongation rate, and also demonstrating the criticality of accurate chemical parameters in establishing absolute quantitative prediction of elongation latency.

#### Note S2. Translation initiation latency relative to translation elongation latency

Translation initiation is slow compared to a single elongation step. Therefrom, one might naively expect initiation to be rate limiting to protein synthesis overall and, thus, expect that any increase in protein synthesis rates should mostly arise due to increases in the rate of initiation.

However, clues to the contrary are apparent. For example, there is overwhelming experimental evidence from work to optimize expression of heterologous proteins (Welch et al., 2009). Specifically, synonymous changes in coding sequences that require ribosomes to make the same protein via coding-sequence unique elongation processes produce dramatic changes in protein synthesis rates and abundances. That is, codon usage changes alone can result in changes in gene expression levels, ranging from undetectable to majority of cell protein. This strong impact on protein synthesis rate is solely due to impacts on translation elongation, not translation initiation. Studies of natural living systems further reveal a literature that advances how both translation initiation and elongation can limit protein synthesis rates, depending on conditions (Subramaniam et al., 2014; Vieira et al., 2016).

Nevertheless, it is important to consider initiation more formally and explain why elongation is most likely to underlie increases in overall protein synthesis rates. To this end, we return to the conventional view of translation elongation in which the correct ternary complex instantaneously presents itself to the ribosome exactly when needed (i.e., they appear instantaneously in exactly the right order). That is, a situation in which there is no additional latency due to physical transport, nor combinatorial sampling by and among competing ternary complexes (i.e., setting aside the fact that our work shows that most elongation latency arises from transport, not kinetics). In these unrealizable conditions, the rate of peptide bond formation is entirely determined by chemical kinetics and is about 42 ms per amino acid. By comparison, translation initiation should be much slower, taking about 1-12 seconds. But protein synthesis requires not just one elongation step but hundreds in series (i.e., the average protein in *E. coli* is ~333 amino acids). Considering elongation of the entire protein, we estimate ~14 seconds total just from chemical kinetic latency alone (42 ms per amino acid). Thus, it becomes apparent that, while the chemical kinetics of a single initiation event might be relatively slow, most of the time is spent in the elongation phase (e.g., 12 seconds for initiation vs. at least 14 seconds for elongation).

#### Note S3. Estimating the evolutionary impact of increasing ribosome productivity

To explore the potential evolutionary impact of increasing ribosome productivity, we estimated how much slower bacterial growth would be if ribosome productivity did not increase (i.e., remained fixed) and only the abundances of translation machinery increased.

For a cell to replicate, it must replicate all its protein machinery. Since a fraction of peptide bonds formed during cell replication belong to translation machinery, as individual cells grow, increased translation machinery should enable more total peptide bonds to be formed per unit time. We can estimate the lower and upper bounds of total peptide bond formation in the time cells typically take to double, and thus growth rate, by assuming that this fraction is 0 (i.e., a lower bound in which no new peptide bonds enable more translation) and by assuming this fraction is 1 (i.e., an upper bound in which all new peptide bonds are incorporated into ribosomes that enable more translation).

As a lower bound estimate,

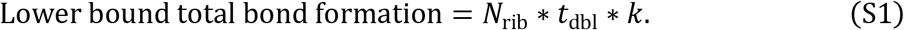

As an upper bound estimate,

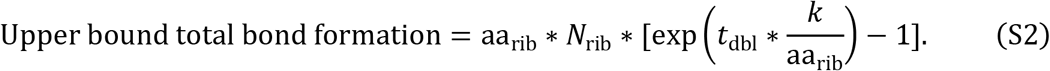

Here, *t*_dbl_ = μ/3600 is the time in seconds per cell doubling and aa_rib_ = 7336 amino acids/ribosome is the number of amino acids in a ribosome.

At μ = 3.0, there are on average *N*_rib_ = 62000 actively elongating ribosomes per cell elongating at a rate of *k* = 21 amino acids/s, leading to:

Lower bound total bond formation = 1,562,400,000 bonds, and
Upper bound total bond formation = 13,660,863,648 bonds.

As a sanity check, we can estimate the total bonds in a cell at μ = 3.0 dbl/hr from the total bonds composing proteins and total (active and non-active) ribosomes (**Table S1, S4**): 7.78 × 10^6^ proteins * 264 aa/average-sized protein + 73000 total ribosomes * aa_rib_ = 2,589,448,000 bonds, which lies between the lower and upper bounds as expected.

We can next consider the hypothetical case in which ribosome productivity at μ = 3.0 dbl/hr does not increase and instead remains fixed at that of μ = 0.6 dbl/hr, *k*’ = 12 amino acids/s. In this case, we can estimate the lower bound and upper bound time needed for total bond formation by solving for *t*_dbl_ in equations S1 and S2.

In the lower bound case,

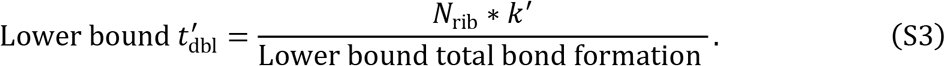

In the upper bound case,

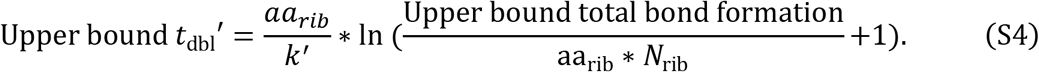

This gives us,

Lower bound *t*_dbl_′ = 2100 s, giving an effective μ’ = 1.7 dbl/hr
Upper bound *t*_dbl_′ = 2100 s, giving an effective μ’ = 1.7 dbl/hr

Thus, we estimate that maximum observed cell doubling would proceed at ~1.7 dbl/hr instead of 3.0 dbl/hr in the hypothetical case in which ribosome productivity did not increase and remained fixed at that of μ = 0.6 dbl/hr.

What would this difference mean practically from an evolutionary perspective? We can compare the expected growth of microbes with these different growth rates:

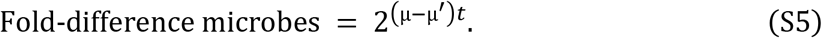

After *t* = 6 hours there would be ~200-fold more of the microbes with increased ribosome productivity, after 12 hours ~50,000-fold more, and after 24 hours 2.5 billion-fold more.

#### Note S4. On the relation of our modeling to other modeling methods

##### Mass-action kinetic models

Mass-action kinetics models of translation elongation such as that by Klumpp et al. abstract away space and stochasticity entirely, approximating diffusion by inserting a value for a diffusion coefficient into a mass balance. In contrast we explicitly represent space, modeling Brownian motion in our simulations directly from the Langevin equation and intra-ribosomal reactions using a stochastic process (see Methods).

Explicit consideration of spatial effects leads to significantly different system dynamics and thus conclusions. One conclusion of the Klumpp et al. model, for example, is that when diffusion is slower, reactions are slower and thus protein synthesis is slower. Yet, by considering spatial implications of well-established experimental data (Methods) (Dennis and Bremer, 2008; Volkmer and Heinemann, 2011), we find that *E. coli* becomes more crowded as it grows faster; by then examining detailed particle trajectories, which captures varied diffusion and transport times due to crowding, polydispersity, and spatial proximity, we find that closer proximity between translation molecules boosts protein synthesis rates even when diffusion slows down. Homogenized diffusion coefficients used in mass-action kinetics models, even adjusted for growth rate and crowding, smear out underlying particle displacements, and thus come to incorrect mechanistic conclusions in this case.

##### Gillespie simulations

A Gillespie simulation is a classic example of how to abstract away the impact of molecular transport, searching, and encounters on the rate of reaction while maintaining an approximation of stochasticity. A Gillespie model imports mass-action kinetics for reaction rates and then uses this information to create a distribution of events over time. In our model, we obtain a distribution of the time it takes for cognate or non-cognate ternary complexes to react with ribosomes once bound. But the rate at which such reactions start (and thus complete) depends on the probability of the cognate ternary complex encountering the ribosome. This is a statistically independent (Brownian) physical search process that is explicitly represented in our simulations, but which in the Gillespie algorithm would proceed instantly without any spatial consideration. After many reactions using many voxels, our simulations will produce data that will recover the smeared-out Gillespie simulation parameters; but our physical resolution reveals the mechanistic process by which the Gillespie parameters speed up.

#### Note S5. On the difference between mass density and volume fraction in cytoplasm

The mass density and volume fraction of cytoplasm are independent quantities. A simple illustration of this fact is that a kilo of lead (5cm x 5cm x 5cm) can easily fit inside a bag that would be too small for a kilo of cork (23cm x 23cm x 23cm). A similar effect is at play in *E. coli* cytoplasm as the cell adjusts its molecular composition across changes in growth rate. Volume fraction changes independent of changes in mass density of cytoplasm (Figure S17) because the mass density varies among individual biomolecules (e.g., proteins are five times denser than ternary complexes). Thus, as the relative molecular abundances change, the occupied volume fraction changes even if the overall mass density of the cell remains fixed (as illustrated in a simplified case in Figure S18). In turn, since occupied volume determines the diffusivity in cytoplasm, it is volume fraction rather than mass density that sets the diffusion coefficient involved in cellular processes.

**Figure S1.**
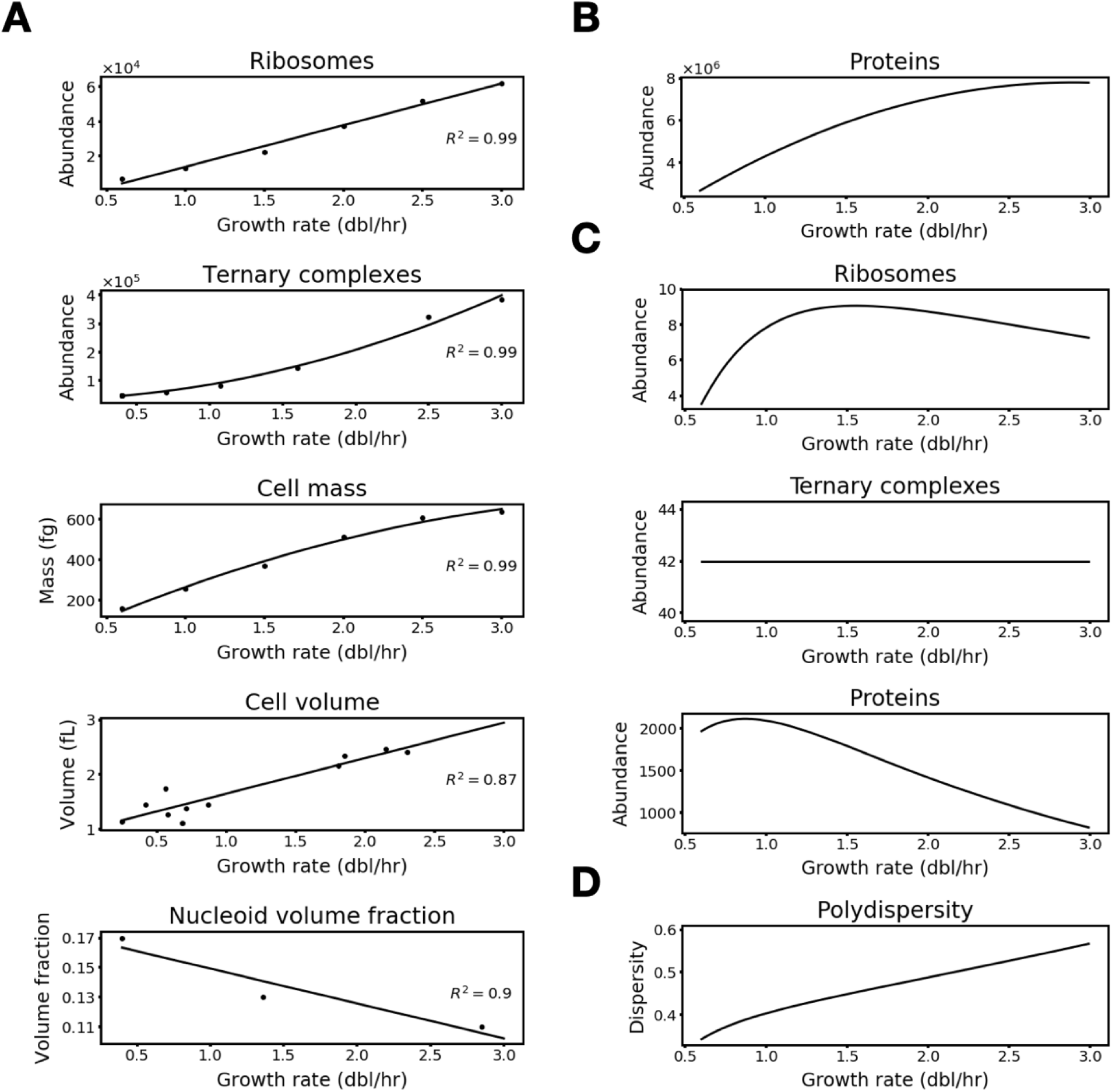
Cell and translation voxel parameters vary with growth rate. **(A)** Polynomial fits of reported cell parameter measurements (data points are taken from Tables S1, S2, and S3). (**B)** Estimate of protein abundances in cells across growth rates. (**C)** Estimate of ribosome, ternary complex, and protein abundances in translation voxels across growth rates. **(D)** Estimate of the polydispersity of translation voxels across growth rates.

**Figure S2.**
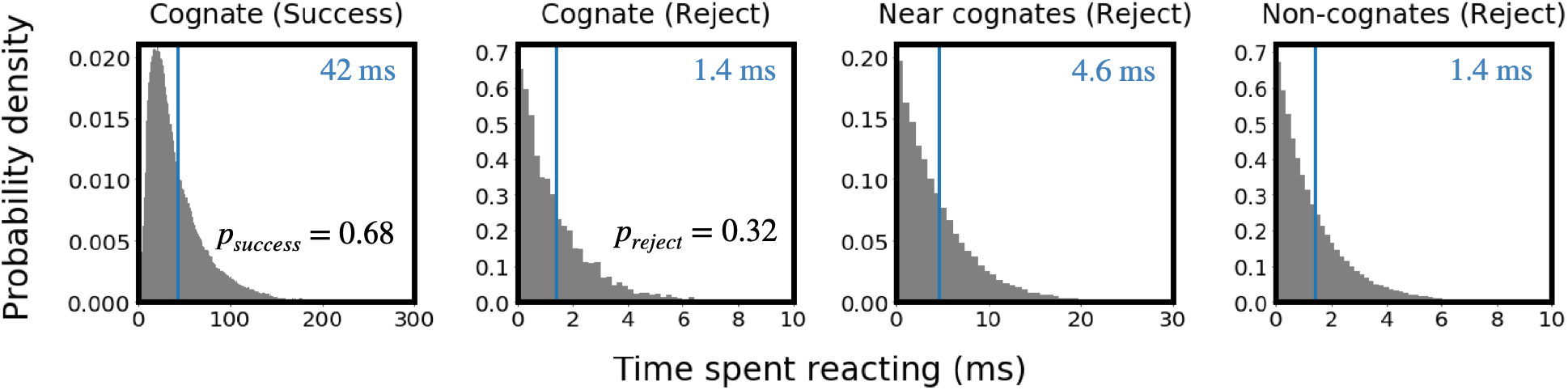
Ribosomal kinetics are dependent on whether a ternary complex is cognate, near-cognate, or non-cognate. **(A-D)**: Distribution of time spent by tRNA within ribosomes, computed by sampling tRNA-ribosome reaction trajectories. **(A)** Cognate ternary complexes that successfully react (probability shown). **(B)** Cognate ternary complexes that are rejected (probability shown). **(C)** Near-cognate ternary complexes that are rejected (only rejections are considered). **(D)** Non-cognate ternary complexes that are rejected (only rejections are considered). In each plot, average latency is marked by a blue line and displayed on the top right.

**Figure S3.**
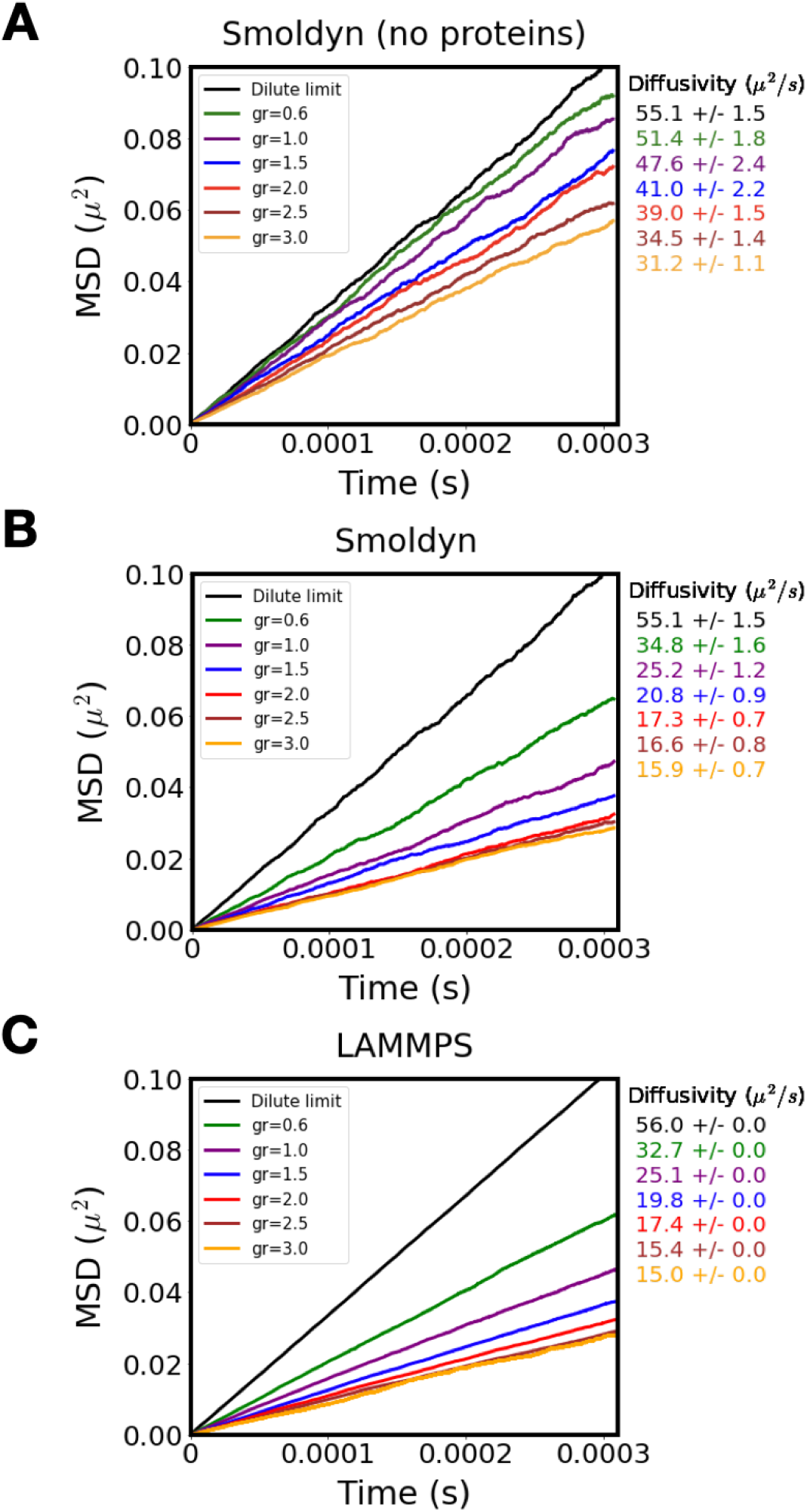
The long-time self-diffusivity of ternary complexes is dependent on crowding and growth rate. **(A)** Measurements of ternary complex mean squared displacement in translation voxels without proteins using Smoldyn. (**B)** Measurements of ternary complex mean squared displacement in complete translation voxels (i.e., with proteins) using Smoldyn. (**C)** Measurements of ternary complex mean squared displacement in complete translation voxels using LAMMPS. System size tested in LAMMPS is 100 times larger (in volume) than in Smoldyn to obtain better statistics and test finite size effects. Agreement with (B) and (C) and confirms the accuracy of our implementation in Smoldyn and that our voxels avoid finite size effects. For all plots, in the dilute limit, ternary complex long-time diffusivity should equal its short-time diffusivity (*D* = 56 μm^2^/s, Table S4).

**Figure S4.**
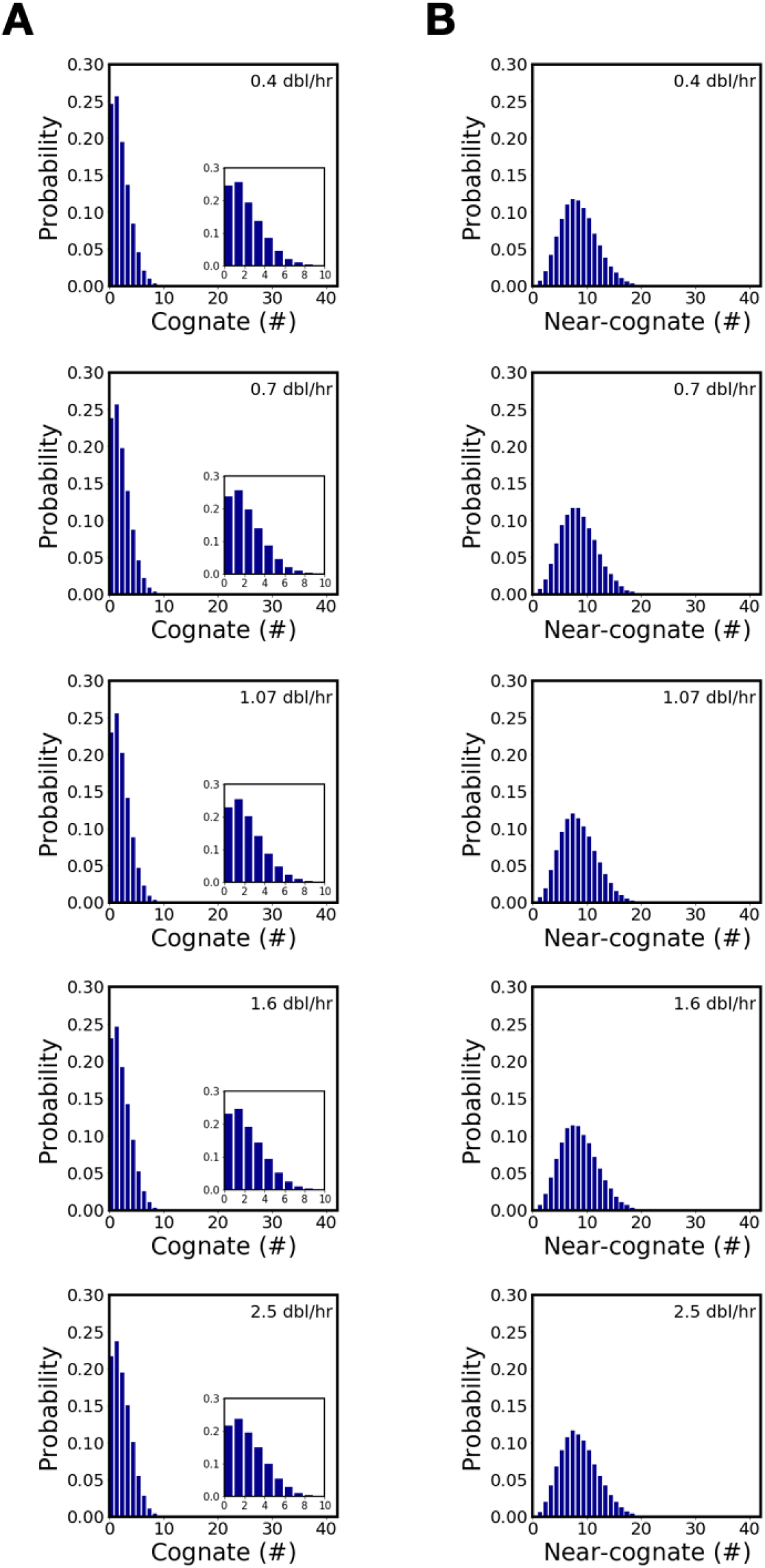
The number of cognate and near-cognate ternary complexes in a translation voxel follows a slightly growth-rate dependent probability distribution. **(A)** Distribution of cognate ternary complexes in translation voxels. (**B)** Distribution of near-cognate ternary complexes in translation voxels. Insets highlight the distribution between values of zero to ten cognate ternary complexes.

**Figure S5.**
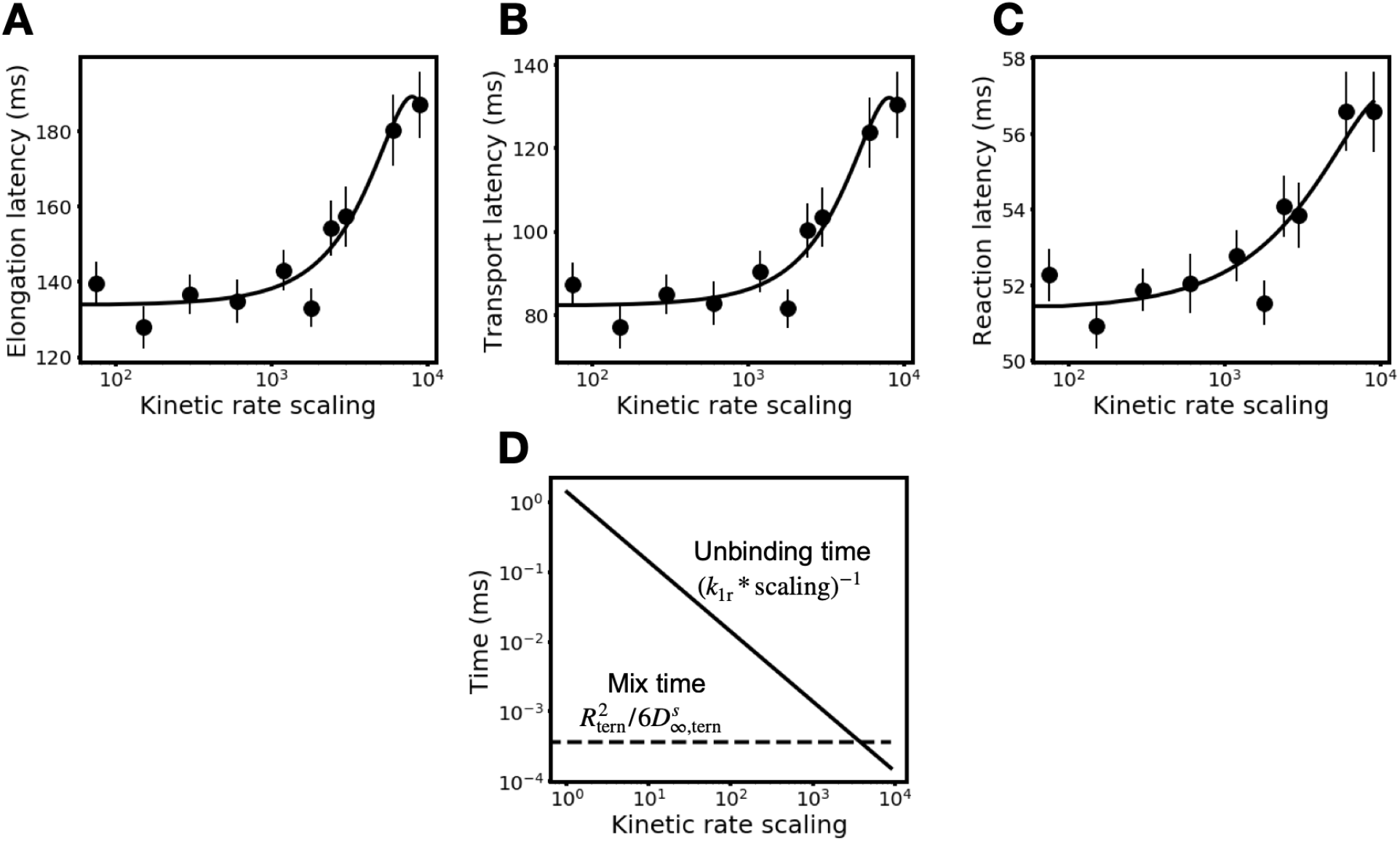
Predictions of translation latencies are insensitive to low and intermediate kinetic rate scaling due to differences in unbinding and mixing timescales. **(A-C)** Elongation latency, transport latency, and reaction latency are insensitive to kinetic rate scaling at a scaling of less than ~2000. Data shown for a translation voxel with growth rate μ = 3.0 dbl/hr. **(D)** The average time ternary complexes take to unbind from ribosomes (unbinding time) remains slower than the typical time ternary complexes take to diffuse their radius within a voxel (mix time, calculated using Equation 18 in the Methods). The mix time plotted is computed for the most crowded growth condition (*ϕ*_vox_= 0.42, μ = 3.0 dbl/hr).

**Figure S6.**
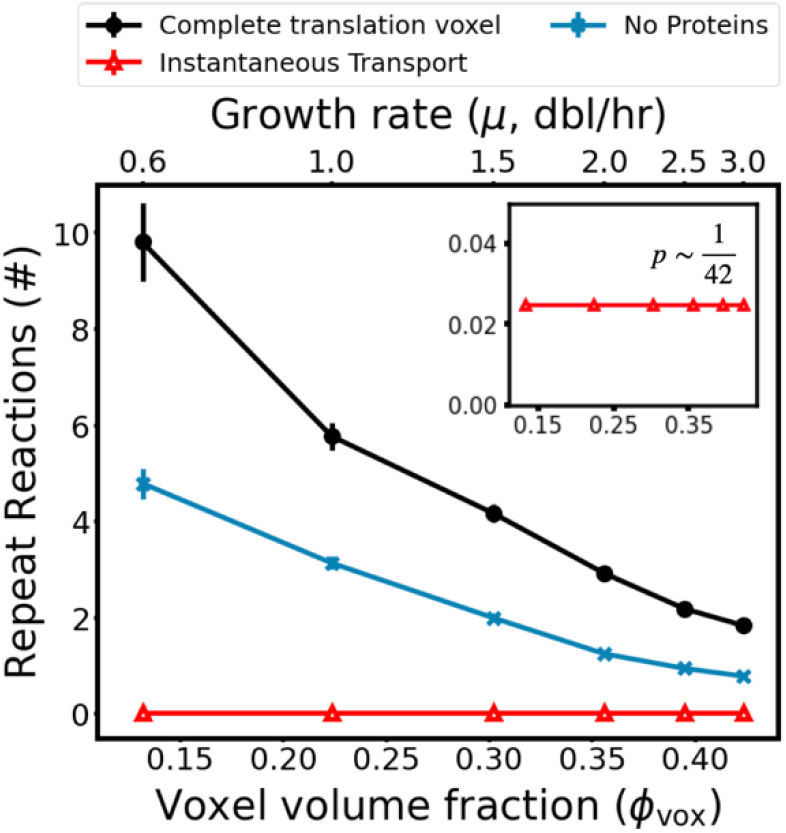
Repeat reactions between mismatching ternary complexes and ribosomes are sensitive to crowding and instantaneous transport. Average number of repeat reactions following initial reaction in complete translation voxels (black curve, filled circles), translation voxels with no proteins (blue curve, x’s), and translation voxels with instantaneous transport (red curve, triangles). Inset shows that simulations with instantaneous transport produce repeat reactions with uniform random likelihood across all growth rates.

**Figure S7.**
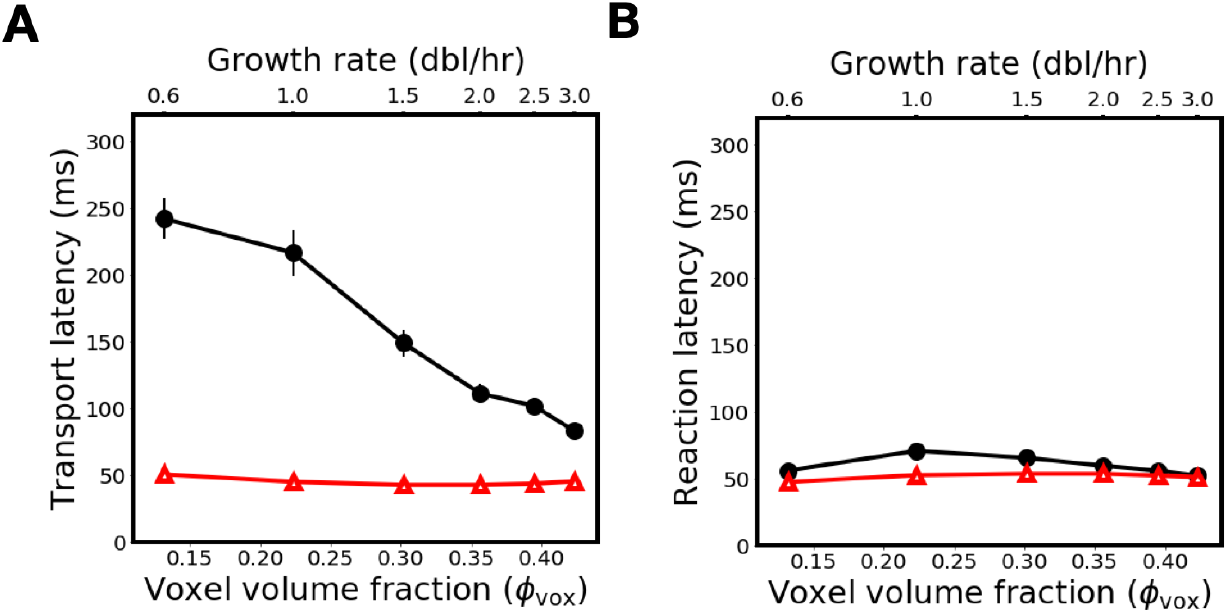
Growth rate dependence of transport and reaction latency is lost when transport is modeled as an instantaneous process. Simulations with instantaneous transport (red curve, triangles) produce **(A)** transport latencies and **(B)** reaction latencies that are uniform across growth rates. Original simulation results are shown for comparison (black curve).

**Figure S8.**
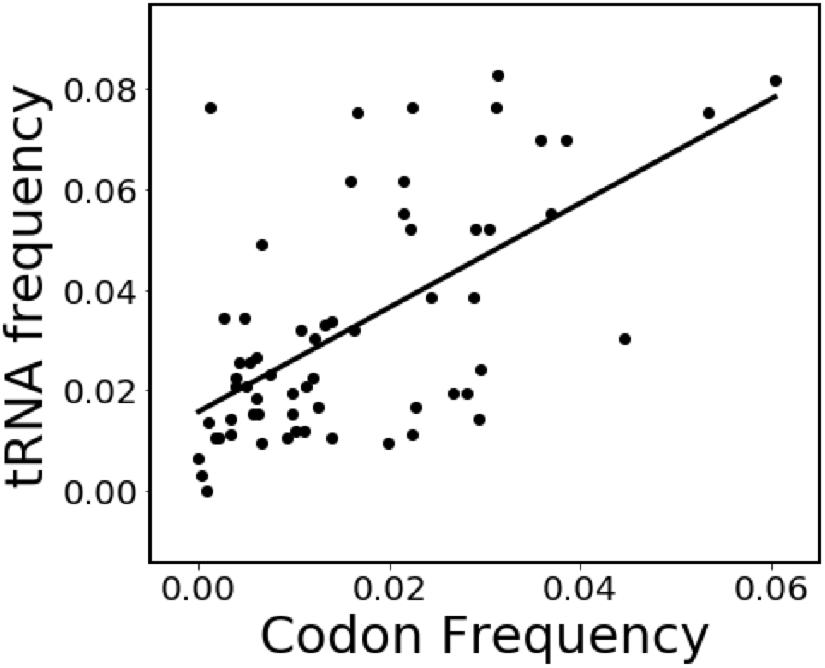
The frequency of a given codon and their corresponding tRNA are weakly correlated. Frequency data from Table S6.

**Figure S9.**
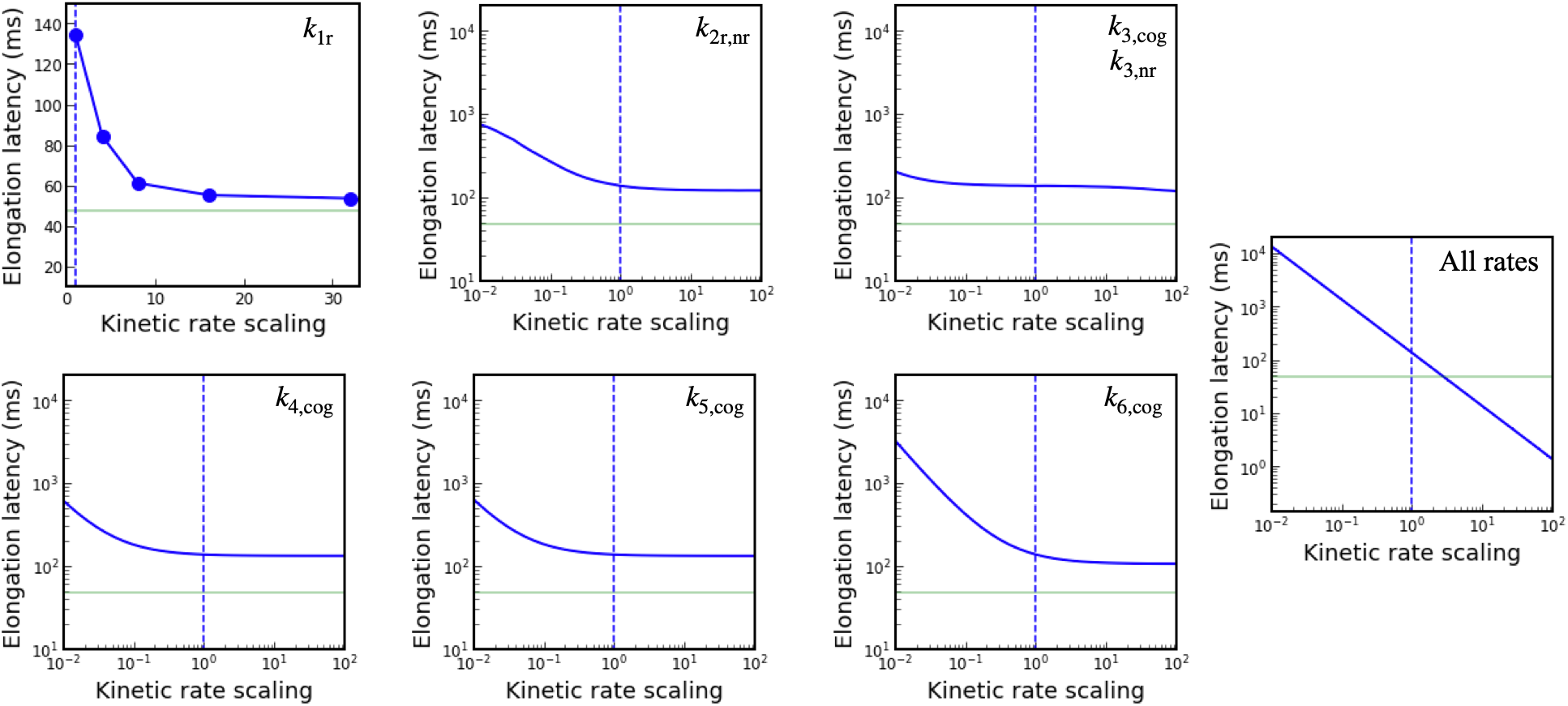
Our chemical kinetics sensitivity analysis shows that only changes in unbinding rate (*k*_1r_) or all kinetic rates together can speed up elongation to experimentally predicted latencies. Each plot shows the elongation latencies (solid blue line) that result from scaling one or more intra-ribosomal kinetic rates (specified in top right) to be faster or slower compared to baseline (dashed blue line). All simulations were for translation voxels at μ = 3.0 dbl/hr. Experimentally measured elongation latency at μ = 3.0 dbl/hr is shown for reference (green line). Scales are identical across plots except for the *k*_1r_ and All rates plots.

**Figure S10.**
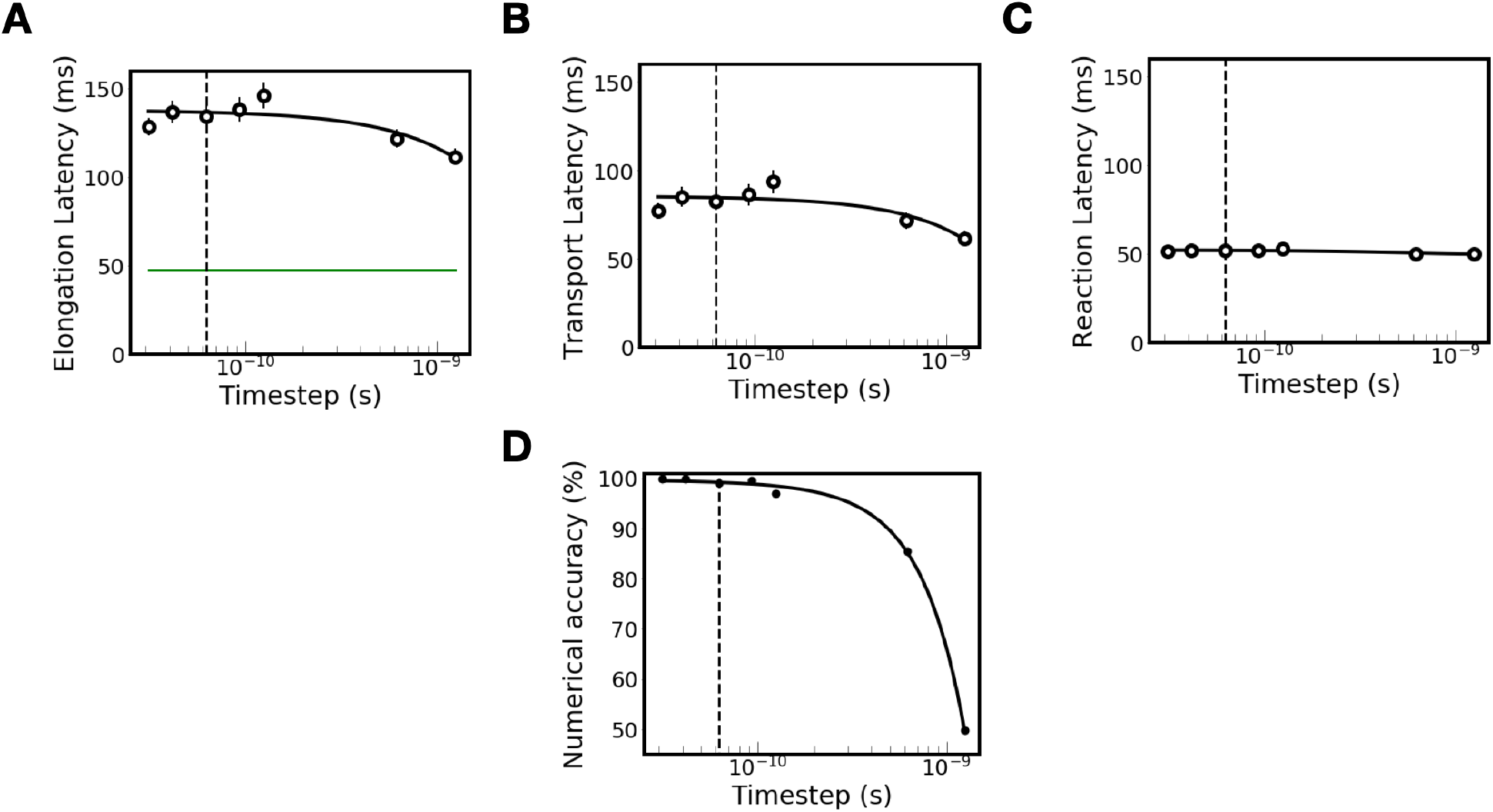
(A-C) Our simulation timestep sensitivity analysis shows that elongation latency, transport latency, and reaction latency each converge near our chosen timestep at the most crowded growth rate (μ = 3.0 dbl/hr). Experimentally measured elongation latency at μ = 3.0 dbl/hr is shown for reference in panel A (green line). **(D)** Numerical accuracy, computed as the percentage of entropic interactions between particles that are resolved, converges to 100% near our chosen timestep. All simulations were for translation voxels at μ = 3.0 dbl/hr. The dashed line corresponds to the timestep used for simulations throughout the paper (Δ*t* = 62 picoseconds).

**Figure S11.**
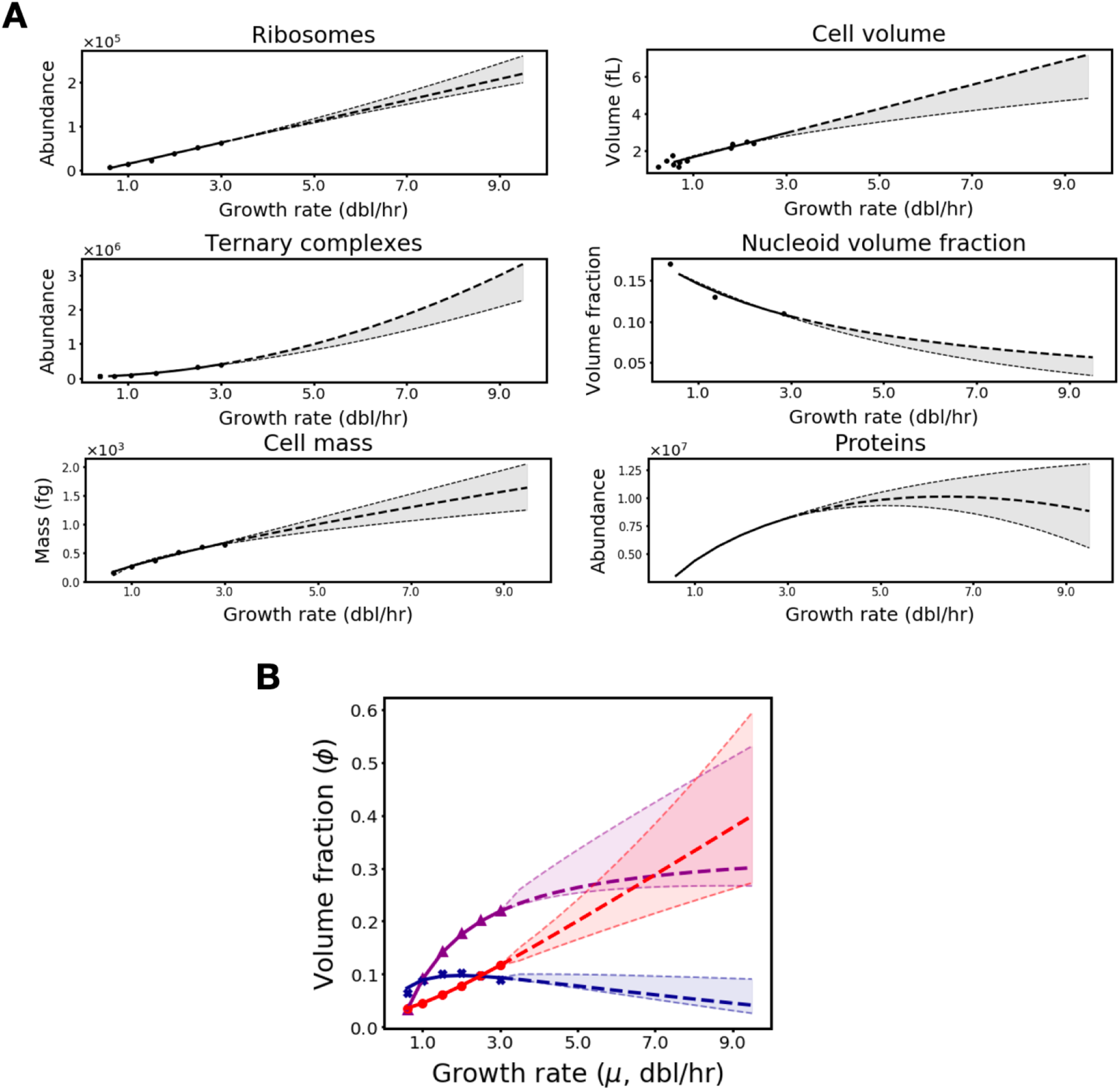
Cell and translation voxel parameters can be projected to faster-than-observed growth rates. **(A)** Extrapolated cell parameter fits (bolded dashed lines) projecting beyond observed growth rates (solid lines) for ribosomes (circle values from Table S1), ternary complexes (circle values from Table S1), cell mass (circle values from Table S1), cell volume (circle values from Table S2), nucleoid volume fraction (circle values from Table S3), and proteins. Perturbations to primary fits provide bounds (dashed lines and grey shaded region). **(B)** Extrapolated volume fractions (bolded dashed lines) projecting beyond observed growth rates (solid lines) for ribosomes (purple curve, triangles), ternary complexes (red curve, circles), and proteins (blue curve, x’s). Volume fractions are computed from cell parameter fits (**Methods**). Bounds for extrapolated volume fractions at hypothetical growth rates (light dashed lines and shading) are derived from permutations of the bounds for all cell parameters in panel A.

**Figure S12.**
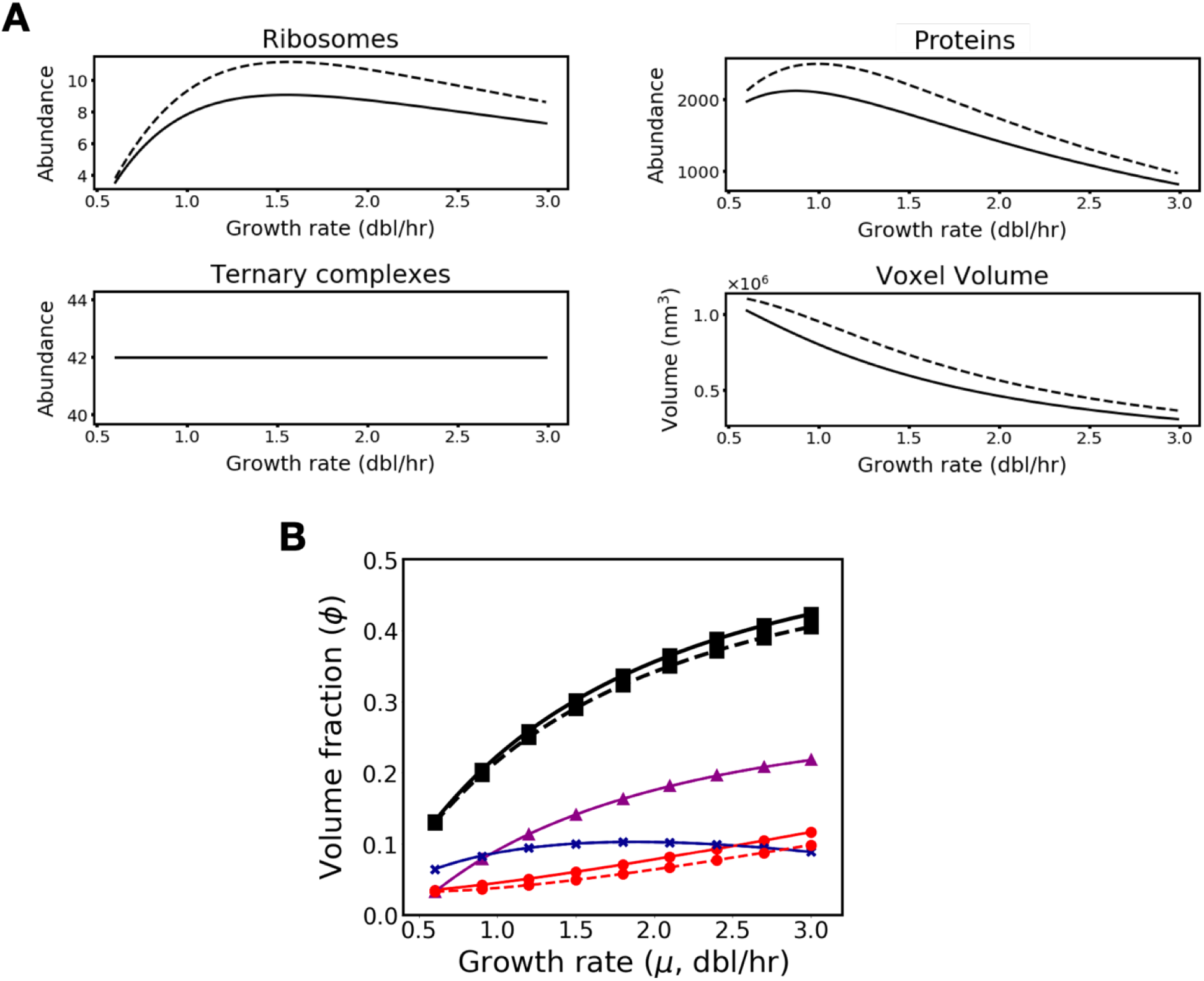
Whether or not literature-determined values for ternary complex abundances account for peptidyl-tRNA does not impact voxel composition or trends. **(A)** Key growth-rate dependent trends in translation voxel composition resulting from the modeling used throughout our manuscript (solid line), in which we interpret ternary complex abundance measurements as not including peptidyl-tRNA, are equivalent to those resulting from assuming that ternary complex abundance measurements do include peptidyl-tRNA (dashed line). **(B)** The volume fraction of ribosomes (purple curve, triangles), ternary complexes (red curve, circles), and proteins (blue curve, x’s), as well as total volume fraction (black curve, squares) are negligibly different for voxels constructed with the assumption that ternary complex measurements do not include peptidyl-tRNA (solid line) and the assumption that they do (dashed line).

**Figure S13.**
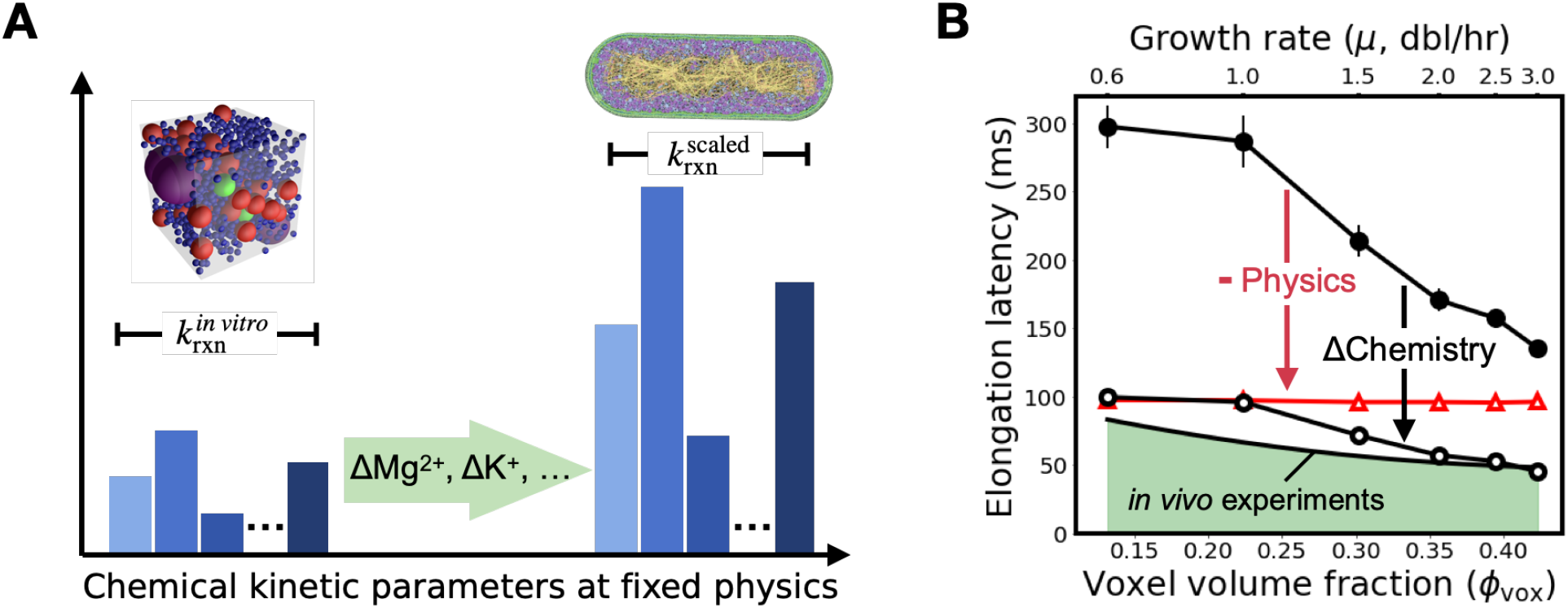
Faster elongation due to stoichiometric crowding is insensitive to changes in chemical kinetic parameters while sensitive to changes in transport physics. **(A)** Accuracy of chemical kinetics: cartoon schematic illustrating how differences in salt concentrations between *in vitro* conditions in which chemical kinetic rates were measured and *in vivo* conditions being modeled could lead to different and perhaps faster kinetic rates than used in our simulations. **(B)** Impact of faster chemistry, instantaneous physics: our simulations with three-fold faster kinetic rates produce elongation latency (open circles) that closes the quantitative gap between our original elongation latency prediction using published *in vitro* kinetic rates (solid circles) and *in vivo* measurements of per-ribosome elongation time (solid black line). The essentiality of physics in this agreement is demonstrated by the lack of faster elongation in simulations with instantaneous transport (red triangles). Depiction of *E. coli* in (A) adapted with permission from Goodsell, 2009.

**Figure S14.**
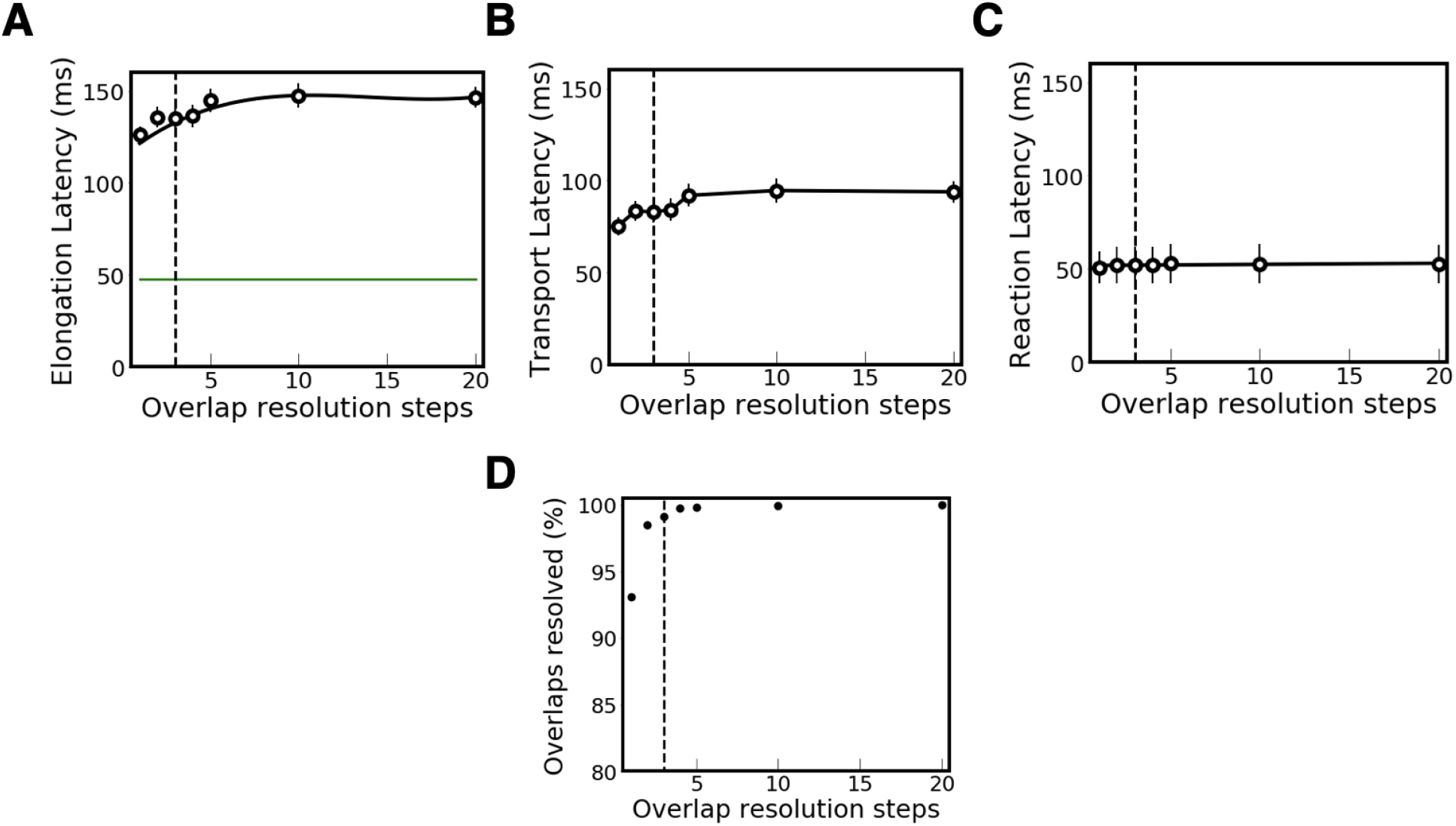
(A-C) Our simulation overlap resolution step analysis shows that elongation latency, transport latency, and reaction latency each converge near our chosen overlap resolution step at the most crowded growth rate (μ = 3.0 dbl/hr). Experimentally measured elongation latency at μ = 3.0 dbl/hr is shown for reference in panel A (green line). **(D)** Overlaps resolved under threshold error converge to near 100% near our chosen number of overlap resolution steps. All simulations were for translation voxels at μ = 3.0 dbl/hr. The dashed line corresponds to the overlap resolution steps used for simulations throughout the paper (three overlap resolution steps).

**Figure S15.**
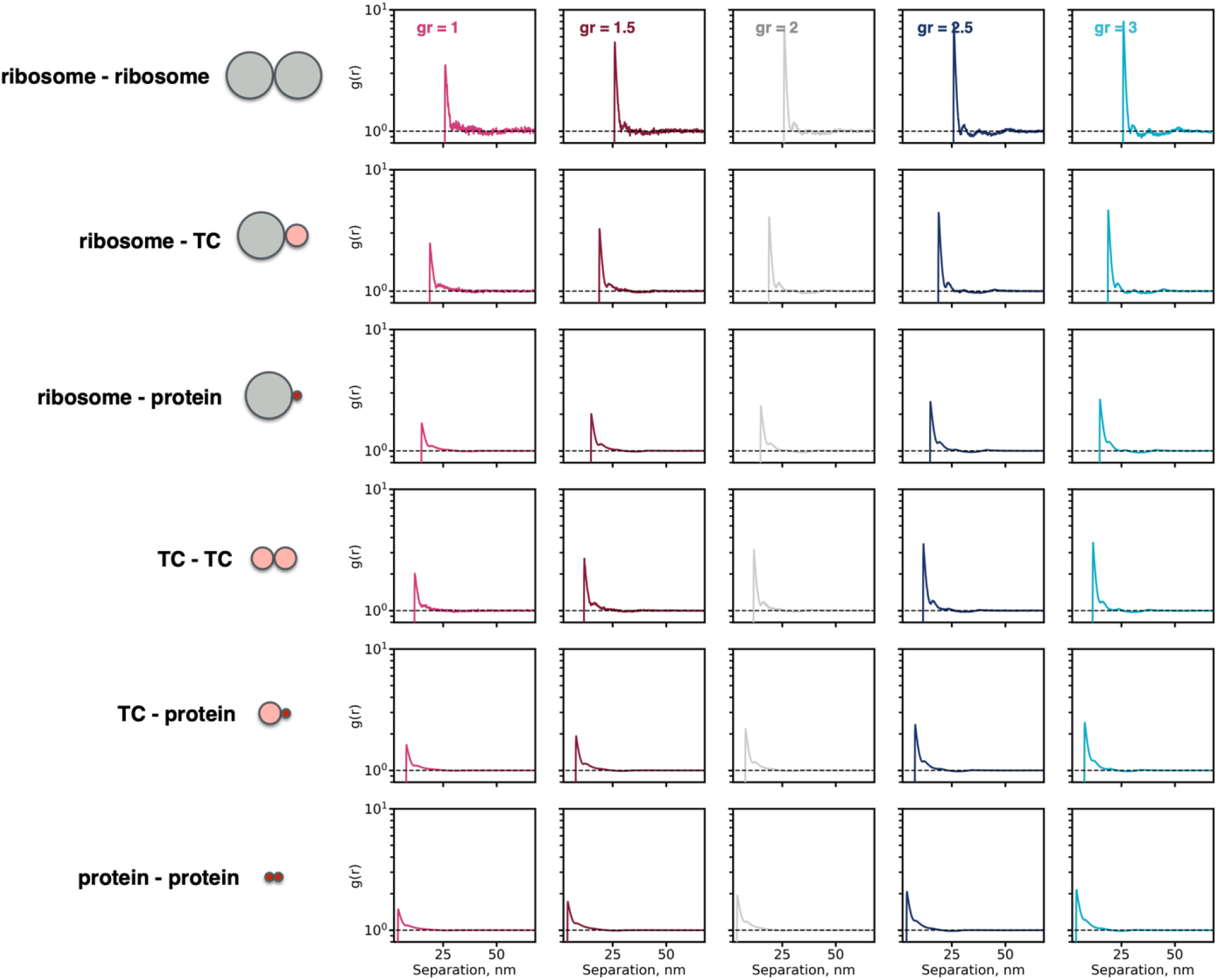
Translation voxels avoid finite size effects. The pair-correlation function *g*(*r*) between all possible combinations of pairing between ribosomes, ternary complexes (TC), and proteins (r is center-center separation) shows that all spatial correlations due to hard-sphere entropic exclusion decay at distances smaller than voxel sizes at all growth rates. Horizontal dashed line marks *g*(*r*) = 1. All simulations were performed using LAMMPS.

**Figure S16.**
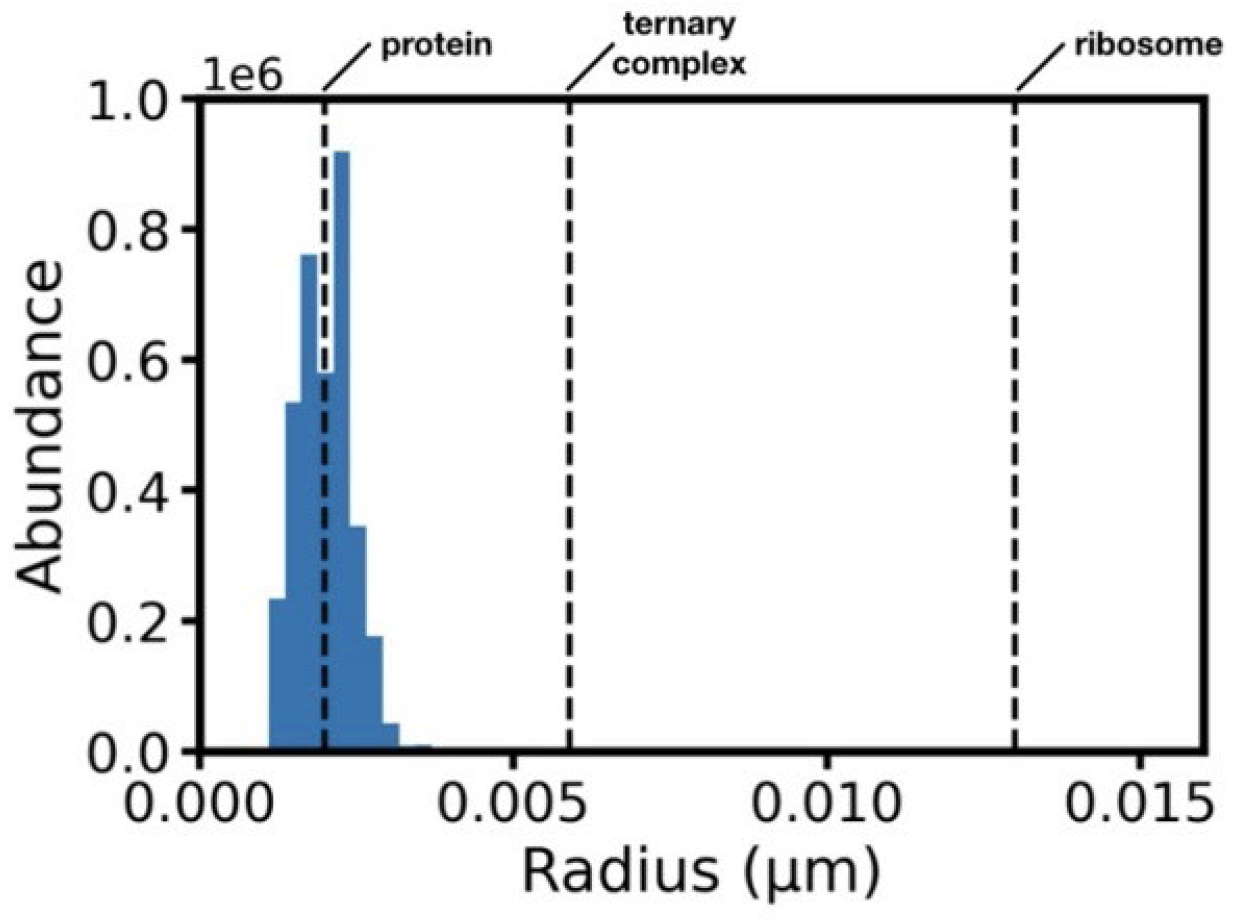
Size polydispersity of proteins in *E. coli*. Dashed lines show the size of an average-sized protein, a ternary complex, and a ribosome.

**Figure S17.**
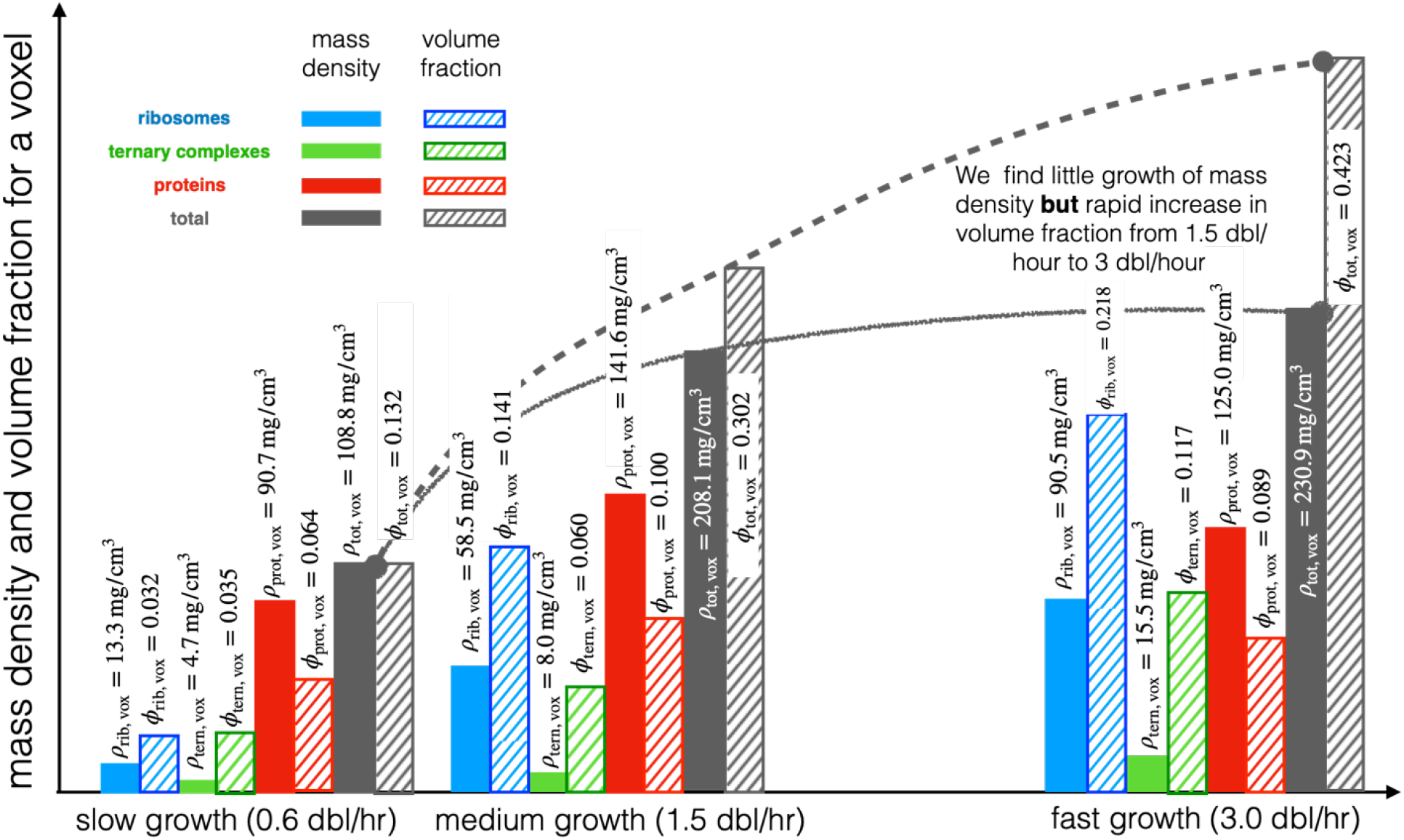
Plot of mass density (filled bars) and volume fraction (hashed bars) at three growth rates. Data shown on a per-molecule and voxel basis. Data is tabulated and cited in our Supplementary Tables 1–4. Changes in volume fraction do not imply a one-to-one change in mass density.

**Figure S18.**
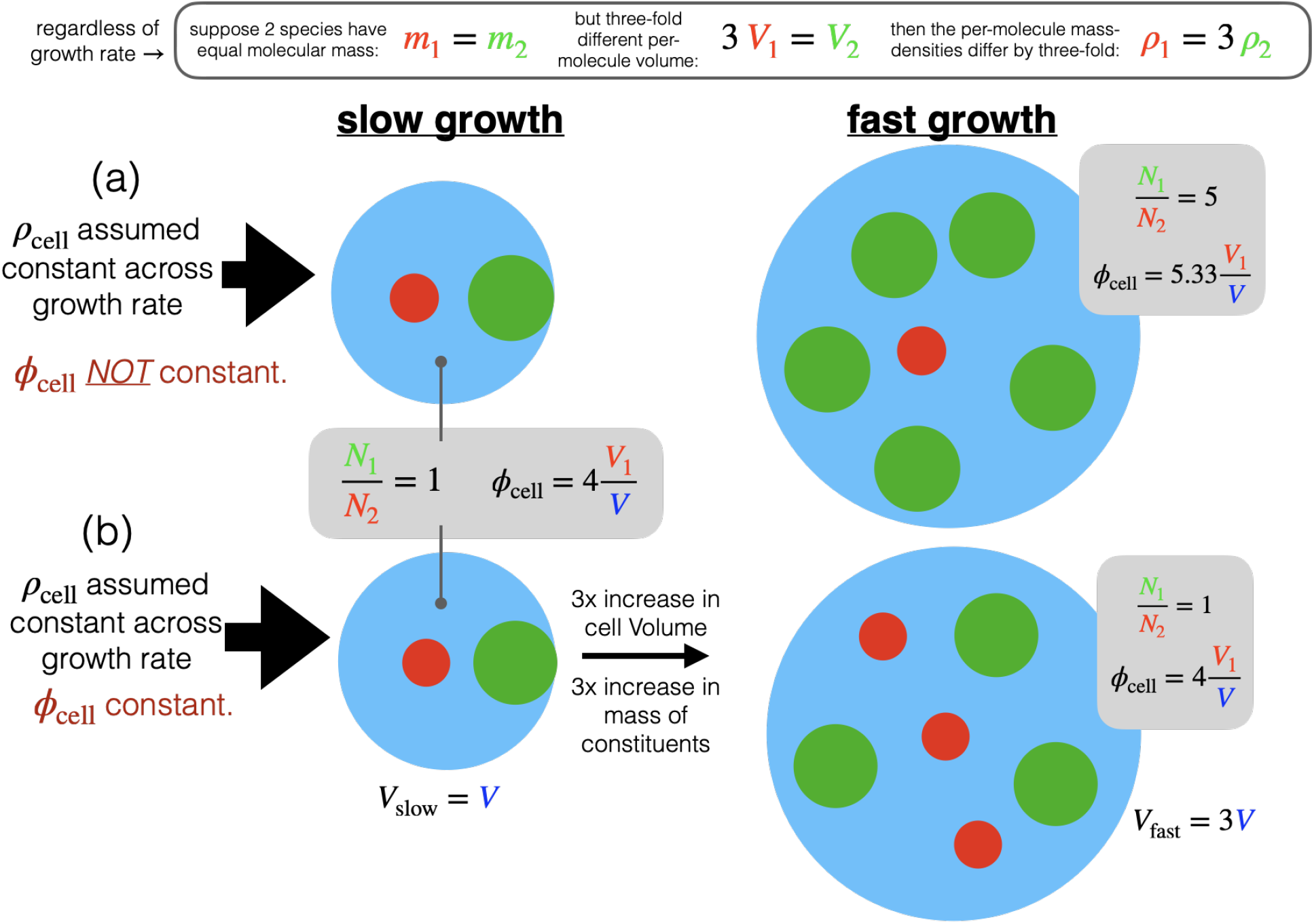
Mass density versus volume fraction. **(A)** Mass density of a cell can remain constant across growth rate (left to right) but volume fraction can grow. **(B)** Mass density and volume fraction can both remain constant but only if stoichiometry is held fixed. In *E. coli*, the stoichiometry (relative abundance of translation molecules) is well known to vary with growth rate and thus represents case (a).

**Table S1.**
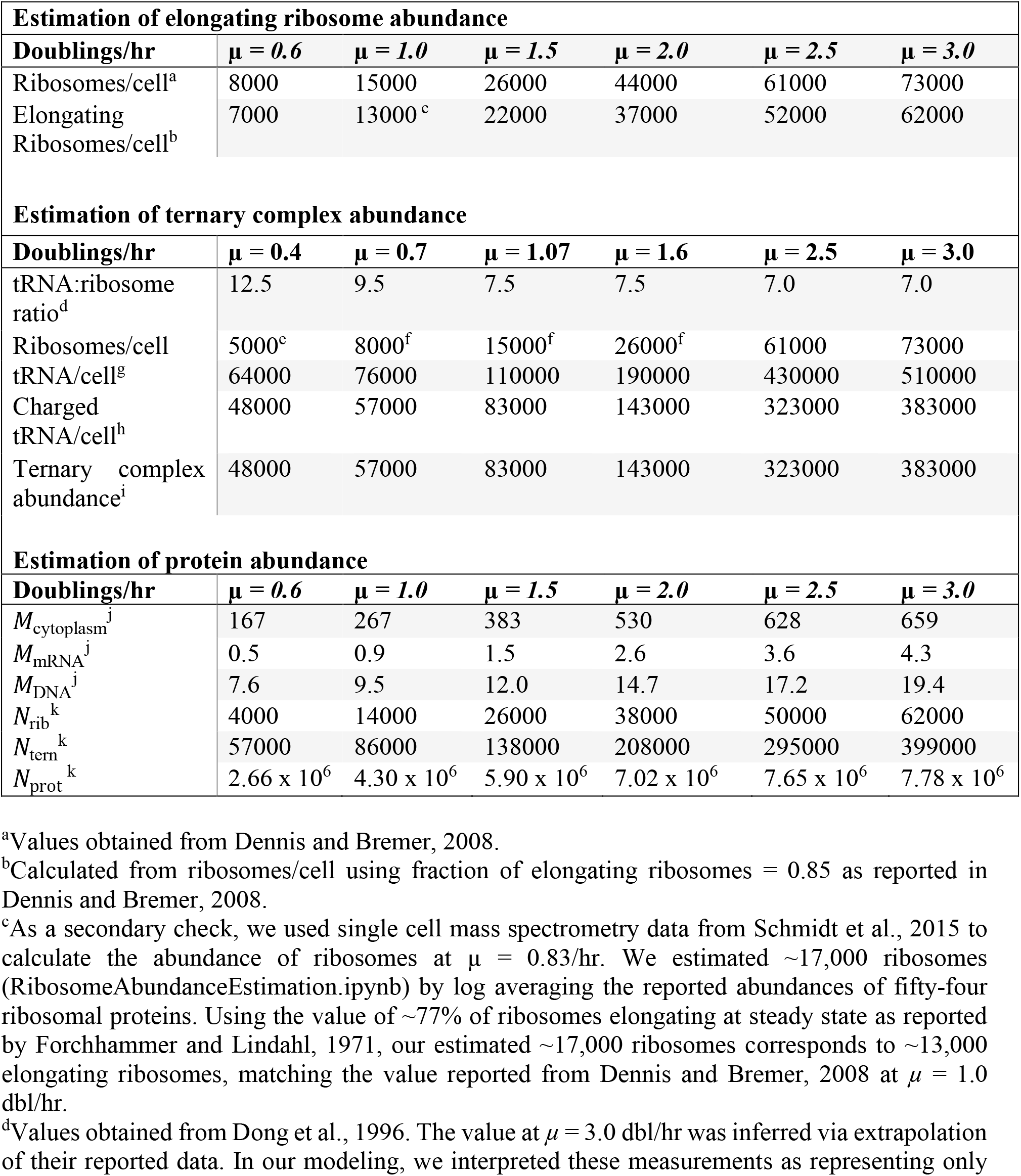
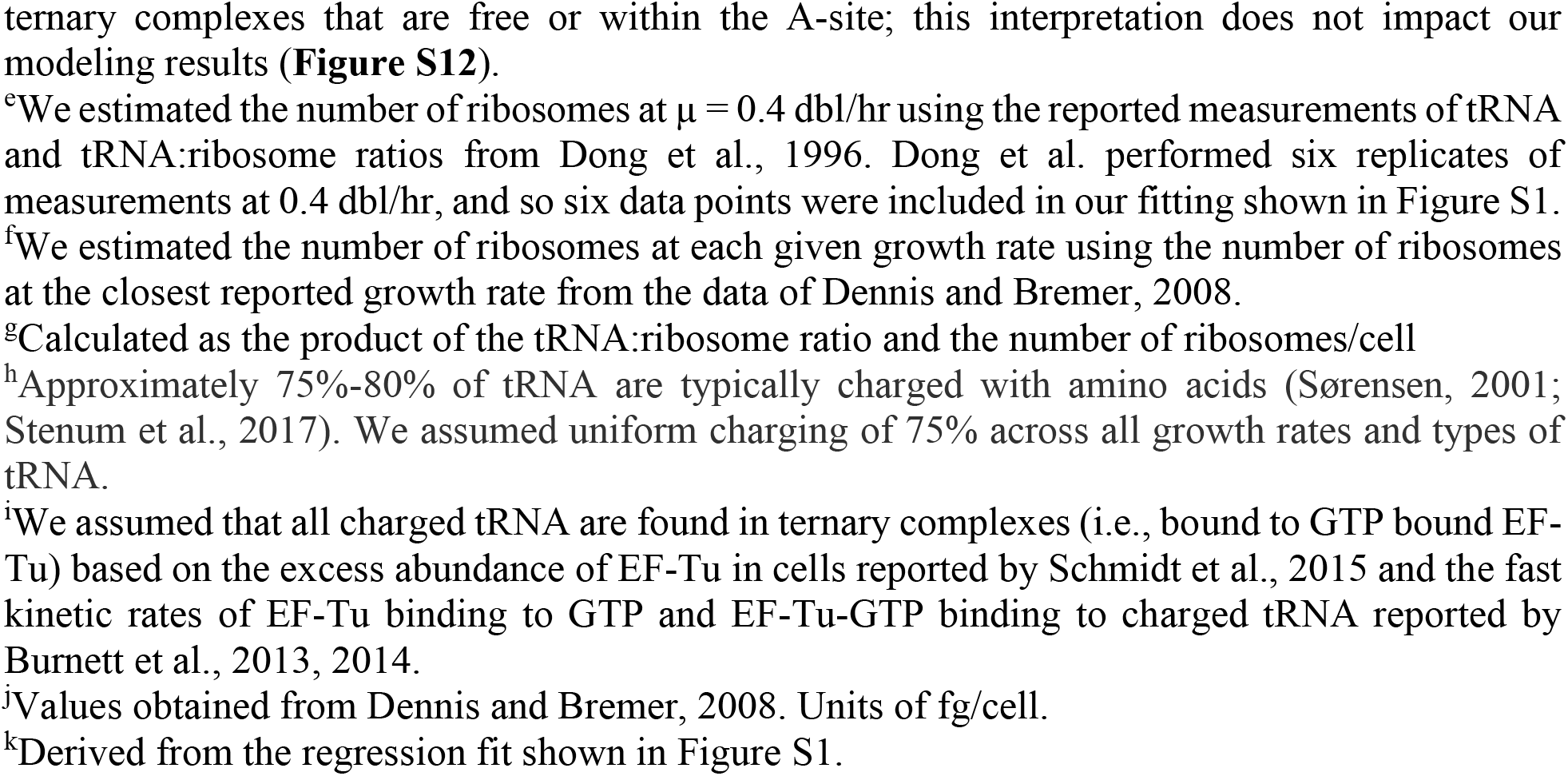
Estimation of elongating ribosome, ternary complex, and protein abundances at varying growth rates.

| **Estimation of elongating ribosome abundance** |  |  |  |  |  |  |
| --- | --- | --- | --- | --- | --- | --- |
| **Doublings/hr** | **$\mu = 0.6$** | **$\mu = 1.0$** | **$\mu = 1.5$** | **$\mu = 2.0$** | **$\mu = 2.5$** | **$\mu = 3.0$** |
| Ribosomes/cell<sup>a</sup> | 8000 | 15000 | 26000 | 44000 | 61000 | 73000 |
| Elongating Ribosomes/cell<sup>b</sup> | 7000 | 13000<sup>c</sup> | 22000 | 37000 | 52000 | 62000 |
| **Estimation of ternary complex abundance** |  |  |  |  |  |  |
| **Doublings/hr** | **$\mu = 0.4$** | **$\mu = 0.7$** | **$\mu = 1.07$** | **$\mu = 1.6$** | **$\mu = 2.5$** | **$\mu = 3.0$** |
| tRNA:ribosome ratio<sup>d</sup> | 12.5 | 9.5 | 7.5 | 7.5 | 7.0 | 7.0 |
| Ribosomes/cell | 5000<sup>e</sup> | 8000<sup>f</sup> | 15000<sup>f</sup> | 26000<sup>f</sup> | 61000 | 73000 |
| tRNA/cell<sup>g</sup> | 64000 | 76000 | 110000 | 190000 | 430000 | 510000 |
| Charged tRNA/cell<sup>h</sup> | 48000 | 57000 | 83000 | 143000 | 323000 | 383000 |
| Ternary complex abundance<sup>i</sup> | 48000 | 57000 | 83000 | 143000 | 323000 | 383000 |
| **Estimation of protein abundance** |  |  |  |  |  |  |
| **Doublings/hr** | **$\mu = 0.6$** | **$\mu = 1.0$** | **$\mu = 1.5$** | **$\mu = 2.0$** | **$\mu = 2.5$** | **$\mu = 3.0$** |
| $M_{\text{cytoplasm}}^j$ | 167 | 267 | 383 | 530 | 628 | 659 |
| $M_{\text{mRNA}}^j$ | 0.5 | 0.9 | 1.5 | 2.6 | 3.6 | 4.3 |
| $M_{\text{DNA}}^j$ | 7.6 | 9.5 | 12.0 | 14.7 | 17.2 | 19.4 |
| $N_{\text{rib}}^k$ | 4000 | 14000 | 26000 | 38000 | 50000 | 62000 |
| $N_{\text{tern}}^k$ | 57000 | 86000 | 138000 | 208000 | 295000 | 399000 |
| $N_{\text{prot}}^k$ | $2.66 \times 10^6$ | $4.30 \times 10^6$ | $5.90 \times 10^6$ | $7.02 \times 10^6$ | $7.65 \times 10^6$ | $7.78 \times 10^6$ |

<sup>a</sup>Values obtained from Dennis and Bremer, 2008.
<sup>b</sup>Calculated from ribosomes/cell using fraction of elongating ribosomes = 0.85 as reported in Dennis and Bremer, 2008.
<sup>c</sup>As a secondary check, we used single cell mass spectrometry data from Schmidt et al., 2015 to calculate the abundance of ribosomes at $\mu = 0.83/\text{hr}$ . We estimated ~17,000 ribosomes (RibosomeAbundanceEstimation.ipynb) by log averaging the reported abundances of fifty-four ribosomal proteins. Using the value of ~77% of ribosomes elongating at steady state as reported by Forchhammer and Lindahl, 1971, our estimated ~17,000 ribosomes corresponds to ~13,000 elongating ribosomes, matching the value reported from Dennis and Bremer, 2008 at $\mu = 1.0$ dbl/hr.
<sup>d</sup>Values obtained from Dong et al., 1996. The value at $\mu = 3.0$ dbl/hr was inferred via extrapolation of their reported data. In our modeling, we interpreted these measurements as representing only
ternary complexes that are free or within the A-site; this interpretation does not impact our modeling results (**Figure S12**).
<sup>e</sup>We estimated the number of ribosomes at $\mu = 0.4$ dbl/hr using the reported measurements of tRNA and tRNA:ribosome ratios from Dong et al., 1996. Dong et al. performed six replicates of measurements at 0.4 dbl/hr, and so six data points were included in our fitting shown in Figure S1.
<sup>f</sup>We estimated the number of ribosomes at each given growth rate using the number of ribosomes at the closest reported growth rate from the data of Dennis and Bremer, 2008.
<sup>g</sup>Calculated as the product of the tRNA:ribosome ratio and the number of ribosomes/cell
<sup>h</sup>Approximately 75%-80% of tRNA are typically charged with amino acids (Sørensen, 2001; Stenum et al., 2017). We assumed uniform charging of 75% across all growth rates and types of tRNA.
<sup>i</sup>We assumed that all charged tRNA are found in ternary complexes (i.e., bound to GTP bound EF-Tu) based on the excess abundance of EF-Tu in cells reported by Schmidt et al., 2015 and the fast kinetic rates of EF-Tu binding to GTP and EF-Tu-GTP binding to charged tRNA reported by Burnett et al., 2013, 2014.
<sup>j</sup>Values obtained from Dennis and Bremer, 2008. Units of fg/cell.
<sup>k</sup>Derived from the regression fit shown in Figure S1.

**Table S2.**
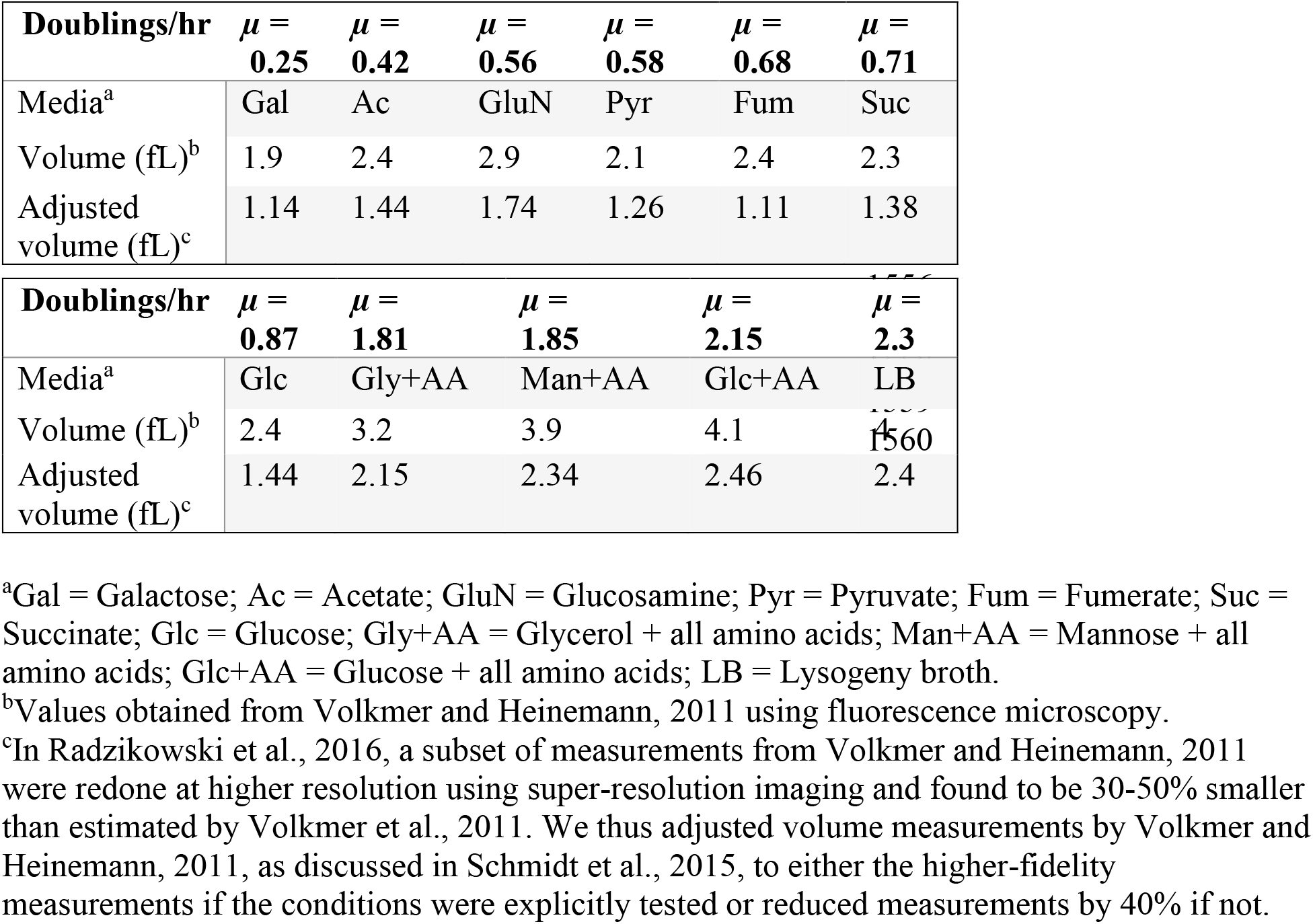
Estimation of cell volume at varying growth rates.

**Table S3.**
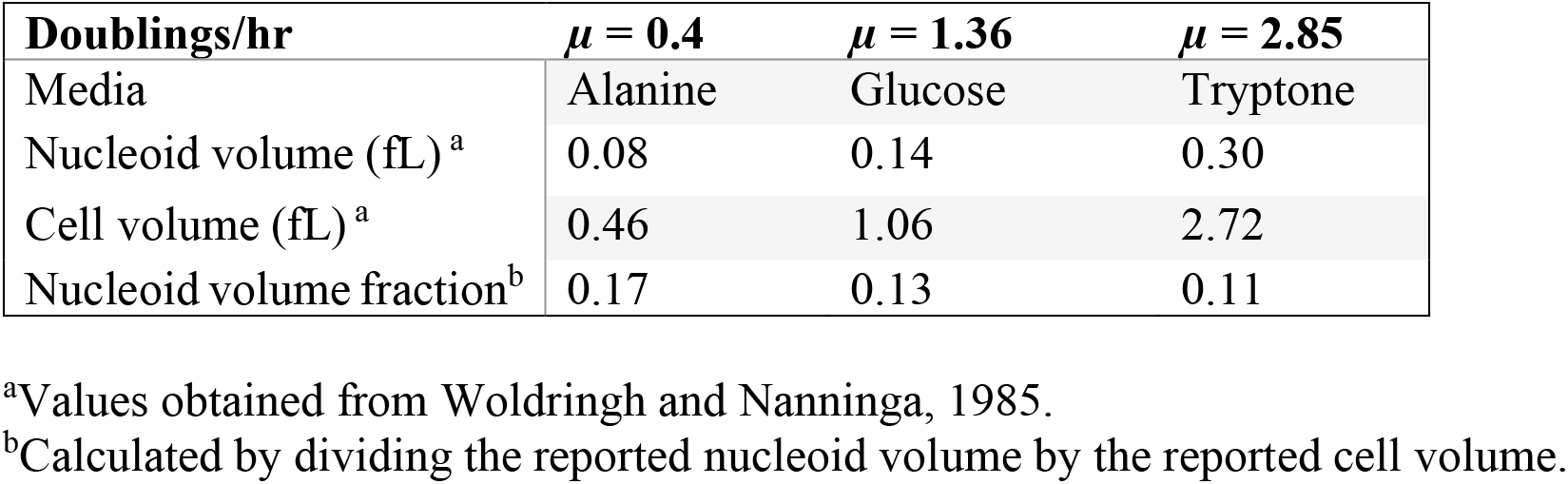
Estimation of nucleoid volume fraction at varying growth rates.

| **Doublings/hr** | **$\mu = 0.4$** | **$\mu = 1.36$** | **$\mu = 2.85$** |
| --- | --- | --- | --- |
| Media | Alanine | Glucose | Tryptone |
| Nucleoid volume (fL)<sup>a</sup> | 0.08 | 0.14 | 0.30 |
| Cell volume (fL)<sup>a</sup> | 0.46 | 1.06 | 2.72 |
| Nucleoid volume fraction<sup>b</sup> | 0.17 | 0.13 | 0.11 |

<sup>a</sup>Values obtained from Woldringh and Nanninga, 1985.
<sup>b</sup>Calculated by dividing the reported nucleoid volume by the reported cell volume.

**Table S4.** Parameters not varying with growth rate.

|  |  |
| --- | --- |
| $M_{\text{rib}}$ | 2300 kDa |
| $M_{\text{tern}}$ | 69 kDa |
| $M_{\text{tRNA}}$ | 25 kDa |
| $M_{\text{amino acid}}$ | 110 Da |
| $M_{\text{EF-Tu}}$ | 43.2 kDa |
| $M_{\text{GTP}}$ | 523 Da |
| $M_{\text{prot}}^{\text{a}}$ | 28.5 kDa |
| $\bar{V}_{\text{prot}}^{\text{b}}$ | $3.35 \times 10^{-8} \mu\text{m}^3$ |
| $\rho_{\text{prot}}^{\text{c}}$ | $1.410 \text{ g/cm}^3$ |
| $R_{\text{prot}}^{\text{d}}$ | 2.0 nm |
| $R_{\text{rib}}^{\text{de}}$ | 13 nm |
| $R_{\text{tern}}^{\text{df}}$ | 5.9 nm |
| $k_B$ | $1.380649 \times 10^{-23} \text{ J/K}$ |
| $T$ | 37 C |
| $\eta$ | 0.6913 mPa*s |
| $D_{\text{tern}}$ | $56 \mu\text{m}^2/\text{s}$ |
| $D_{\text{rib}}$ | $25 \mu\text{m}^2/\text{s}$ |
| $D_{\text{prot}}$ | $165 \mu\text{m}^2/\text{s}$ |

<sup>a</sup>We computed the mass of individual average-sized proteins as described in the methods. Given an average amino acid mass of 108 Da, as reported by Bremer and Dennis, 2008, the protein mass corresponds to 264 amino acids on average. This protein length corresponds well to estimates in the literature of the average and median number of amino acids for a protein, 309 and 252 amino acids respectively (Ishihama et al., 2008), supporting our calculations.
<sup>b</sup>The volume of an average-sized protein was calculated as described in the methods.
<sup>c</sup>Value obtained from Fischer et al., 2004.
<sup>d</sup>Biomolecule radius was calculated as described in the methods. Note that we use $R$ and $a$ interchangeably.
<sup>e</sup>Measured using PDB 4V4Q from Schuwirth et al., 2005 as described in the methods.
<sup>f</sup>Measured using PDB 1B23 from Nissen et al., 1999 as described in the methods.

**Table S5.**
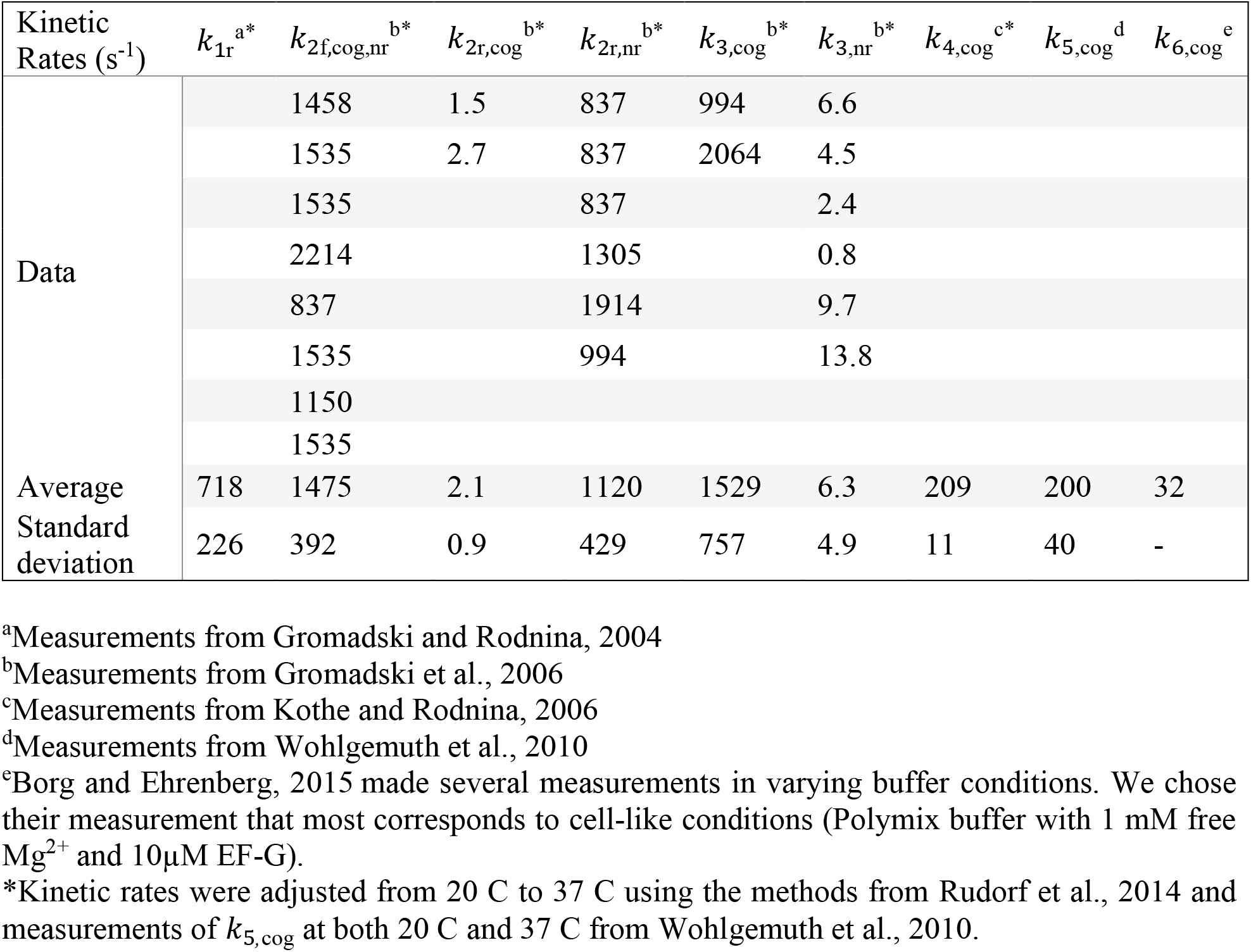
tRNA-ribosome reaction kinetic parameters for non-cognate, near-cognate, and cognate tRNA.

**Table S6.**
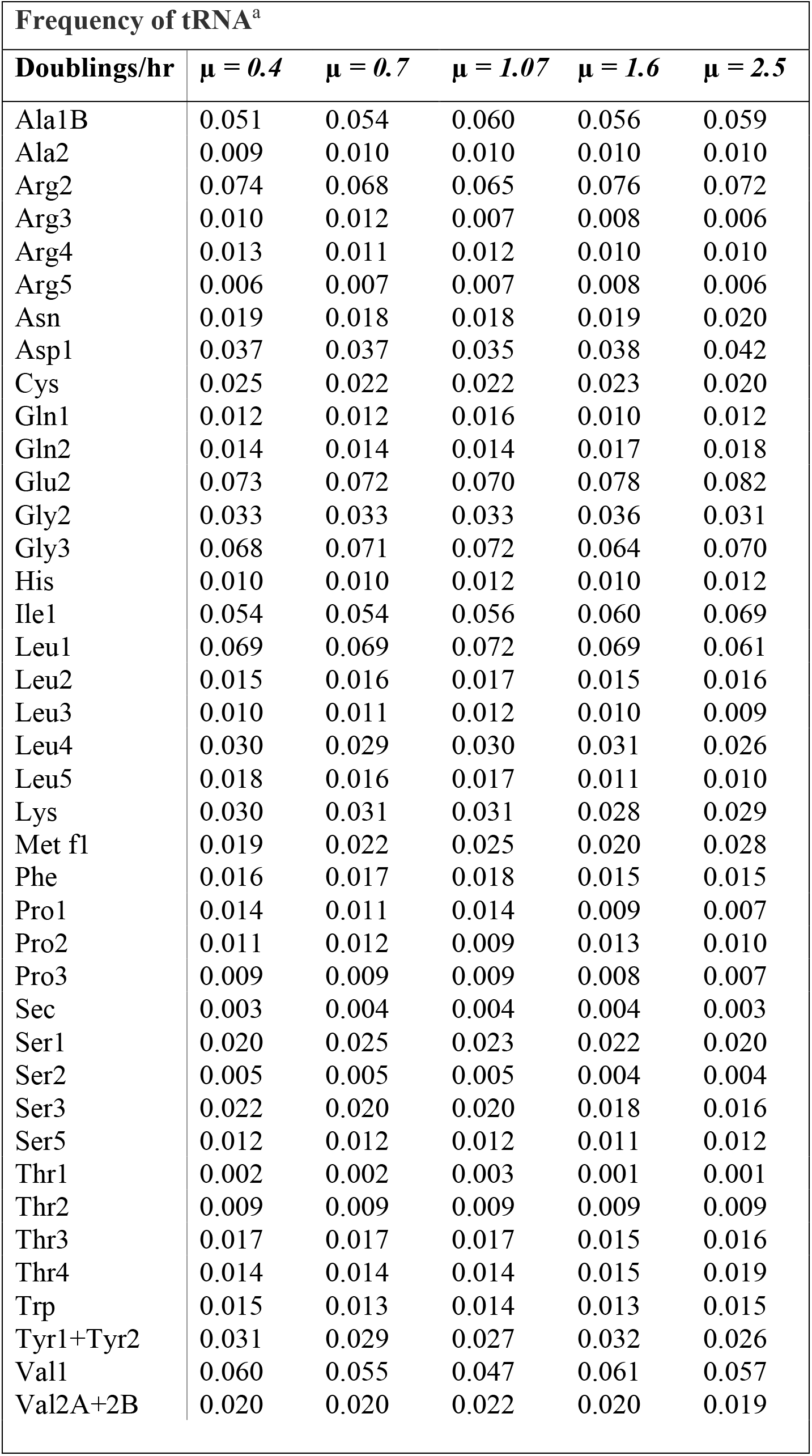
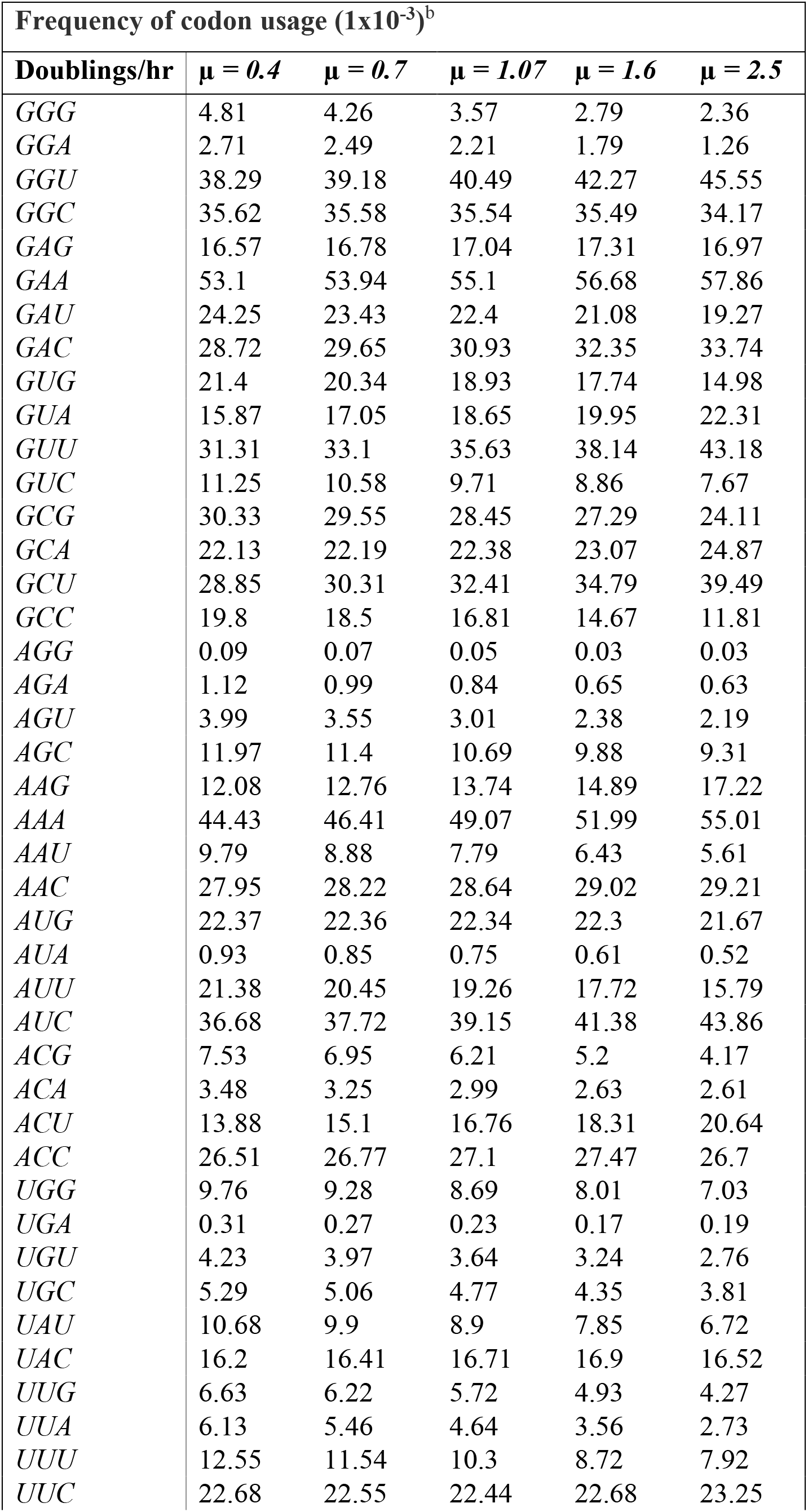
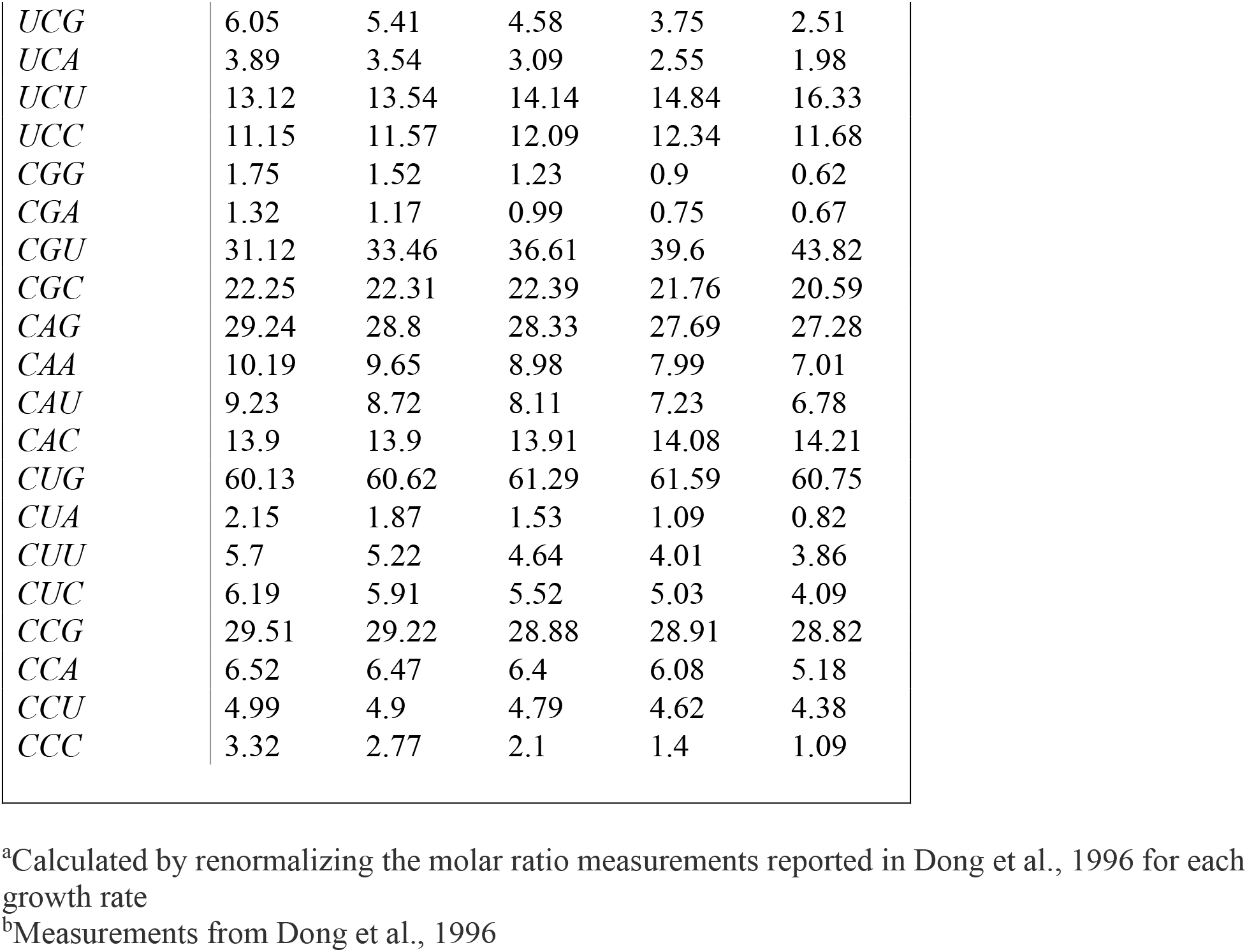
Frequency of tRNA and codon usage at varying growth rates.

## References

Andrews, S.S., and Bray, D. (2004). Stochastic simulation of chemical reactions with spatial resolution and single molecule detail. Phys. Biol. 1, 137–151.

Andrews, S.S., Addy, N.J., Brent, R., and Arkin, A.P. (2010). Detailed simulations of cell biology with Smoldyn 2.1. PLoS Comput. Biol. 6.

Aponte-Rivera, C., Su, Y., and Zia, R.N. (2018). Equilibrium structure and diffusion in concentrated hydrodynamically interacting suspensions confined by a spherical cavity. J. Fluid Mech. 836, 413–450.

Avcilar-Kucukgoze, I., Bartholomäus, A., Cordero Varela, J.A., Kaml, R.F.X., Neubauer, P., Budisa, N., and Ignatova, Z. (2016). Discharging tRNAs: A tug of war between translation and detoxification in Escherichia coli. Nucleic Acids Res. 44, 8324–8334.

Banchio, A.J., and Brady, J.F. (2003). Accelerated stokesian dynamics: Brownian motion. J. Chem. Phys.

Batchelor, G.K. (1977). The effect of Brownian motion on the bulk stress in a suspension of spherical particles. J. Fluid Mech.

Borg, A., and Ehrenberg, M. (2015). Determinants of the rate of mRNA translocation in bacterial protein synthesis. J. Mol. Biol. 427, 1835–1847.

Brady, J.F. (1993). Brownian motion, hydrodynamics, and the osmotic pressure. J. Chem. Phys.

Bremer, H., and Dennis, P. (1996). Modulation of chemical composition and other parameters of the cell by growth rate. In Escherichia Coli and Salmonella: Cellular and Molecular Biology, F. Neidthard, ed. (Washington, DC: ASM Press), pp. 1553–1569.

Burnett, B.J., Altman, R.B., Ferrao, R., Alejo, J.L., Kaur, N., Kanji, J., and Blanchard, S.C. (2013). Elongation factor Ts directly facilitates the formation and disassembly of the escherichia coli elongation factor Tu·GTP·aminoacyl-tRNA ternary complex. J. Biol. Chem. 288, 13917–13928.

Burnett, B.J., Altman, R.B., Ferguson, A., Wasserman, M.R., Zhou, Z., and Blanchard, S.C. (2014). Direct evidence of an elongation factor-Tu/Ts · GTP · aminoacyl-tRNA quaternary complex. J. Biol. Chem. 289, 23917–23927.

Dalbow, D.G., and Young, R. (1975). Synthesis time of β galactosidase in Escherichia coli B/r as a function of growth rate. Biochem. J. 150, 13–20.

Dennis, P.P., and Bremer, H. (2008). Modulation of Chemical Composition and Other Parameters of the Cell at Different Exponential Growth Rates. EcoSal Plus 3.

Dill, K.A., Ghosh, K., and Schmit, J.D. (2011). Physical limits of cells and proteomes. Proc. Natl. Acad. Sci. U. S. A. 108, 17876–17882.

Dong, H., Nilsson, L., and Kurland, C.G. (1996). Co-variation of tRNA Abundance and Codon Usage in Escherichia coli at Different Growth Rates. J. Mol. Biol. 260, 649–663.

Durlofsky, L., Brady, J.F., and Bossis, G. (1987). Dynamic Simulation of Hydrodynamically Interacting Particles. J. Fluid Mech.

Einstein, A. (1905). On the Motion of Small Particles Suspended in a Stationary Liquid, as Required by the Molecular Kinetic Theory of Heat. Ann. Phys. 322, 549–560.

Endy, D., and Brent, R. (2001). Modeling cellular behavior. Nature 409, 391–395.

Ermak, D.L., and McCammon, J.A. (1977). Brownian dynamics with hydrodynamic interactions. J. Chem. Phys. 66, 12–521.

Fairhurst, D.J. (1999). Polydispersity in Colloidal Phase Transitions. University of Edinburgh.

Farr, R.S., and Groot, R.D. (2009). Close packing density of polydisperse hard spheres. J. Chem. Phys. 131.

Farris, R.J. (1968). Prediction of the Viscosity of Multimodal Suspensions from Unimodal Viscosity Data. Trans. Soc. Rheol. 12, 281–301.

Fischer, H., Polikarpov, I., and Craievich, A.F. (2004). Average protein density is a molecular-weight-dependent function. Protein Sci. 13, 2825–2828.

Forchhammer, J., and Lindahl, L. (1971). Growth rate of polypeptide chains as a function of the cell growth rate in a mutant of Escherichia coli 15. J. Mol. Biol. 55, 563–568.

Foss, D.R., and Brady, J.F. (2000). Brownian Dynamics simulation of hard-sphere colloidal dispersions. J. Rheol. (N. Y. N. Y). 44, 629–651.

Gonzalez, E., Aponte-Rivera, C., and Zia, R.N. (2021). Impact of polydispersity and confinement on diffusion in hydrodynamically interacting colloidal suspensions. J. Fluid Mech. In press.

Goodsell, D.S. (2009). The Machinery of Life (Springer Science & Business Media).

Gromadski, K.B., and Rodnina, M. V. (2004). Kinetic Determinants of High-Fidelity tRNA Discrimination on the Ribosome. Mol. Cell 13, 191–200.

Gromadski, K.B., Daviter, T., and Rodnina, M. V. (2006). A uniform response to mismatches in codon-anticodon complexes ensures ribosomal fidelity. Mol. Cell 21, 369–377.

Grosjean, H., and Chantrenne, H. (1980). On Codon-Anticodon Interactions.

Heyes, D.M., and Melrose, J.R. (1993). Brownian dynamics simulations of model hard-sphere suspensions. 46, 1–28.

Hoh, N.J., and Zia, R.N. (2016a). Force-induced diffusion in suspensions of hydrodynamically interacting colloids. J. Fluid Mech.

Hoh, N.J., and Zia, R.N. (2016b). The impact of probe size on measurements of diffusion in active microrheology. Lab Chip.

Ishihama, Y., Schmidt, T., Rappsilber, J., Mann, M., Harlt, F.U., Kerner, M.J., and Frishman, D. (2008). Protein abundance profiling of the Escherichia coli cytosol. BMC Genomics 9, 1–17.

Kinz-Thompson, C.D., Bailey, N.A., and Gonzalez, R.L. (2016). Precisely and Accurately Inferring Single-Molecule Rate Constants. Methods Enzymol. 581, 187–225.

Klumpp, S., Scott, M., Pedersen, S., and Hwa, T. (2013). Molecular crowding limits translation and cell growth. Proc. Natl. Acad. Sci.

Kothe, U., and Rodnina, M. V. (2006). Delayed release of inorganic phosphate from elongation factor Tu following GTP hydrolysis on the ribosome. Biochemistry 45, 12767–12774.

Kothe, U., Wieden, H.J., Mohr, D., and Rodnina, M. V. (2004). Interaction of Helix D of Elongation Factor Tu with Helices 4 and 5 of Protein L7/12 on the Ribosome. J. Mol. Biol. 336, 1011–1021.

Langevin, P. (1908). Sur la theorie du mouvement brownien. C.R. Acad. Sci., 146.

Lionberger, R.A. (2002). Viscosity of bimodal and polydisperse colloidal suspensions. Phys. Rev. E - Stat. Physics, Plasmas, Fluids, Relat. Interdiscip. Top. 65.

Maheshwari, A.J., Sunol, A.M., Gonzalez, E., Endy, D., and Zia, R.N. (2019). Colloidal hydrodynamics of biological cells: A frontier spanning two fields. Phys. Rev. Fluids 4, 1–26.

Mustafi, M., and Weisshaar, J.C. (2018). Simultaneous binding of multiple EF-Tu copies to translating ribosomes in live Escherichia coli. MBio 9, 1–16.

Nissen, P., Thirup, S., Kjeldgaard, M., and Nyborg, J. (1999). The crystal structure of Cys-tRNA(Cys)-EF-Tu-GDPNP reveals general and specific features in the ternary complex and in tRNA. Structure 7, 143–156.

Oldewurtel, E.R., Kitahara, Y., and van Teeffelen, S. (2021). Robust surface-to-mass coupling and turgor-dependent cell width determine bacterial dry-mass density. Proc. Natl. Acad. Sci. U. S. A. 118.

Ouaknin, G.Y., Su, Y., and Zia, R.N. (2021). Simulation of large-scale particle systems at low Reynolds number: Parallel algorithms for Accelerated Stokesian Dynamics. J. Comput. Phys. In Review.

Pedersen, S. (1984). Escherichia coli ribosomes translate in vivo with variable rate. EMBO J. 3, 2895–2898.

Pedersen, S., Bloch, P., Reeh, S., and Neidhardt, F. (1978). Patterns of protein synthesis in E. coli: a catalog of the amount of 140 individual proteins at different growth rates. Cell 14, 179–190.

Plimpton, S. (1995). Fast parallel algorithms for short-range molecular dynamics. J. Comput. Phys. 117, 1–19.

Radzikowski, J.L., Vedelaar, S., Siegel, D., Ortega, Á.D., Schmidt, A., and Heinemann, M. (2016). Bacterial persistence is an active σ^S^ stress response to metabolic flux limitation. Mol. Syst. Biol. 12, 882.

Rudorf, S., Thommen, M., Rodnina, M. V., and Lipowsky, R. (2014). Deducing the Kinetics of Protein Synthesis In Vivo from the Transition Rates Measured In Vitro. PLoS Comput. Biol. 10.

Russel, W.B. (1984). The Huggins coefficient as a means for characterizing suspended particles. J. Chem. Soc. Faraday Trans. 2 Mol. Chem. Phys.

Russel, W.B., Saville, D.A., and Schowalter, W.R. (1989). Colloidal Dispersions (Cambridge University Press).

Sannuga, S., and Ramakrishnan, V. (2004). The Ribosome in Protein Synthesis.

Schmidt, A., Kochanowski, K., Vedelaar, S., Ahrne, E., Volkmer, B., Callipo, L., Knoops, K., Bauer, M., Aebersold, R., and Heinemann, M. (2015). The quantitative and condition-dependent Escherichia coli proteome. Nat. Biotechnol. 34, 104–110.

Schuwirth, B.S., Borovinskaya, M.A., Hau, C.W., Zhang, W., Vila-Sanjurjo, A., Holton, J.M., and Cate, J.H.D. (2005). Structures of the bacterial ribosome at 3.5 Å resolution. Science (80-.). 310, 827–834.

Sierou, A., and Brady, J.F. (2001). Accelerated Stokesian Dynamics simulations. J. Fluid Mech. 448, 115–146.

Sørensen, M.A. (2001). Charging levels of four tRNA species in Escherichia coli Rel+and Rel-strains during amino acid starvation: A simple model for the effect of ppGpp on translational accuracy. J. Mol. Biol. 307, 785–798.

Stenum, T.S., Sørensen, M.A., and Svenningsen, S. Lo (2017). Quantification of the Abundance and Charging Levels of Transfer RNAs in Escherichia coli. J. Vis. Exp. 5621237915, 1–10.

Stokes, G.G. (1850). On the effect of the Internal friction of fluids on the motion of pendulums - Section III. Trans. Cambridge Philos. Soc.

Subramaniam, A.R., Zid, B.M., and O’Shea, E.K. (2014). An integrated approach reveals regulatory controls on bacterial translation elongation. Cell 159, 1200–1211.

Takahashi, K., Tǎnase-Nicola, S., and Ten Wolde, P.R. (2010). Spatio-temporal correlations can drastically change the response of a MAPK pathway. Proc. Natl. Acad. Sci. U. S. A. 107, 2473–2478.

Vieira, J.P., Racle, J., and Hatzimanikatis, V. (2016). Analysis of Translation Elongation Dynamics in the Context of an Escherichia coli Cell. Biophys. J. 110, 2120–2131.

Volkmer, B., and Heinemann, M. (2011). Condition-Dependent cell volume and concentration of Escherichia coli to facilitate data conversion for systems biology modeling. PLoS One 6, 1–6.

Welch, M., Govindarajan, S., Ness, J.E., Villalobos, A., Gurney, A., Minshull, J., and Gustafsson, C. (2009). Design parameters to control synthetic gene expression in Eschorichia coli. PLoS One 4.

Wohlgemuth, I., Pohl, C., and Rodnina, M. V. (2010). Optimization of speed and accuracy of decoding in translation. EMBO J. 29, 3701–3709.

Woldringh, C.L., and Nanninga, N. (1985). Structure of nucleoid and cytoplasm of the intact cell. In Molecular Cytology of Escherichia Coli, (London: Academic Press), pp. 161–197.

Young, R., and Bremer, H. (1976). Polypeptide-chain-elongation rate in Escherichia coli B/r as a function of growth rate. Biochem. J. 160, 185–194.

Zakhari, M.E.A., Anderson, P.D., and Hütter, M. (2017). Effect of particle-size dynamics on properties of dense spongy-particle systems: Approach towards equilibrium. Phys. Rev. E 96, 1–16.

Zia, R.N. (2018). Active and Passive Microrheology: Theory and Simulation. Annu. Rev. Fluid Mech. 50, 371–405.

Zia, R.N., and Brady, J.F. (2010). Single-particle motion in colloids: force-induced diffusion. J. Fluid Mech. 658, 188–210.

Zia, R.N., and Brady, J.F. (2012). Microviscosity, microdiffusivity, and normal stresses in colloidal dispersions. J. Rheol. (N. Y. N. Y). 56, 1175–1208.

